# Quantitative measurement of synthetic repression curves reveals design challenges for genetic circuit engineering under growth arrest

**DOI:** 10.64898/2026.02.01.703179

**Authors:** John P. Marken, Mark L. Prator, Bruce A. Hay, Richard M. Murray

**Affiliations:** Division of Biology and Biological Engineering, California Institute of Technology, 1200 E California Blvd, Pasadena, CA, USA 91125; Resnick Sustainability Institute, California Institute of Technology, 1200 E California Blvd, Pasadena, CA, USA 91125; Section of Microbiology, University of Copenhagen, Universitetsparken 15, DK-2100, Copenhagen, Denmark

**Keywords:** Synthetic biology, genetic circuits, growth arrest, microbial physiology, gene regulation

## Abstract

Despite the fact that microbes in natural environments spend most of their time in growth arrest, we understand little about how this physiological state affects the performance of engineered genetic circuits. Here, we measure repression curves from a library of genetic NOT gates at single-cell resolution in *Escherichia coli* under both active growth and growth arrest to systematically investigate how growth arrest affects circuit behavior. We find that the impact of growth arrest on circuit performance is almost entirely dominated by a single effect: a >100-fold reduction in unrepressed expression levels. Growth arrest caused gene expression noise to increase moderately and had only minimal impacts on the sensitivity and sharpness of the repression curves. Our work shows both that conventional genetic circuit design paradigms are currently insufficient to develop circuits that can function properly under growth arrest, but also that addressing the reduction in just a single performance parameter would be sufficient to resolve this problem. This work expands our understanding of bacterial gene regulation under growth arrest and lays the groundwork for new design paradigms that will be essential in ensuring the safe and reliable performance of synthetic biology systems in real-world environments.

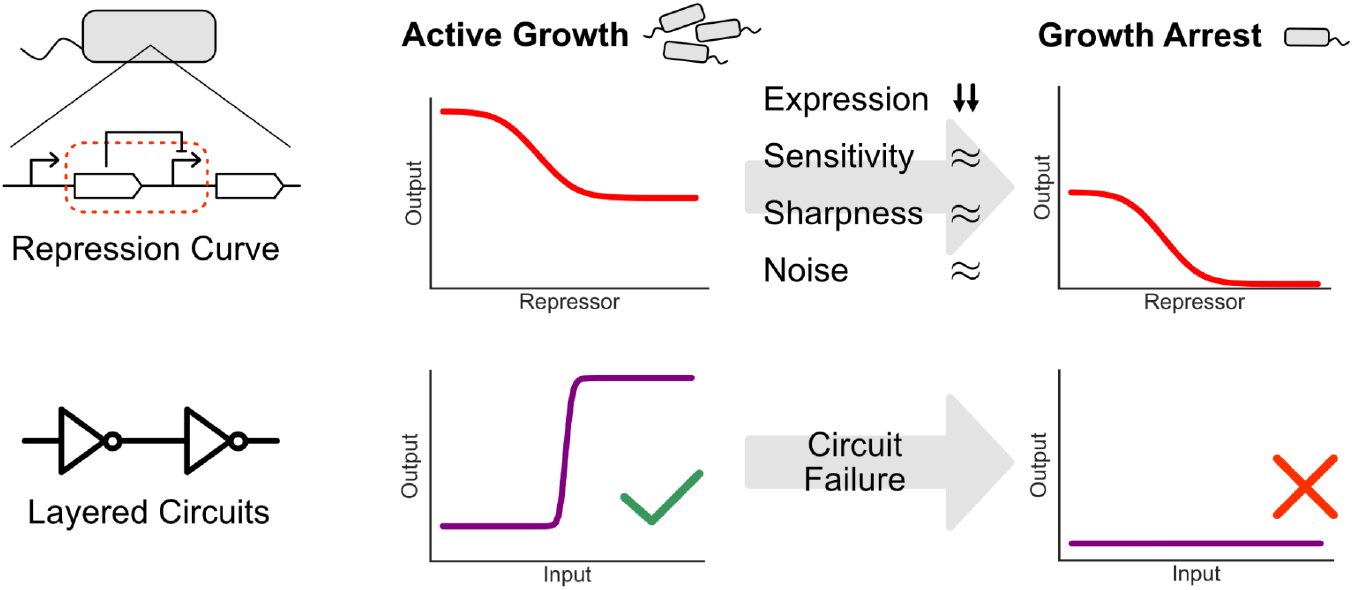

## Introduction

Genetic circuit engineering within living cells must contend with the fact that the physiological state of the host cell will change under different environmental conditions and thereby affect the behavior of the circuit (1–3). Ensuring predictable and reliable circuit performance across different conditions is therefore a central goal of synthetic biology. Such efforts are becoming increasingly important as the field moves towards developing engineered microbes intended for use outside of controlled laboratory environments, such as the gut, soil, and engineered structures (4, 5).

Growth arrest is thought to be one of the most common physiological states for microbes in nature (6). A number of global physiological responses associated with growth arrest are known. These include a reduction in the number of ribosomes and RNA polymerases, compaction of the DNA and modifications to supercoiling, and major changes to metabolism (6–8). In order for complex genetic circuits to maintain long-term, reliable function in natural environments, it will be important to understand how such cellular responses to growth arrest affect the performance of these circuits’ constituent parts.

Many of the most complex genetic circuits constructed to date have been built using a repression-based architecture, wherein transcriptional repressors are used to implement modular NOT and NOR gates that can be wired together to form, in theory, logic circuits of arbitrary complexity (9, 10). The behavior of the individual repressors within the circuit are characterized by their repression curves, which are typically represented by the Hill relation

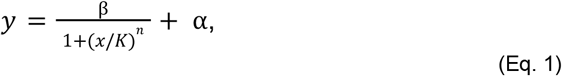

where the input *x* represents the repressor concentration and the output *y* represents the expression level from the repressed promoter. The parameters *β* and *α* determine the unrepressed and maximally-repressed expression levels of the output gene, and *K* and *n* represent the sensitivity and sharpness of the response to repressor concentration.

Proper circuit function requires that layered repression curves are properly aligned– a simple way to represent this condition is to say that the sensitivity of the downstream repressor must lie within the dynamic range of the upstream repressor, i.e. *α*_1_ *< K*_*2*_ *< β*_1_+*α*_1_. Factors like the Hill coefficient *n* or the level of gene expression noise affect the magnitude by which these inequalities need to hold. Characterizing the full repression curve is therefore essential to predicting the performance of the larger circuit. We illustrate the complexities associated with repressor composition in Fig. 1 using a NOT-NOT circuit, which buffers signals via OFF-to-ON logic and forms the basis for more sophisticated circuit architectures (9, 11)

**Fig. 1:**
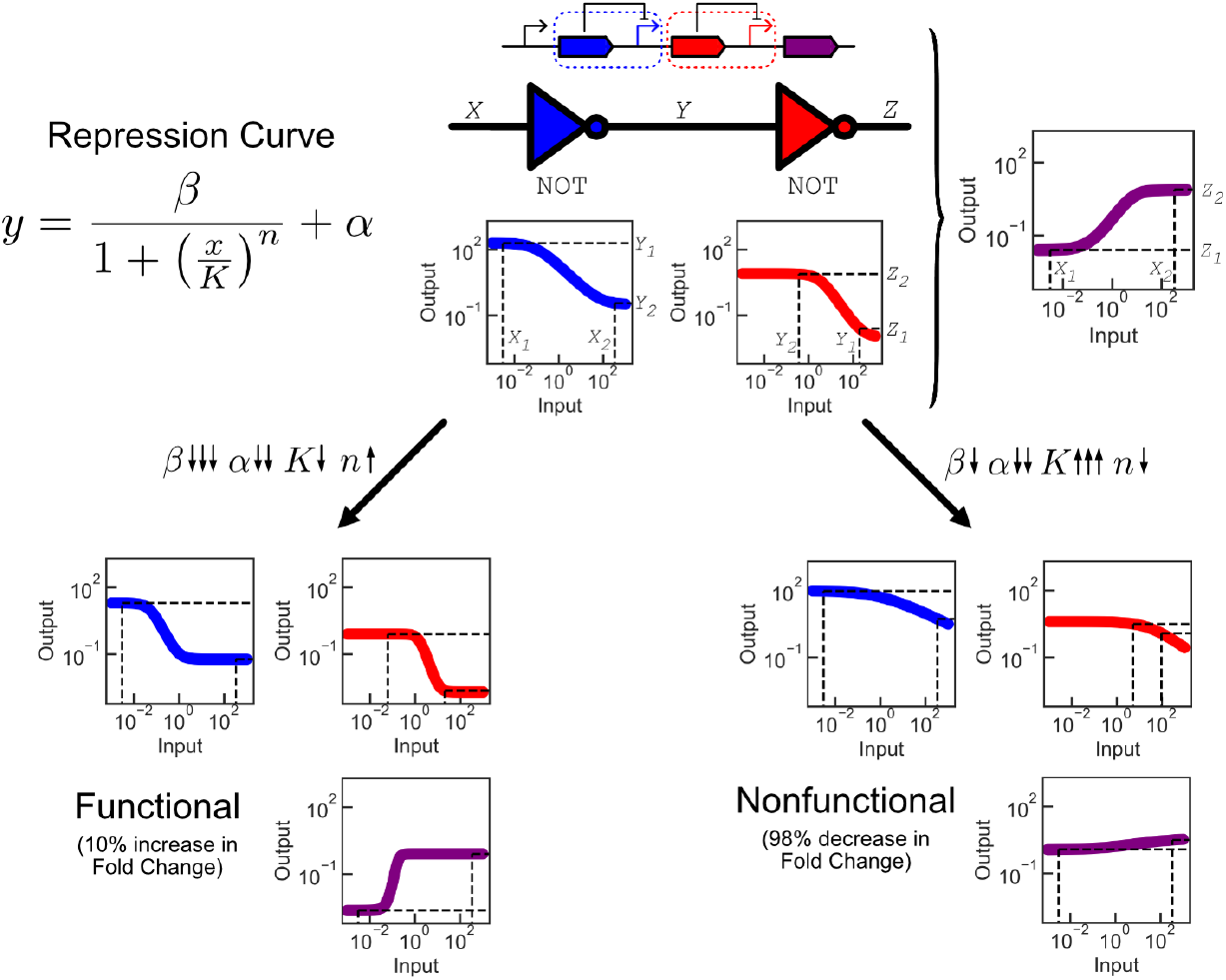
Schematic representation of the composition of repressors into larger circuits, illustrated by a NOT-NOT circuit. Inputs into the first gate (X) are transformed into output values (Y) according to the properties of the repression curve, which then become inputs into the second gate to generate outputs of the overall circuit (Z). Global changes to the parameters of the repression curve can lead to both enhancement (bottom left) and failure (bottom right) of circuit performance. The top circuit has parameters β_1_=200, α_1_=0.3, K_1_=0.1, n_1_=1 and β_2_=8, α_1_=0.01, K_1_=3, n_2_=1.5. The left transition scales all β,α,K,n values by 1/10,1/5,1/2,2, respectively. The right transition scales all β,α,K,n values by 1/2,1/5,10,1/2, respectively. All units are arbitrary.

However, despite the prominence of growth arrest in application environments like the gut or soil, there has been little investigation of its impact on the performance of engineered genetic components, including repressors. As a consequence, it is not clear *a priori* how the cell’s physiological responses to growth arrest would affect any of the parameters governing the repression curve, and whether these impacts would preserve or inhibit the ability of a circuit built from these repressors to function under growth arrest.

In this study, we set out to investigate whether there are any systematic effects associated with growth arrest on the performance of genetic repressors. We used a library of NOT gates based on dCas9-mediated repression of the T7 promoter at differing binding locations, engineered in *Escherichia coli*, as a model system to address this question. We found that functional NOT gates tended to retain their function under growth arrest with a slightly reduced fold change, but that this occurred at expression levels that were ∼100-fold lower than in active growth. The shape parameters of the repression curves tended to be robust to the change in growth state, with the sensitivity and sharpness of the curves affected only minimally. Similarly, gene expression noise of NOT gate outputs increased only moderately under growth arrest. We further confirmed that these conclusions were consistent with characterization of additional sets of NOT gates based on the PhlF repressor and a dCas9*:PhlF fusion protein. Within each set of gates, we generally observed that the repression behavior was more similar across gates under growth arrest than in active growth.

These results show that current design paradigms for layered repression-based circuits cannot lead to reliable circuit performance under growth arrest conditions, but that this failure is almost entirely tied to the change in a single parameter, *β*. As such, our findings reveal directions for future research by which these concrete design challenges could be overcome. Taken together, this work enhances our understanding of how growth arrest affects the quantitative properties of gene regulation and helps develop a foundation for genetic design principles in natural environments.

## Materials and Methods

### Construction of genetic circuits

All circuits were constructed using 3G Assembly (12) using parts from the CIDAR MoClo Extension Part Kit (13), with the exception of custom gRNAs and fusion proteins created for this work (see following section). NOT gates were assembled in two cassettes: one contained P_LacO_-driven T7 RNA Polymerase and P_T7_-driven sfGFP and the other contained P_Tet_-driven repressor:mScarlet3 fusion protein and P_OR1OR2_-driven gRNA. The two cassettes were integrated into the Lambda and P21 integration sites, respectively, of the *E. coli* genome using the pOSIP clonetegration system plasmids KL and CT (14). The resulting strains are kanamycin and chloramphenicol-resistant. The background strain for all experiments was *E. coli* Marionette MG1655 (15), which natively expresses the LacI and TetR repressors which regulate P_LacO_ and P_Tet_, respectively. The existing chloramphenicol resistance gene in the Marionette strain was excised prior to circuit insertion using the temperature-sensitive plasmid pE-FLP (14), which was then cured by repeated passaging of the strain at 37C. Sequences of all circuit components are provided in Supplemental Table 1.

For gates where the PhlO sequence was inserted between genetic parts of the P_T7_-driven sfGFP cassette in various locations and orientations, the relevant portion of the cassette was commercially synthesized as a dsDNA fragment flanked by BsaI cut sites and used in lieu of the relevant part plasmids in the Golden Gate step of 3G Assembly. All subsequent construction steps proceeded as described above.

### Construction of gRNAs and fusion proteins

gRNAs were constructed as part plasmids in CIDAR MoClo format (16) with a Hammerhead ribozyme upstream of the gRNA sequence to standardize the 5’ sequence of the resulting RNA. Different gRNA variants were constructed by PCR amplifying the gRNA part plasmid into two segments that exclude the 20bp variable sequence using custom primers containing the new 20bp variable sequence as overhangs, and religating the plasmid using Gibson assembly.

Repressor:mScarlet3 fusion proteins were constructed as part plasmids in CIDAR MoClo format by using Gibson assembly to combine PCR amplicons of the mScarlet3 part plasmid (without the start codon) and the dCas9 or PhlF part plasmids (without the stop codon) with a 2x GGGS linker as the overhang. mScarlet3 was always attached to the C terminus of the repressor.

The dCas9*:PhlF fusion protein was created by commercially synthesizing a dsDNA fragment containing the modified region of dCas9 with overhangs to PCR amplicons of the dCas9 and PhlF:mScarlet3 part plasmids, and using Gibson assembly to combine the fragments into a full dCas9*:PhlF:mScarlet3 part plasmid in CIDAR MoClo format.

### Growth media design

Carbon-limited media was created by first preparing a stock of freshwater base (100g NaCl, 40g MgCl_2_ᐧ6H_2_O, 10g CaCl_2_ᐧ2H_2_O, 50g KCl in 1L Water), a stock of 0.5M NH_4_Cl to act as a nitrogen source, a stock of 1M Na_2_SO_4_ to act as a sulfur source, and a stock of 100 mM KH_2_PO_4_ at pH 7.2 to act as a phosphorous source. To create 1L of carbon-limited media, 10 mL of freshwater base, 10 mL of the nitrogen source, 250 uL of the sulfur source, and 1 mL of the phosphorous source were combined with one 10 mL vial of Trace Mineral Supplement (ATCC MD-TMS) and added to MilliQ water to a total volume of 1L. The resulting mixture was then filter-sterilized.

For active growth conditions, commercial M9CA minimal media (Teknova M8010) was used, which contains 1% glucose, 0.1% casamino acids, 0.5 ug/mL thiamine, 0.2 mM magnesium sulfate, and 0.1 mM calcium chloride at pH 7.0.

### Strain growth and induction conditions

Glycerol stocks of engineered strains were streaked onto LB agar plates with appropriate selection markers and grown overnight at 37C. One colony from each strain was then picked and inoculated into 3mL M9CA media with antibiotic selection (25 ug/mL kanamycin and 17 ug/mL chloramphenicol) and grown for 24h in a 15 mL tube in a shaking incubator at 30C and 250rpm to generate a stationary phase culture. The optical density (OD600) values of the resulting cultures were then measured with a spectrophotometer. For each strain, fresh batches of M9CA media with selective antibiotics were prepared in 96-well deep-well culture plates (NEST prod. no. 502062) with 12 500 uL wells per strain. 11 of these wells were used to titrate anhydrotetracycline (atc; Sigma-Aldrich cat. no. 37919) concentrations to induce P_Tet_-driven expression of the repressor at concentrations spanning 0 to 200 ng/mL (see Supplemental Figures 1-24 for specific concentrations used for each experimental condition). Overnight cultures were diluted 1:10,000 into these wells and returned to the shaking incubator covered with a Breathe-Easier Sealing Film (Diversified Biotech BERM-2000).

Meanwhile, 1 mL of the overnight cultures for each strain were transferred into 1.5mL Eppendorf tubes and spun down at 1,377 g on a tabletop centrifuge for 10 minutes. The supernatant was removed and resuspended in 1 mL of carbon-limited media, and the spin procedure was repeated. The supernatant was again removed and replaced with 1 mL of carbon-limited media. The resulting solutions were then re-diluted into 7 mL of carbon-limited media to bring the final density of each culture to an OD600 of 0.0025. The dilute cultures were then split into twelve 200 uL aliquots per strain in 96-well polystyrene microplates (Corning prod. no. 3370), and 11 of these wells were induced with varying concentrations of atc. The plates were then covered with the provided plastic lid and placed into the shaking incubator under the same conditions as above. Antibiotics were not added to the carbon-limited media to avoid adding potential additional stressors to the cells.

All cultures were grown for 24 hours, after which 1 mM IPTG (Sigma-Aldrich cat. no. 420322) was added to the 11 atc-induced wells and the cultures were returned to the shaking incubator to continue growing for an additional 24 hours. Throughout this whole period, the M9CA cultures were diluted 1:1000 every 12 hours by transferring 0.5 uL media from each well into 500 uL of fresh media that contained the appropriate antibiotic and inducer concentrations for each condition.

24 hours after the IPTG induction, the M9CA cultures were transferred into 96-well flow cytometry plates via a 1:400 dilution into 200 uL of fresh M9CA media, and all cultures were measured via flow cytometry.

### Flow cytometry

Cells were measured on a Cytoflex S flow cytometer. Forward and Side Scatter thresholds to detect bacteria, as opposed to spurious debris or instrument noise, were determined manually based on comparisons to readings from samples containing only media. sfGFP expression was measured using 488 nm excitation and a 525/40 nm bandpass filter, while mScarlet3 expression was measured using 561 nm excitation and 610/20 nm bandpass filter. At the beginning of each measurement session, calibration beads (Spherotech RCP-30-5A) were measured to allow conversion of arbitrary fluorescent units into absolute fluorescence units (Molecules of Equivalent Fluorescein (MEFL) for sfGFP and Molecules of Equivalent PE-TexasRed (MEPETR) for mScarlet3) by fitting to an 8-point standard curve. M9CA cultures were measured until 50,000 putative bacterial events were observed, while carbon-limited cultures were measured for 1 minute at the maximal standard flow rate of 60 uL/min, which typically yielded 5-10,000 putative bacterial events per sample.

### Analysis of flow cytometry data

Flow cytometry data were analyzed using the Python package FlowCal (17). Fluorescence values were converted into absolute units using the appropriate bead data from each run using FlowCal’s built-in calibration functions. We then performed density gating on each set of measurements to keep only the events that were in the densest region of the forward scatter/side scatter plot in order to better exclude non-singlet events from our analysis. We used a density threshold of 0.4, meaning we discarded 60% of the measured events from each sample.

### Noise analysis and determination of non-unimodality

In order to compare the distributions of sfGFP fluorescence values from each sample across different conditions, we performed the following normalization procedure. For a given sample, we extracted the measured GFP values, log-transformed them, and divided each resulting value by the median of this distribution. We then generated a kernel density estimate (KDE) from these values using the scipy.stats.gaussian_kde() function (18) with default parameters and then standardized the height of the KDE by normalizing it to a height of 1. This procedure allowed us to overlay all GFP distributions from all conditions on top of each other to analyze the shape and width of the distributions. Distribution widths were calculated by finding the first and last points at which the KDE intersects a value of 0.2, and dividing these points to obtain the width of this distance in logarithmic space.

Distributions were classified as ‘notably non-unimodal’ if their derivative was negative at any point below the median (which is rescaled to be one). Buffer values of 0.1 below the median and 0.005 below zero for the derivatives were applied to accommodate noise in the data.

### Bayesian Parameter Estimation

Markov Chain Monte Carlo (MCMC) was implemented using the Python package emcee (19). Priors for each of the parameters in the Hill repression function (Eq. 1) were set as lognormal distributions with scale and shape parameters μ _α_ = *_ylow_*, σ_α_ = 1, μ_β_ = *y*_*high*_ − *_ylow_*, σ_β_ = 0. 5, μ_*K*_ = 0. 1, σ_*K*_ = 1, μ_*n*_ = 1, σ_*n*_ = 0. 5, where *y*_*high*_ is the geometric mean of the median unrepressed GFP values from each replicate and *y*_*low*_ is the geometric mean of the median maximally-repressed GFP values from each replicate. Median (RFP,GFP) fluorescence values from each measured sample were extracted and the values from all induction conditions and replicates were pooled together for each gate to generate 11*3=33 data points along the repression curve for each gate. MCMC was run with 32 walkers for 10,000 iterations fitting the data against the Hill repression function. Autocorrelation analysis was used to assess convergence by confirming that the maximal autocorrelation value *τ*_*max*_ was less than the number of iterations divided by 50.

## Results

### Design of general NOT gate architecture for measurement in growth arrest

Although we do not know *a priori* exactly how growth arrest will affect the behavior of engineered NOT gates, our general understanding of microbial growth arrest suggests that our assay will need to be sensitive enough to detect low levels of gene expression and be able to capture cell-to-cell variability in repression. Both of these points require modifications to the conventional procedures used in the field to measure repression curves. Typically, repression curves are measured by driving the expression of a fluorescent protein from a repressible promoter and expressing its cognate repressor from a separate inducible promoter (Fig. 2a). By adding different concentrations of inducer in separate experiments, one generates different concentrations of repressor that lead to different expression levels of the output fluorescent protein.

**Fig. 2:**
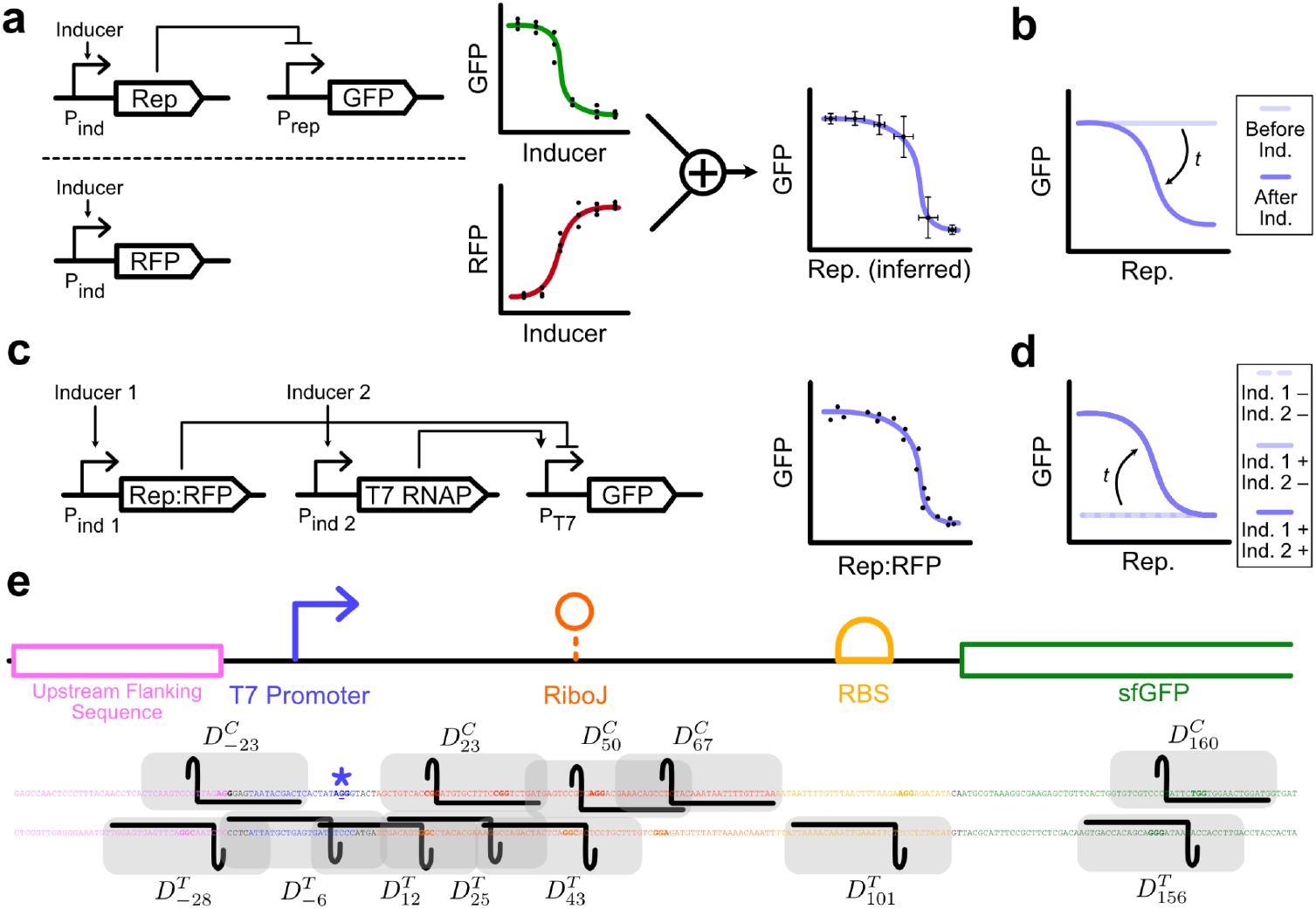
Design of NOT gates compatible with growth arrest measurements. (**a**) Schematic of the conventional procedure for measuring repression curves. The repressor (Rep) is expressed from an inducible promoter (P_ind_) to repress the expression of a fluorescent protein (GFP) from the repressible promoter (P_rep_). In a separate experiment, P_ind_ is used to express another fluorescent protein (RFP) to determine the expression level associated with a particular inducer concentration. This data is used to infer the repressor concentration in the first experiment, yielding a repression curve. (**b**) In the gate architecture in (a), GFP expression starts high in all conditions because P_rep_ is active in the absence of repressor. The experimenter must wait for GFP levels to fall to their repressed level over time via dilution by cell division or active degradation. (**c**) Design of the T7-based NOT gate used in this work. The repressor of the T7 promoter P_T7_ is fused to RFP, so single-cell measurements of both repressor and GFP levels can be made simultaneously. (**d**) For the gate architecture in (c), the GFP levels remain low until T7 RNAP is induced, ensuring that all observed GFP was expressed when P_T7_ was in a repressed regime. (**e**) Representation of dCas9 binding footprints for the 12 designed NOT gates. PAM sequences are bolded and the +1 transcriptional start site is underlined and marked with a star.

Importantly, because the repressible promoter is unrepressed prior to the start of the experiment, one must wait for any output protein produced prior to repressor induction to be removed from the cell in order to accurately measure the expression level from the promoter in its repressed state (Fig. 2b). This removal relies on either active degradation or dilution of the output protein through cell division, both of which pose challenges for measurements taken in growth arrest. One cannot rely on dilution to remove the initial output proteins, because cells rarely divide under nutrient starvation. Meanwhile, applying active degradation to the output protein decreases its concentration associated with a particular level of promoter activity. As global gene expression levels are already predicted to be much lower under growth arrest (7), decreasing the signal strength in this way is undesirable.

We resolved these challenges by choosing to base our NOT gates around repression of the T7 promoter (Fig. 2c). Because the T7 promoter does not exhibit cross-activation from *E. coli*’s native RNA polymerase, it remains inactive until its cognate RNA polymerase (T7 RNAP) is expressed. As such, we can perform a two step procedure where we first induce the repressor, and afterwards induce the T7 RNAP in order to activate expression from the T7 promoter. This ensures that all observed output proteins were expressed when the promoter was under a repressed regime (Fig. 2d).

Another challenge with applying conventional repression curve measurement techniques to growth arrest conditions is that the conventional approach does not measure the concentration of repressor directly. Instead, the repressor concentration is assumed to be equivalent to that of a separate fluorescent protein expressed from that same promoter under the same conditions, in a separate experiment (Fig. 2a). This approach does not allow us to measure cell-to-cell variability in repression because one cannot quantify the repressor concentration against the output protein concentration within the same cell.

We resolved this challenge by choosing to fuse our repressor to an orthogonal fluorescent protein, allowing direct single-cell measurements of repressor concentration alongside output protein concentration (Fig. 2c). Furthermore, in order not to conflate the effects of cell-cell variability on gene expression with those from within-cell variability in plasmid-borne circuit copy number, we chose to integrate the entire system onto the *E. coli* genome.

### Design of NOT gate library

Having created a general gate architecture in which fluorescent protein-fused repressors target the T7 promoter, we next needed to choose the specific set of gates to test. Existing repression-based circuits have typically been constructed either from libraries of genomically-mined transcription factors (20) or from dCas9 (21). Transcription factors exhibit a large diversity among various properties that affect repression, such as the protein’s size and its tendency for multimerization. They also vary in their actual mechanism of repression, including for example DNA looping, steric hinderance, or even stabilization of host RNA polymerases (22, 23). This means that if two gates driven by different repressors behave differently under growth arrest, it will be difficult to disentangle which molecular factors led to these different responses.

We therefore chose to use dCas9-based repression as the basis for our NOT gate library, so that all of the gates share the same repressor protein and vary only in where it binds. It is known that dCas9 can repress by inhibiting either transcriptional activation or elongation based on its binding position, and that in the latter case the binding orientation has a large impact on repression strength (24). Although to our knowledge there has been no systematic investigation of how dCas9 binding location affects the general properties of the repression curve as a whole, we reasoned that the existing evidence nonetheless suggests that varying dCas9 binding position should create a library of NOT gates with diverse repression curve profiles.

We designed a construct expressing a sfGFP gene from the T7 promoter, incorporating the RiboJ ribozyme to serve as a genetic insulator by standardizing the 5’ untranslated region of the mRNA (25). We identified 12 PAM sites on both strands spanning 30bp upstream to 160bp downstream of the transcriptional start site and constructed gRNAs to target dCas9 to these positions, generating a library of 12 different NOT gates (Fig. 2e). We label each gate as D^{T/C}^_N_, where the superscript indicates whether dCas9 binds to the Template or Coding strand and the subscript N indicates the location of the midpoint of the predicted 33bp dCas9 binding footprint with respect to the transcription start site (26). We also created a 13^th^ gate with an off-target gRNA, D^0^, as a negative control.

Because measuring the repressor concentration is a critical part of our assay, we fused dCas9 to a red fluorescent protein (mScarlet3) and expressed it from an inducible promoter (P_Tet_), while each gRNA was expressed from a strong constitutive promoter (P_OR1OR2_) to ensure that it is in stoichiometric excess. While dCas9-based NOT gates typically express dCas9 constitutively and titrate the concentration of gRNA, these two approaches are functionally equivalent in that they both lead to the titration of the amount of active repressor (the dCas9:gRNA complex). The T7 RNAP was separately expressed from an orthogonal inducible promoter (P_LacO_). Both inducer molecules, atc and IPTG, are not metabolized by *E. coli*, ensuring that cells remain in growth arrest even upon addition of inducer.

### Design and validation of measurement assay

Studies in the literature utilize a number of different approaches to induce growth arrest, which can have different implications for the resulting physiological response. We chose to follow the general approach used by Bergkessel and Delavaine, which allows a culture of cells to naturally reach stationary phase before washing and diluting them into carbon-limited media (27). This approach simulates the gradual entry into starvation that is likely more representative of natural environments while also avoiding potential confounding effects from the high densities associated with stationary phase cultures.

We then set out to design a measurement procedure that allows the behavior of a genetic circuit in the same population of cells to be compared under two different growth conditions. While synthetic biologists often measure the performance of their circuits in both exponential phase and stationary phase to assess robustness across growth phases, this is typically done by activating the circuit in exponential phase, measuring its output, and then waiting until that same population reaches stationary phase and then re-measuring the output (15, 20). This approach captures how the circuit’s performance persists over time but does not capture its intrinsic behavior under growth arrest, as the circuit is not activated during growth arrest specifically, and outputs from its prior activation during exponential phase could carry over into stationary phase.

We therefore designed a measurement procedure that can independently measure a circuit’s behavior under both growth arrest and active growth (Fig. 3a, Methods). An overnight culture of cells engineered with the NOT gate is split into 24 different cultures, half of which are washed and diluted into carbon-limited media alongside the addition of 11 different concentrations of atc to induce dCas9 expression to various levels. 24h later, 1 mM IPTG is added to 11 of the cultures to induce T7 RNAP expression, with the 12th culture remaining as an uninduced negative control. The same induction procedure is applied to the other 12 cultures from the initial split, except these are repeatedly diluted into fresh batches of the original growth media to ensure the cells remain actively growing. 24h after the T7 RNAP induction, all cultures are measured for mScarlet3 and sfGFP expression via flow cytometry.

**Fig. 3:**
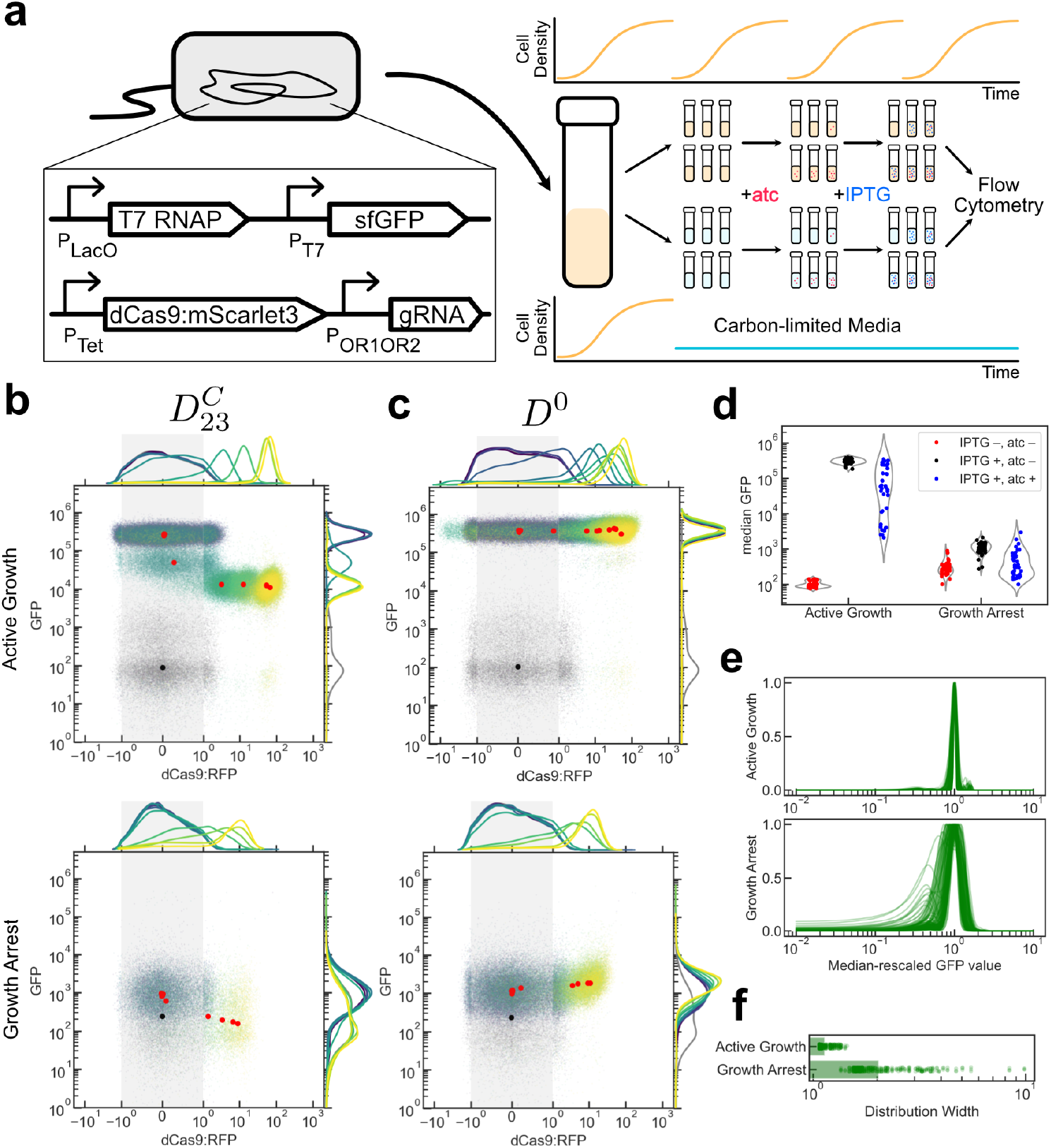
Measurement of NOT gate repression curves under active growth and growth arrest. (**a**) Schematic representation of measurement assay. (**b**) Representative example of a repression curve in both active growth and growth arrest. (**c**) Representative example of the negative control circuit driven by an off-target gRNA. Points in (b,c) are colored according to the atc concentration, with grey points from the condition where no IPTG was added. Circles represent the median GFP,RFP values for each condition. RFP values are plotted on a symmetric log scale (28) where the grey shaded region is linear. Full plots of each of 3 replicates for each gate are in Supplemental Figures 1a-13a. (**d**) Median GFP values across all gates at selected induction conditions. IPTG –/+ corresponds to 0 or 1 mM induction of T7 RNAP and atc –/+ corresponds to 0 or maximal (200 ng/mL for active growth, 20 ng/mL for growth arrest) induction of repressor. (**e**) Kernel density estimates (KDEs) of log-transformed GFP values across all experimental conditions where T7 RNAP was induced, rescaled to the median of the distribution and normalized to a maximal value of 1. (**f**) Width of each KDE in (e), defined by the distance in logspace between the two points where the KDE crosses the value of 0.2. Bars show the geometric mean.

Our assay was able to measure repression behavior in both active growth and growth arrest for the NOT gates in our library, with dCas9 concentrations spanning a wide range of values (from 0 to ∼100 MEPETR in active growth and to ∼10 MEPETR in growth arrest) and driving a decrease in GFP levels (Fig. 3b, Supplemental Fig. 1a-12a). The off-target gate D0 did not show a decrease in GFP associated with increasing dCas9 concentration (Fig. 3c, Supplemental Fig. 13a), indicating that observed decreases in GFP level are indeed due to the impact of active repression by dCas9.

We also saw that the unrepressed GFP level was similar across all gates within a growth condition, suggesting that leaky expression of dCas9 in the absence of induction was not a notable issue (Fig. 3d). Additionally, we saw a clear separation in GFP values between conditions where T7 RNAP was induced or uninduced in the absence of repressor across all gates, suggesting that leaky expression of T7 RNAP was also not an issue (p < 10^-6^ for both growth conditions, paired *t* test). However, the overall dynamic range of P_T7_ activation under growth arrest (3.75-fold) was significantly lower than in active growth (3000-fold). This reduction in dynamic range was primarily due to growth arrest imposing a large reduction in P_T7_’s ON state (geometric mean of 300,000 MEFL to 1,000 MEFL), as compared with the smaller increase in P_T7_’s OFF state (geometric mean of 100 MEFL to 275 MEFL) (Fig. 3d). Overall, these results showed that our assay is able to reliably measure repression curves from our NOT gate library in both active growth and growth arrest.

### Gene expression noise in NOT gate outputs increases moderately under growth arrest

We next investigated how growth arrest impacted the level of noise in the NOT gates’ output. We plotted kernel density estimates of the GFP values from every experimental condition where T7 RNAP was induced and rescaled them to their median value to overlay them against each other (Fig. 3e). GFP distributions in active growth were dominated by a tight peak around the median, as expected. GFP distributions in growth arrest, however, tended to be wider and 24% of them exhibited notable non-unimodality, compared to none in active growth (Methods). Calculating the width of the rescaled distributions found that growth arrest increased the average distribution width by 1.8-fold (Fig. 3f). These results show that growth arrest applies a consistent but moderate increase in the noise of NOT gate outputs.

### dCas9 binding location affects repression curve properties during active growth

We next analyzed our data to determine whether our library of NOT gates, in which gates differ from each other based on the binding location of dCas9, indeed generated a diversity of repression curves as predicted. Although prior work has shown that dCas9 binding location affects the extent of overall repression (24, 29, 30), to our knowledge our dataset is the first to measure full repression curves for different dCas9 binding locations. In order to extract the parameters of the repression curve from our measurements, we performed Bayesian Parameter Estimation on the data from each gate to obtain a distribution of values for each of the four parameters in the Hill repression function (Eq. 1) for each growth condition (Fig. 4a).

**Fig. 4:**
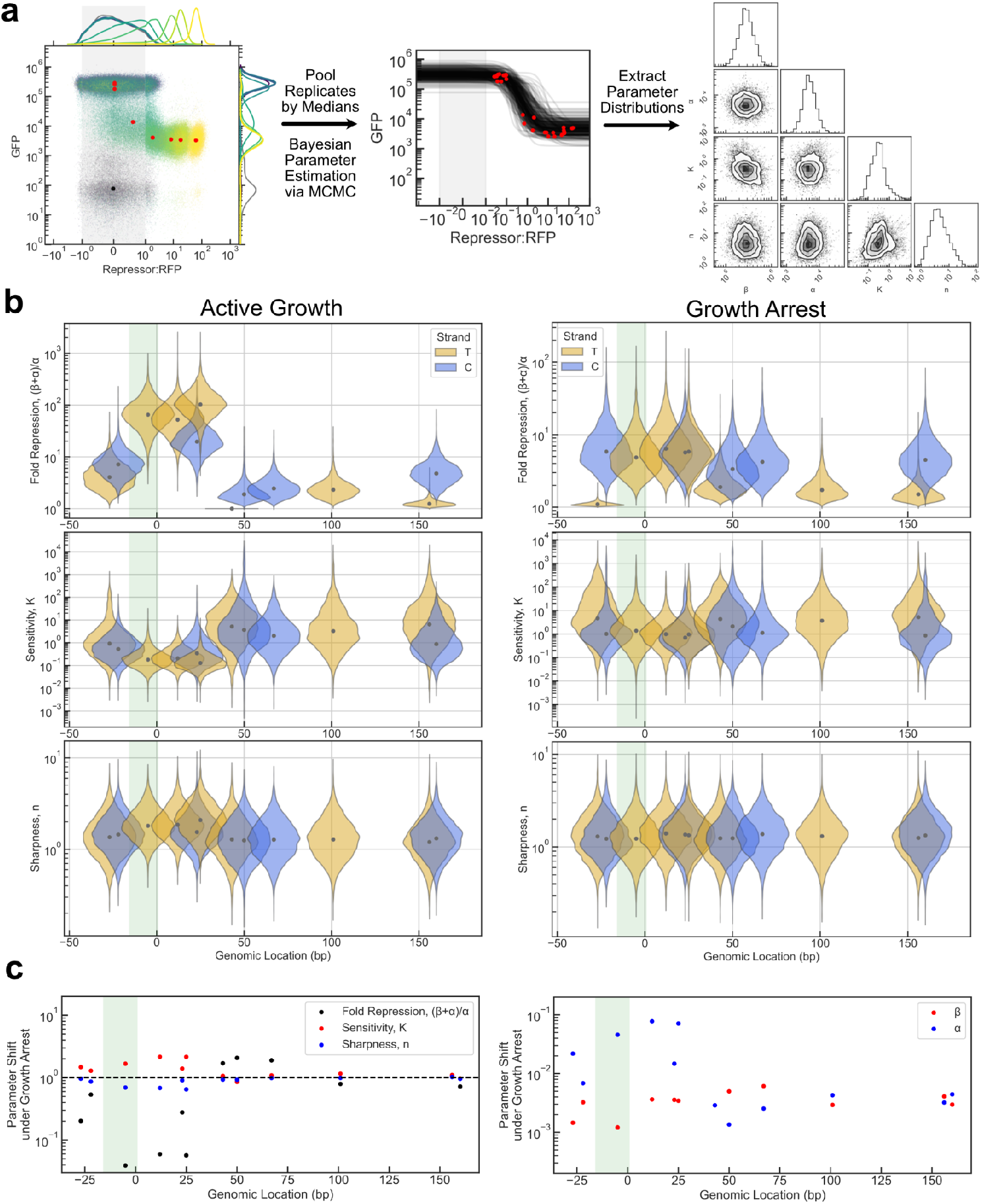
Repression curve behavior of dCas9 NOT gate library. (**a**) Schematic of the Bayesian Parameter Estimation (BPE) procedure for obtaining distributions of parameter values. Median (GFP,RFP) values from each induction condition are pooled across replicates to generate a master repression curve, to which the Hill Repression function (Eq. 1) is fit using Markov Chain Monte Carlo (MCMC)-based BPE. Full plots of each NOT gate in each growth condition are given in Supplemental Figs. 1b-13b. (**b**) Posterior parameter distributions for fold repression (β+α)/α, K, and n for each NOT gate in active growth and growth arrest, derived from three biological replicates measured on three different days. Dots represent medians, and each distribution is scaled to a constant width that marks the size of the dCas9 binding footprint on the genome. The location of the T7 promoter sequence is marked by the green shaded region. (**c**) Shifts in median parameter values from active growth to growth arrest for all repression curve parameters for all NOT gates, plotted against genomic location as in (b). The β shift is ommitted for gates with no repression in a growth condition as β = 0 when the fold repression is 1.

We first checked whether, under active growth conditions, the observed fold repression (*β*+*α*)/*α* followed the trends expected from existing literature (Fig. 4b, left). Gates where dCas9 bound upstream of the T7 promoter exhibited low levels of repression, while gates where dCas9 bound on or immediately downstream of the promoter showed strong repression (up to a maximum of 116-fold from Gate D^T^_25_). In contrast, binding positions further downstream showed very low levels of repression, with only Gate D^C^_160_ repressing at a similar level to the upstream-binding gates. This stronger repression from Gate D^C^_160_ is consistent with previous observations that dCas9 blocks transcriptional elongation more strongly when bound to the complement strand (24). Interestingly, Gate D^T^_43_ exhibited no detectable repression, despite binding only 18bp downstream, and on the same strand, from the most repressive gate D^T^_25_ (Supplemental Fig. 7).

Overall, our results suggest that dCas9 is a better repressor when blocking transcriptional activation than elongation, which goes against the results of prior work on native *E. coli* promoters (24, 29) but is consistent with prior results on the T7 promoter (30). Our data therefore further support the notion that dCas9’s interactions with the T7 RNAP may differ significantly from those with *E. coli*’s native RNAP.

Both the Hill coefficient *n* and the sensitivity *K* were fairly consistent across gates, with median *n* values ranging between 1.2 and 2.1 and median *K* values spanning a 2.6-fold range across the gates. Gates where the repressor blocked transcriptional elongation had a slightly lower sensitivity (i.e., higher *K*) to repressor concentration (Fig. 4b, left middle panel). The robustness of the Hill coefficient *n* to changes in binding position is expected, as *n* is thought to be a protein-intrinsic property. The fact that *K* values are higher for more downstream-binding gates, however, suggests that dCas9 may require higher concentrations to repress the T7 promoter via blocking transcriptional elongation than via blocking transcriptional activation.

### Impact of growth arrest on repression curve properties

We next characterized repression curves properties for these same gates under growth arrest to understand how these properties vary between the two growth conditions (Fig. 4b, right). Overall, repression values were much lower under growth arrest, reaching 6.6-fold repression at maximum. This is likely due in large part to the reduction in the overall possible dynamic range for the gates under growth arrest due to the decrease in P_T7_’s ON level (Fig. 3d). Position-dependent effects on repression strength were also homogenized under growth arrest, with upstream-binding and elongation-blocking gates able to reach similar levels of repression to the gates where dCas9 bound on or immediately downstream of the T7 promoter. Values of *K* and *n* also became more similar between gates under growth arrest, with median *K* values now spanning only a 1.4-fold range and median *n* values lying between 1.2 and 1.3.

Interestingly, Gate D^T^_43_ exhibited slight but detectable repression (1.7-fold) in growth arrest despite exhibiting no detectable repression in active growth (Fig. 4b, Supplemental Fig. 7). This result hints at the possibility that growth arrest, even though it decreases the level of maximal achievable repression, may impose physiological changes that broaden the capacity to repress across different repression schemes. One way in which this might occur is that differences in nucleoid organization between growth phases (31) could cause short-range DNA interactions that pervent dCas9 binding to the +43 position, present during active growth, to be weakened under growth arrest.

Further analyzing the changes in repression curve properties at the individual gate level, we found that all gates exhibited a significant drop in overall expression level, with *β* and *α* decreasing by a geometric mean of 321 and 99-fold, respectively (Fig. 4c). Depending on whether *α* decreased more, or less, than *β*, however, this led to either an increase or decrease in the overall repression strength, with the six gates binding closest to the promoter experiencing a reduction in repression strength (geometric mean of 7.9-fold reduction) and the remaining six gates experiencing a minimal changes that averaged to a slight increase in repression strength (geometric mean of 1.3-fold increase). Changes to *K* and *n* were minimal across all gates, with no gate shifting these parameters more than 2.2-fold (geometric mean increase in median *K* and *n* was 1.3 and 1.2-fold, respectively).

These results, taken together, suggest that the impact of growth arrest on repression curves is almost entirely dominated by a drop in overall expression level captured by the decreases in *β* and *α*, with a moderate increase in gene expression noise and minimal changes to the sensitivity and sharpness of the curve.

### Generalization of results to other repressors

We next asked whether the previous conclusions were specific to dCas9-mediated repression or whether they might generalize to NOT gates based on other repressors. To investigate this question, we turned to a previously-published system where dCas9 is mutated to reduce its native capacity for DNA binding (generating dCas9*) and then fused to the TetR-family transcriptional repressor PhlF (32). The resulting protein requires both a PhlF operator site (PhlO) and a PAM-targeting gRNA in order to bind to the DNA, and acts as an intermediate condition between dCas9-mediated repression and PhlF-mediated repression.

We identified 3 candidate positions where a PhlO sequence could be inserted in the regulatory region of our design (before the promoter, before RiboJ, and before the ribosome binding site) near a PAM site, and inserted the PhlO sequences in both possible orientations to create a set of 6 repressible constructs. We added spacer sequences as necessary to implement the previously-determined optimal 12bp spacing between the PhlO site and the PAM site (32) to ensure that these constructs could be repressed by either the dCas9*:PhlF fusion or by PhlF alone, thus creating a new set of 12 NOT gates (Fig. 5a). We label the fusion and PhlF gates as F^{T/C}^_N_ and P^{T/C}^_N_, respectively, following the previous convention. For P gates, N indicates the position of the midpoint of the PhlO site, while for F gates, N indicates the midpoint of the 66bp window that spans the dCas9 footprint and the PhlO site.

**Fig. 5:**
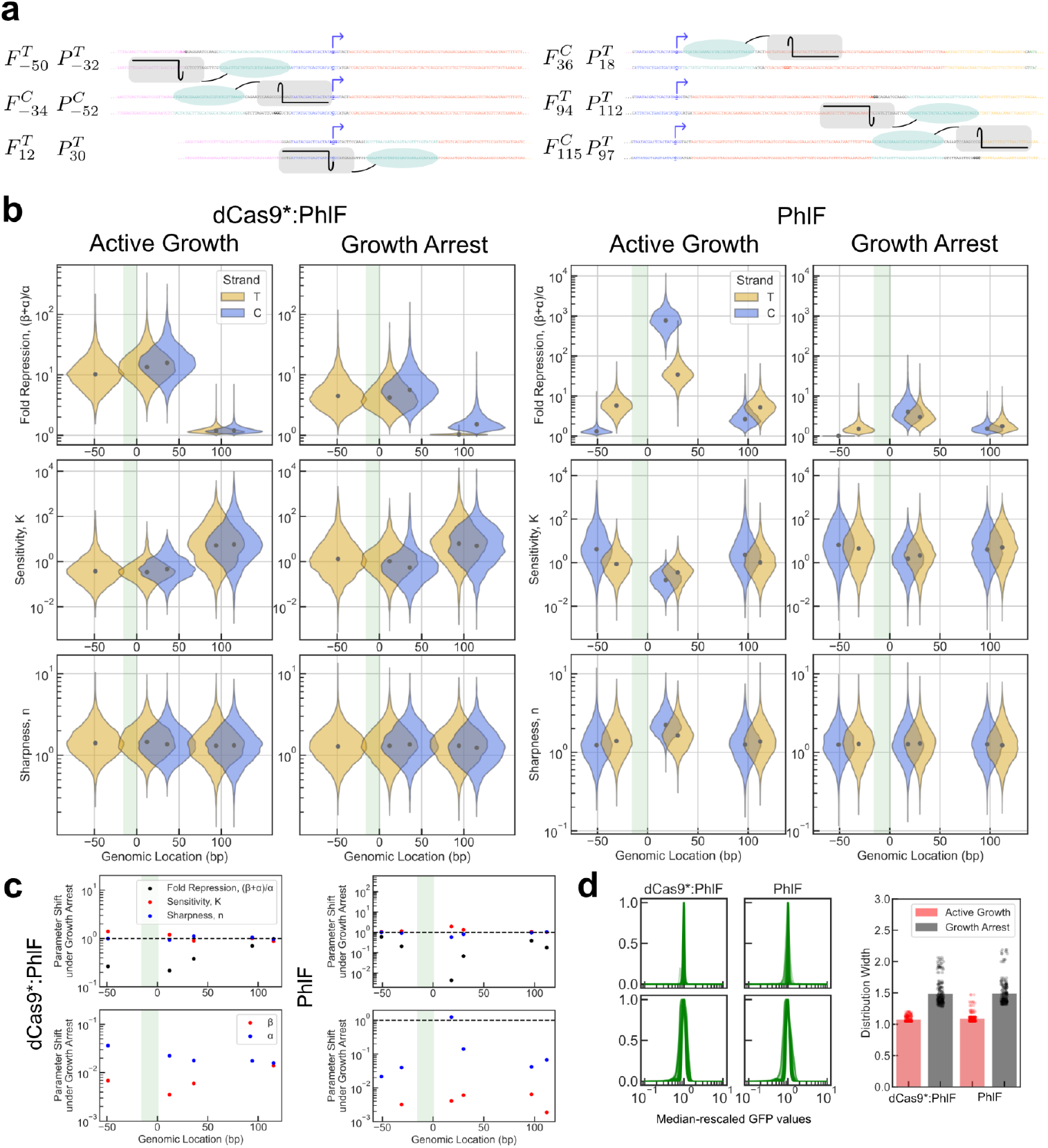
Measurement of repression curves from NOT gates driven by alternative repressors. (**a**) Schematic of repressed constructs. Genetic sequences are colored as in Fig. 1e, with the PhlO site added in green. The +1 transcriptional start site is bolded and underlined and marked with a hooked arrow, and PAM sequences are bolded. Each of the six repressible constructs generates two different NOT gates depending on whether it is repressed by the dCas9:PhlF fusion or by PhlF alone. (**b**) Posterior distributions for repression curve parameters in active growth and growth arrest for the dCas9:PhlF and PhlF NOT gates, derived from 3 biological replicates measured on 3 different days. Full repression curve measurements and MCMC outputs are shown in Supplemental Figs. 14-24. Distribution widths are scaled to a constant value that indicates the width of the repressor’s genetic footprint. The location of the T7 promoter sequence is indicated by the green shaded region. Dots represent the median of the distribution. (**c**) Shifts in median parameter values from active growth to growth arrest for all repression curve parameters for all gates, plotted against genomic location as in (b). The same parameters are plotted across graphs within a row. The β shift is ommitted for gates with no repression in a growth condition as β = 0 when the fold repression is 1. (**d**) Rescaled and normalized kernel density estimates (KDEs) of GFP values across all induced conditions (left), and widths of these KDEs defined by the width of the crossing at a value of 0.2 (right). Bars represent the geometric mean.

We fused mScarlet3 to the C-terminus of dCas9*:PhlF and PhlF and measured their repression curves using the same methodology as was previously applied to the dCas9 gates. Data from Gate F^C^_-34_ was thrown out after later investigation revealed that the gRNA was assembled incorrectly, leaving repression curve data for 11 additional NOT gates.

We first assessed whether these new sets of gates followed the same trends relating the repressor binding location to properties of the repression curve as was observed for the dCas9 gates under active growth (Fig. 5b). In both sets of gates, repression was strongest when the repressor bound immediately downstream of the +1 site (15-fold repression for Gate F^T^_12_, 865-fold repression for Gate P^T^_18_), although in the fusion gate library, the gate that bound upstream of the T7 promoter (Gate F^T^_-50_) exhibited comparable levels of repression. We note that in our hands, dCas9*:PhlF achieved lower levels of repression than were originally reported (15-fold maximum versus ∼50-fold repression in ref. 32). This discrepancy may be due to the fact that we used a full-length dCas9* protein in our fusion to keep the repressor identity more similar to our previous dCas9 gates, while the original study replaced dCas9’s HNH domain with a GGGSx2 linker.

The sensitivity, *K*, varied across only a 1.6- and 2.1-fold range for the fusion and PhlF gates, while the Hill coefficient *n* only ranged between 1.2 and 1.4 for the fusion gates and 1.3 and 2.1 for the PhlF gates. Overall, we concluded that the general relationship between repressor binding location and NOT gate behavior was similar in these gates as it was for the dCas9-based gates.

We then assessed whether the repression curves from these gates also shifted in response to growth arrest in a similar way to the dCas9 gates (Fig. 5c). As before, both the fusion and PhlF gates experienced a large decrease in overall expression levels, with median *β* values decreasing by a geometric mean of 149 and 253-fold and median *α* values decreasing by a geometric mean of 46 and 8.7-fold for the fusion and PhlF gates, respectively. *K* and *n*, as during active growth, exhibited minimal changes during growth arrest, with median *K* values increasing 1.1 and 1.3-fold and median *n* values increasing 1.0- and 1.2-fold across the fusion and PhlF gates, respectively. Interestingly, none of the tested gates exhibited notable multimodality in either growth condition, although the average widths of the GFP distributions under growth arrest were still wider than under active growth for both sets of gates, by 1.4-fold (Fig. 5d).

Taken together, these trends are broadly consistent with those that we observed in the dCas9-based NOT gate library, suggesting that the impacts of growth arrest on repression curves observed in this work may generalize to other repressors with different molecular implementations.

## Discussion

We have presented a systematic characterization of engineered genetic circuit components that directly compares their performance in active growth and growth arrest conditions. Doing so required the development of specialized NOT gate architectures and novel measurement approaches to quantitatively characterize repression curves in a way that is compatible with low gene expression levels and cell-to-cell variability. After measuring the performance of 23 different NOT gates based on three different repressors, we find that the impact of growth arrest on repression curves is dominated by a single effect, the significant decrease in output expression levels by two orders of magnitude. All other curve parameters, meanwhile, either experienced minor changes (such as the moderate increase in gene expression noise) or were unaffected (as were the sensitivity and sharpness of the curves).

Because the decrease in expression strength tended to occur asymmetrically, with unrepressed expression levels (driven by *β*) decreasing more than fully-repressed expression levels (driven by *α*), most NOT gates experienced a decrease in repression strength under growth arrest. However, in several cases, we found that individual gates were able to maintain similar performance measures between the two growth conditions. For example, even though Gate D^C^_-23_ experienced a 309-fold reduction in *β* and a 146-fold reduction in *α*, its overall fold change only decreased by 1.3-fold (going from 8.5-fold repression in active growth to 6.3-fold repression under growth arrest) while its sensitivity and sharpness experienced similarly small changes (1.3-fold and 1.2-fold increases in *K* and *n*) (Fig. 4b,c). This means that the repression curve almost completely preserved its shape and simply shifted downwards on the input/output plot as a consequence of growth arrest.

What are the implications of these results for the engineering of larger, more complex genetic circuits that can function under growth arrest? Recall that repression-based circuits require their component repression curves to align, which means that the sensitivity of the downstream repressor must lie within the dynamic range of the upstream repressor (*α*_1_ *< K*_*2*_ *< β*_1_+*α*_1_). Because we observed that *K* values change minimally while *β* and *α* values can decrease significantly under growth arrest, circuit failure occurs when the *β* value of the upstream circuit drops below the value of *K* for the downstream circuit. We illustrate this in Fig. 6 with a NOT-NOT circuit composed from the two strongest repressors from a library of 73 genomically-mined repressors (20). We can see that the reduction in expression level is strong enough to fully negate the predicted responsiveness of the circuit under growth arrest.

**Fig. 6:**
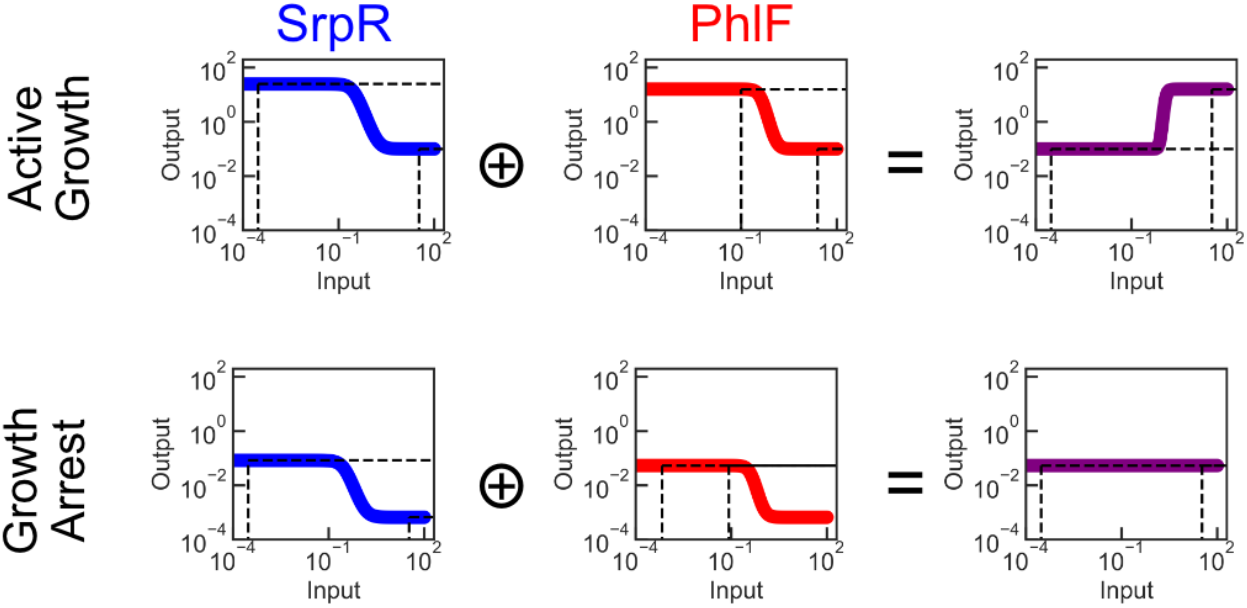
Illustration of the predicted impacts of growth arrest on circuit function. Predicted performance of a NOT-NOT circuit built from the TetR family repressors SrpR and PhlF. Although the circuit is in an always-ON failure mode under growth arrest, the decrease in overall expression levels means that its performance is nearly identical to that of an always-OFF failure mode in active growth. Repression curve parameters for active growth were obtained from ref. (20), and growth arrest curves were generated by scaling the active growth β and α values by 1/300 and 1/150, respectively, following the behavior of Gate D^C^_-23_. All values are in Relative Expression Units (REUs).

Despite this conclusion, however, it is nonetheless the case that genetic circuits with nontrivial complexity have been validated to function in natural environments where growth arrest is expected to occur, like the gut or the soil (33–35). Our results provide a possible explanation for this discrepancy. In an OFF-to-ON circuit like the NOT-NOT circuit described above, the misalignment of the repression curves under growth arrest leads to an always-ON behavior in the full circuit. However, because the level of the ON state under growth arrest is over 100 fold lower than it is in active growth, this state is functionally indistinguishable from an OFF state in active growth. This serendipitous alignment of effects means that growth-arrested cells would appear as false negatives for circuit function, and not impede the measured signal from the subset of cells in the population that are actively growing and exhibiting proper circuit function. Indeed, single-cell measurements of genetic circuit performance in natural environments have shown a high proportion of non-responsive cells even when genetic analyses confirm that they contain the circuit (36).

The conclusion from these results is that a circuit consisting of only a single NOT gate can indeed function under growth arrest, sometimes exhibiting very similar performance between active growth and growth arrest, provided that the measurement devices are calibrated to capture outputs with ∼100-fold lower signal values. However, the fact that the sensitivity of genetic components do not scale alongside the expression levels means that circuits consisting of layered gates will not, as a rule, function properly under growth arrest. Therefore, in applications where cells must consistently perform circuit computations specifically under growth arrested conditions, such as in long-term biomonitoring or in embedded structural materials, new genetic design paradigms will be required.

One possible approach would be to extend the expected timescale of circuit function, so that output proteins from each circuit layer are given a longer time to accumulate. A previous study has shown that exogenously-induced proteins can accumulate linearly in stationary phase *E. coli* cells over a period of at least 24 hours (37). If this effect persists over multiple days, then enough protein might accumulate to mitigate the reduction in *β*, allowing gates to be wired together and retain their expected performance even under growth arrest. In the NOT-NOT circuit proposed above, for example, mitigating the reduction in expression level by only 8-fold (bringing the growth arrest-associated reductions in *β* and *α* to 38-fold and 19-fold, respectively) would allow the overall circuit to exhibit a 10-fold response range under growth arrest (Supplemental Fig. 25). Under such a scheme, however, new strategies to resolve the challenges associated with circuit computation over weeklong timescales would need to be developed, like solutions for increased leak and alternative mechanisms of protein removal.

Synthetic biology holds great promise for transforming the physical world in ways that can advance both human and ecological welfare, but doing so will require the development of new sets of design principles to allow the safe and reliable performance of engineered biosolutions into real-world environments. This work takes a first step towards this goal by characterizing the impacts of microbial growth arrest on genetic circuit behavior to reveal which performance parameters are most affected, revealing new avenues for future research to address these challenges.

## Acknowledgments

We thank M. Bergkessel, J. Ciemniecki, and D. Newman for insightful initial discussions, and A. Halleran and S. Clamons for feedback on the manuscript. We thank A. Pandey for technical guidance on conducting Bayesian Parameter Estimation, and J. Tijerina and the staff of the Caltech Flow Cytometry Facility for technical assistance with data collection. This work was supported by the Resnick Sustainability Institute and by the Institute for Collaborative Biotechnologies through contract W911NF-19-D-0001 from the U.S. Army Research Office. The content of the information on this page does not necessarily reflect the position or the policy of the Government, and no official endorsement should be inferred.

## Author Contributions

Conceptualization - J.P.M.

Data curation - J.P.M.

Formal analysis - J.P.M.

Investigation - J.P.M., M.L.P.

Methodology - J.P.M., M.L.P.

Supervision - B.A.H., R.M.M.

Writing – original draft - J.P.M.

Writing – review & editing - J.P.M., M.L.P., B.A.H., R.M.M.

## Data and Code Availability

All raw data from this manuscript are uploaded to the Zenodo repository accessible at doi: 10.5281/zenodo.18415279 and code to reproduce all analyses and graphs in this work are available at https://github.com/jpmarken/GrowthArrestRepression.

