## Supplemental Data for "Quantitative measurement of synthetic repression curves reveals design challenges for genetic circuit engineering under growth arrest"

### Supplemental Material

#### Supplemental Figures

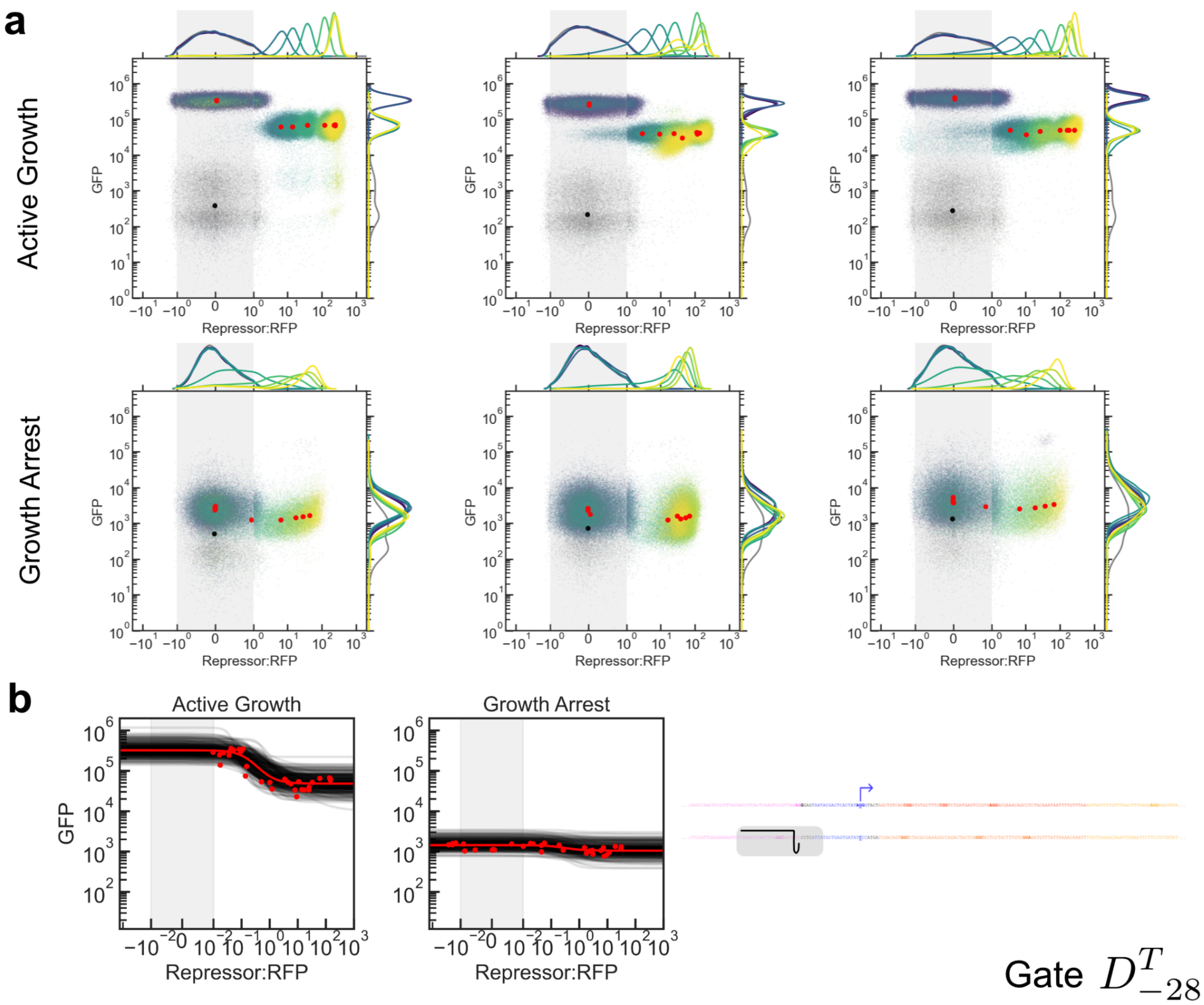


***Supplemental Fig. 1:*** *Repression curves for gate DT-28. (**a**) Flow cytometry data. Three biological replicates (columns) were measured on three different days. Dots represent medians of distributions for each induction condition, and colors represent atc concentrations. Induction conditions for atc were: Active Growth, Replicate 1: 0, 0.02, 0.06, 0.2, 0.43, 0.93, 2, 6, 20, 60, 200 ng/mL. Active Growth, Replicate 2: 0, 0.02, 0.2, 0.43, 0.93, 2, 3.5, 6.3, 11.3, 20, 200 ng/mL. Active Growth, Replicate 3: 0, 0.2, 0.43, 0.93, 2, 2.6, 3.5, 6.3, 11.3, 20, 200 ng/mL. Growth Arrest, all replicates: 0, 0.02, 0.06, 0.2, 0.43, 0.93, 2, 6, 20, 60, 200 ng/mL. Grey points indicate cultures that were not induced with either IPTG or atc. Data are plotted on a symmetric log scale where the grey shaded region is linear. (**b**) 300 trajectories taken from the end of the MCMC process overlaid against the median (RFP,GFP) values from all three replicates.*

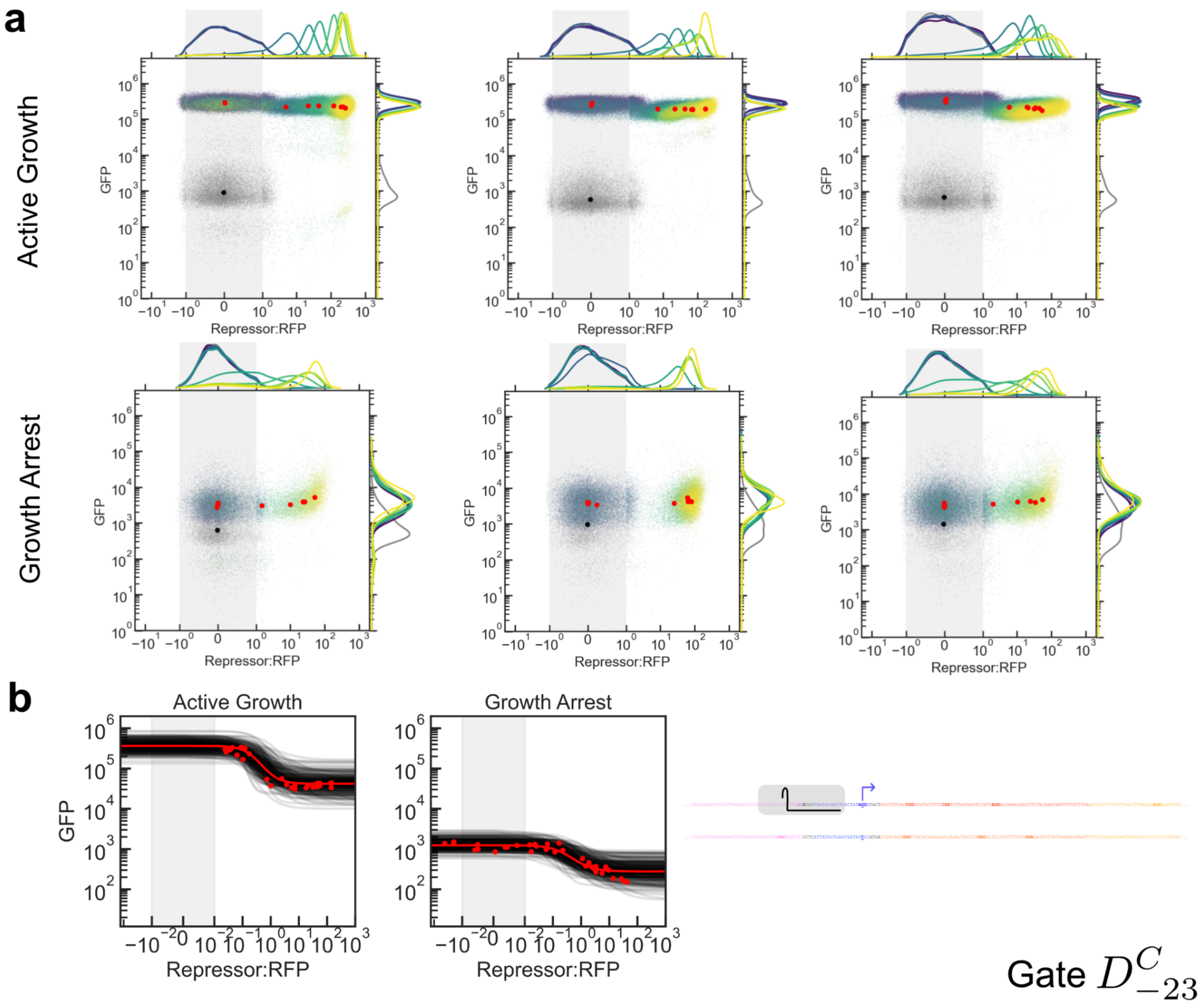


***Supplemental Fig. 2:*** *Repression curves for gate DC-23. (**a**) Flow cytometry data. Three biological replicates (columns) were measured on three different days. Dots represent medians of distributions for each induction condition, and colors represent atc concentrations. Induction conditions for atc were: Active Growth, Replicate 1: 0, 0.02, 0.06, 0.2, 0.43, 0.93, 2, 6, 20, 60, 200 ng/mL. Active Growth, Replicate 2: 0, 0.02, 0.2, 0.43, 0.93, 2, 3.5, 6.3, 11.3, 20, 200 ng/mL. Active Growth, Replicate 3: 0, 0.2, 0.43, 0.93, 2, 2.6, 3.5, 6.3, 11.3, 20, 200 ng/mL. Growth Arrest, all replicates: 0, 0.02, 0.06, 0.2, 0.43, 0.93, 2, 6, 20, 60, 200 ng/mL. Grey points indicate cultures that were not induced with either IPTG or atc. Data are plotted on a symmetric log scale where the grey shaded region is linear. (**b**) 300 trajectories taken from the end of the MCMC process overlaid against the median (RFP,GFP) values from all three replicates.*

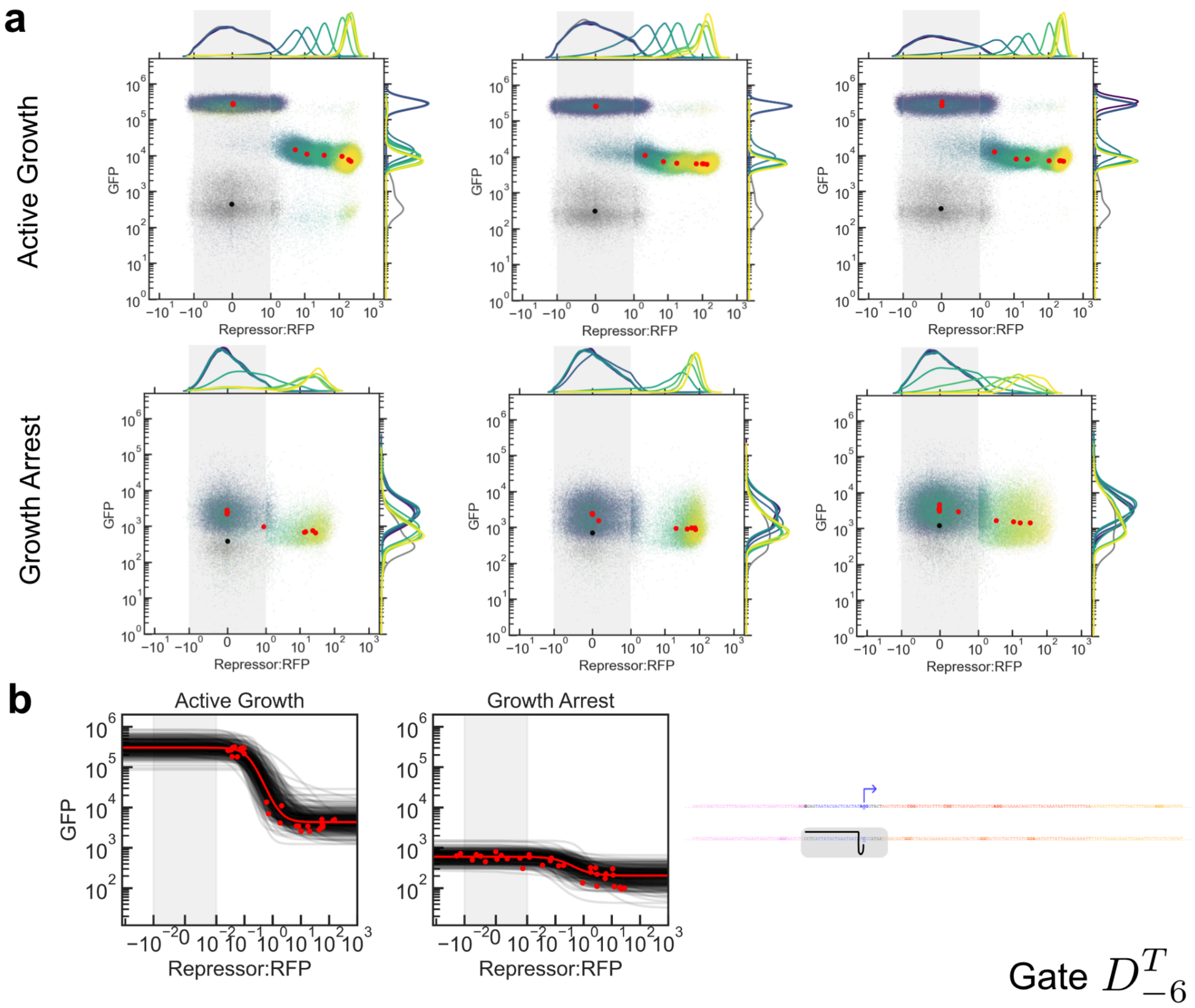


***Supplemental Fig. 3:*** *Repression curves for gate $D^T_{-6}$. (**a**) Flow cytometry data. Three biological replicates (columns) were measured on three different days. Dots represent medians of distributions for each induction condition, and colors represent atc concentrations. Induction conditions for atc were: Active Growth, Replicate 1: 0, 0.02, 0.06, 0.2, 0.43, 0.93, 2, 6, 20, 60, 200 ng/mL. Active Growth, Replicate 2: 0, 0.02, 0.2, 0.43, 0.93, 2, 3.5, 6.3, 11.3, 20, 200 ng/mL. Active Growth, Replicate 3: 0, 0.2, 0.43, 0.93, 2, 2.6, 3.5, 6.3, 11.3, 20, 200 ng/mL. Growth Arrest, all replicates: 0, 0.02, 0.06, 0.2, 0.43, 0.93, 2, 6, 20, 60, 200 ng/mL. Grey points indicate cultures that were not induced with either IPTG or atc. Data are plotted on a symmetric log scale where the grey shaded region is linear. (**b**) 300 trajectories taken from the end of the MCMC process overlaid against the median (RFP,GFP) values from all three replicates.*

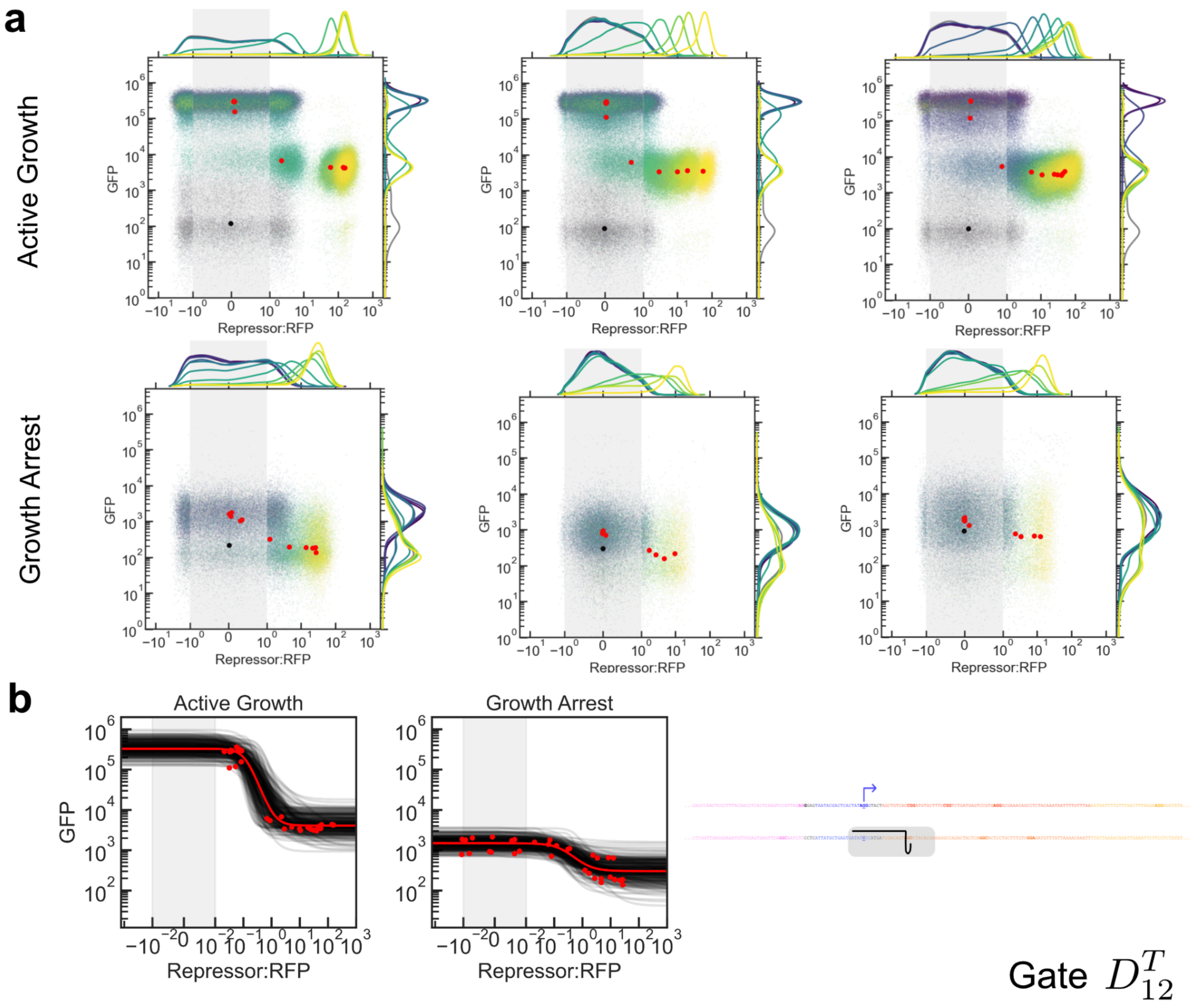


***Supplemental Fig. 4:*** *Repression curves for gate $D^{T}_{12}$. (**a**) Flow cytometry data. Three biological replicates (columns) were measured on three different days. Dots represent medians of distributions for each induction condition, and colors represent atc concentrations. Induction conditions for atc were: Active Growth, Replicate 1: 0, 0.02, 0.06, 0.2, 0.43, 0.93, 2, 6, 20, 60, 200 ng/mL. Active Growth, Replicate 2: 0, 0.02, 0.2, 0.43, 0.93, 2, 3.5, 6.3, 11.3, 20, 200 ng/mL. Active Growth, Replicate 3: 0, 0.2, 0.43, 0.93, 2, 2.6, 3.5, 6.3, 11.3, 20, 200 ng/mL. Growth Arrest, all replicates: 0, 0.02, 0.06, 0.2, 0.43, 0.93, 2, 6, 20, 60, 200 ng/mL. Grey points indicate cultures that were not induced with either IPTG or atc. Data are plotted on a symmetric log scale where the grey shaded region is linear. (**b**) 300 trajectories taken from the end of the MCMC process overlaid against the median (RFP,GFP) values from all three replicates.*

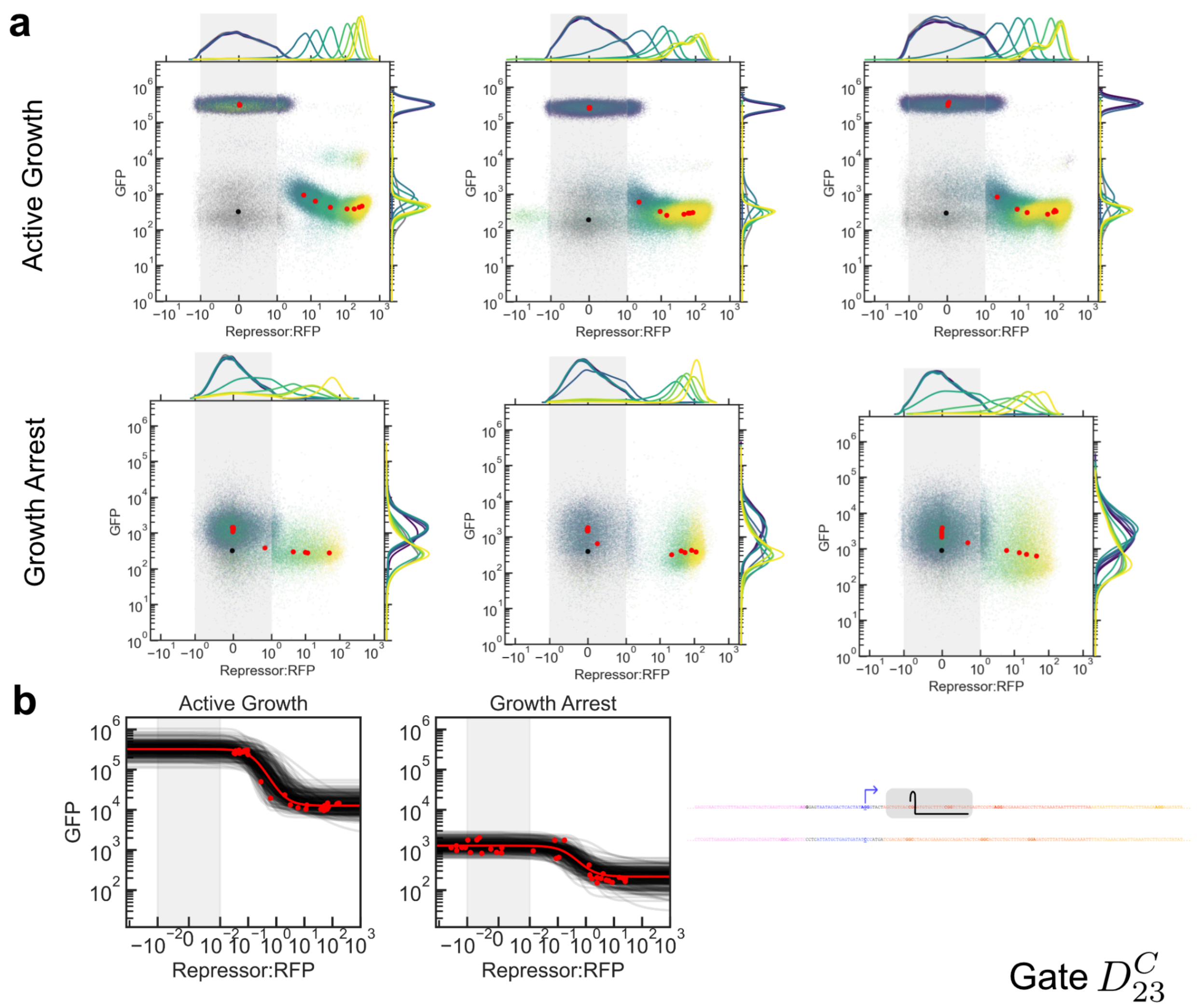


***Supplemental Fig. 5:*** *Repression curves for gate $D^C_{23}$. (**a**) Flow cytometry data. Three biological replicates (columns) were measured on three different days. Dots represent medians of distributions for each induction condition, and colors represent atc concentrations. Induction conditions for atc were: Active Growth, Replicate 1: 0, 0.02, 0.06, 0.2, 0.43, 0.93, 2, 6, 20, 60, 200 ng/mL. Active Growth, Replicate 2: 0, 0.02, 0.2, 0.43, 0.93, 2, 3.5, 6.3, 11.3, 20, 200 ng/mL. Active Growth, Replicate 3: 0, 0.2, 0.43, 0.93, 2, 2.6, 3.5, 6.3, 11.3, 20, 200 ng/mL. Growth Arrest, all replicates: 0, 0.02, 0.06, 0.2, 0.43, 0.93, 2, 6, 20, 60, 200 ng/mL. Grey points indicate cultures that were not induced with either IPTG or atc. Data are plotted on a symmetric log scale where the grey shaded region is linear. (**b**) 300 trajectories taken from the end of the MCMC process overlaid against the median (RFP,GFP) values from all three replicates.*

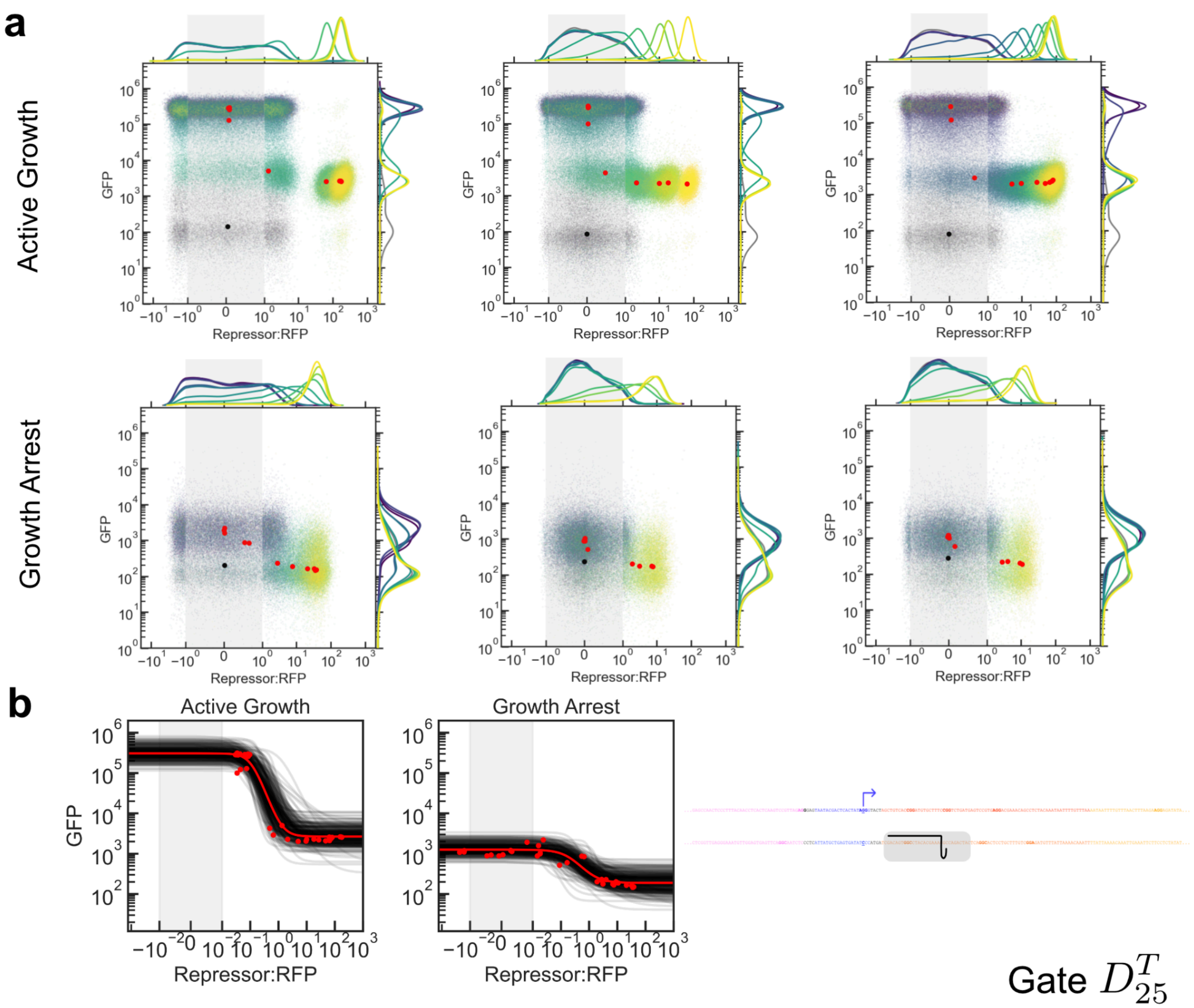


***Supplemental Fig. 6:*** *Repression curves for gate $D^T{}_{25}$. (**a**) Flow cytometry data. Three biological replicates (columns) were measured on three different days. Dots represent medians of distributions for each induction condition, and colors represent atc concentrations. Induction conditions for atc were: Active Growth, Replicate 1: 0, 0.02, 0.06, 0.2, 0.43, 0.93, 2, 6, 20, 60, 200 ng/mL. Active Growth, Replicate 2: 0, 0.02, 0.2, 0.43, 0.93, 2, 3.5, 6.3, 11.3, 20, 200 ng/mL. Active Growth, Replicate 3: 0, 0.2, 0.43, 0.93, 2, 2.6, 3.5, 6.3, 11.3, 20, 200 ng/mL. Growth Arrest, all replicates: 0, 0.02, 0.06, 0.2, 0.43, 0.93, 2, 6, 20, 60, 200 ng/mL. Grey points indicate cultures that were not induced with either IPTG or atc. Data are plotted on a symmetric log scale where the grey shaded region is linear. (**b**) 300 trajectories taken from the end of the MCMC process overlaid against the median (RFP,GFP) values from all three replicates.*

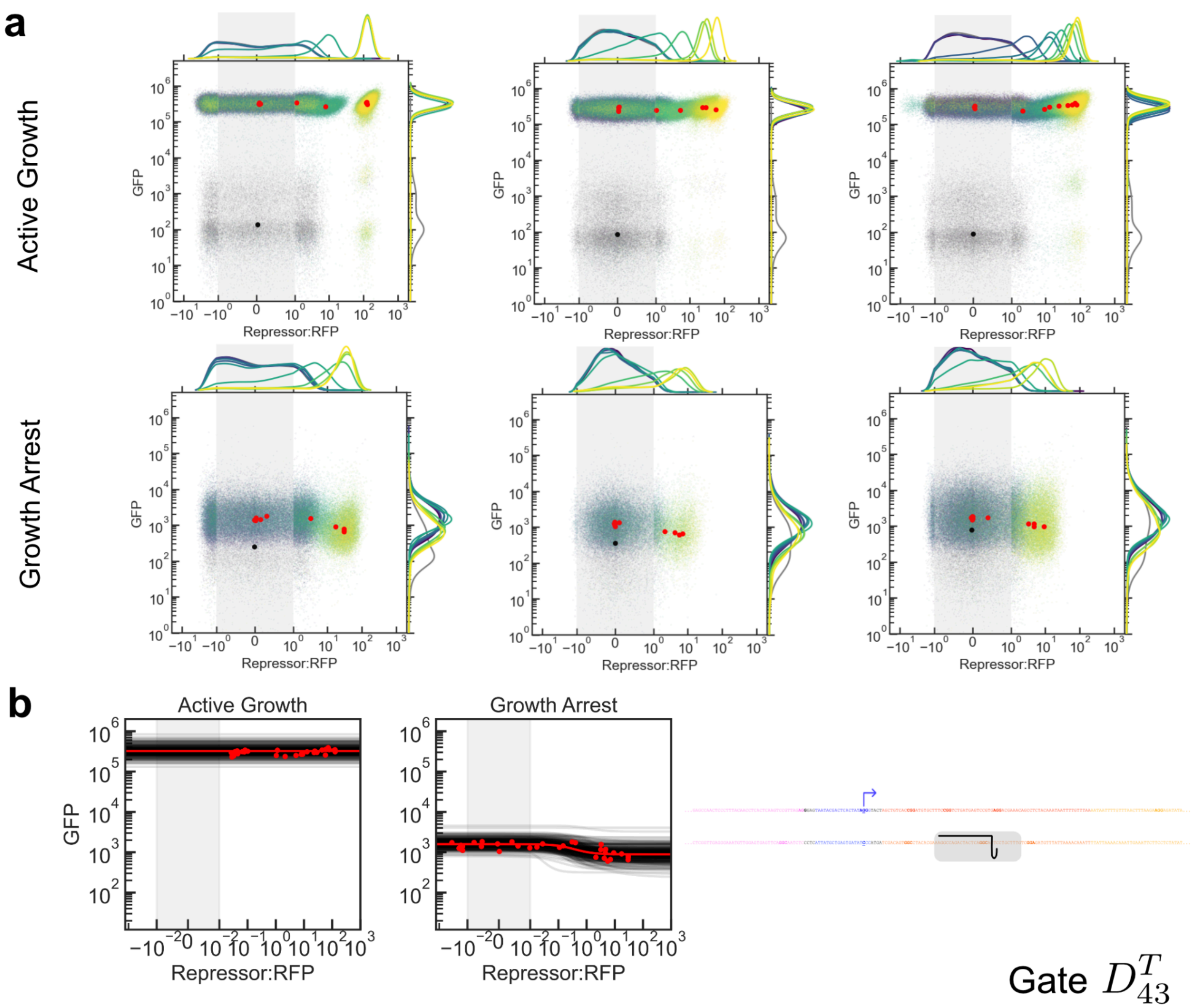


***Supplemental Fig. 7:*** *Repression curves for gate $D^T_{43}$. (**a**) Flow cytometry data. Three biological replicates (columns) were measured on three different days. Dots represent medians of distributions for each induction condition, and colors represent atc concentrations. Induction conditions for atc were: Active Growth, Replicate 1: 0, 0.02, 0.06, 0.2, 0.43, 0.93, 2, 6, 20, 60, 200 ng/mL. Active Growth, Replicate 2: 0, 0.02, 0.2, 0.43, 0.93, 2, 3.5, 6.3, 11.3, 20, 200 ng/mL. Active Growth, Replicate 3: 0, 0.2, 0.43, 0.93, 2, 2.6, 3.5, 6.3, 11.3, 20, 200 ng/mL. Growth Arrest, all replicates: 0, 0.02, 0.06, 0.2, 0.43, 0.93, 2, 6, 20, 60, 200 ng/mL. Grey points indicate cultures that were not induced with either IPTG or atc. Data are plotted on a symmetric log scale where the grey shaded region is linear. (**b**) 300 trajectories taken from the end of the MCMC process overlaid against the median (RFP,GFP) values from all three replicates.*

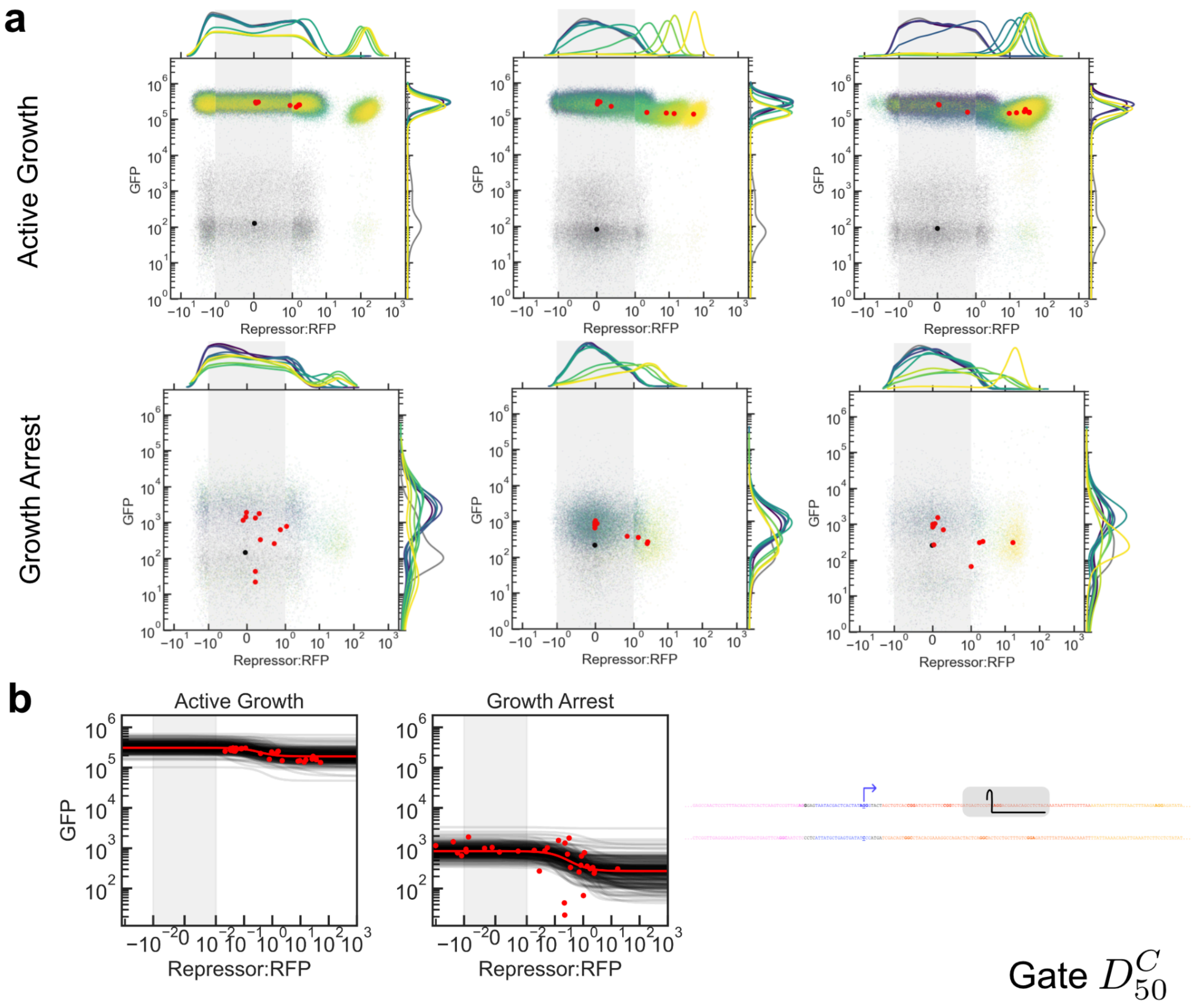


***Supplemental Fig. 8:*** *Repression curves for gate $D^C_{50}$. (**a**) Flow cytometry data. Three biological replicates (columns) were measured on three different days. Dots represent medians of distributions for each induction condition, and colors represent atc concentrations. Induction conditions for atc were: Active Growth, Replicate 1: 0, 0.02, 0.06, 0.2, 0.43, 0.93, 2, 6, 20, 60, 200 ng/mL. Active Growth, Replicate 2: 0, 0.02, 0.2, 0.43, 0.93, 2, 3.5, 6.3, 11.3, 20, 200 ng/mL. Active Growth, Replicate 3: 0, 0.2, 0.43, 0.93, 2, 2.6, 3.5, 6.3, 11.3, 20, 200 ng/mL. Growth Arrest, all replicates: 0, 0.02, 0.06, 0.2, 0.43, 0.93, 2, 6, 20, 60, 200 ng/mL. Grey points indicate cultures that were not induced with either IPTG or atc. Data are plotted on a symmetric log scale where the grey shaded region is linear. (**b**) 300 trajectories taken from the end of the MCMC process overlaid against the median (RFP,GFP) values from all three replicates.*

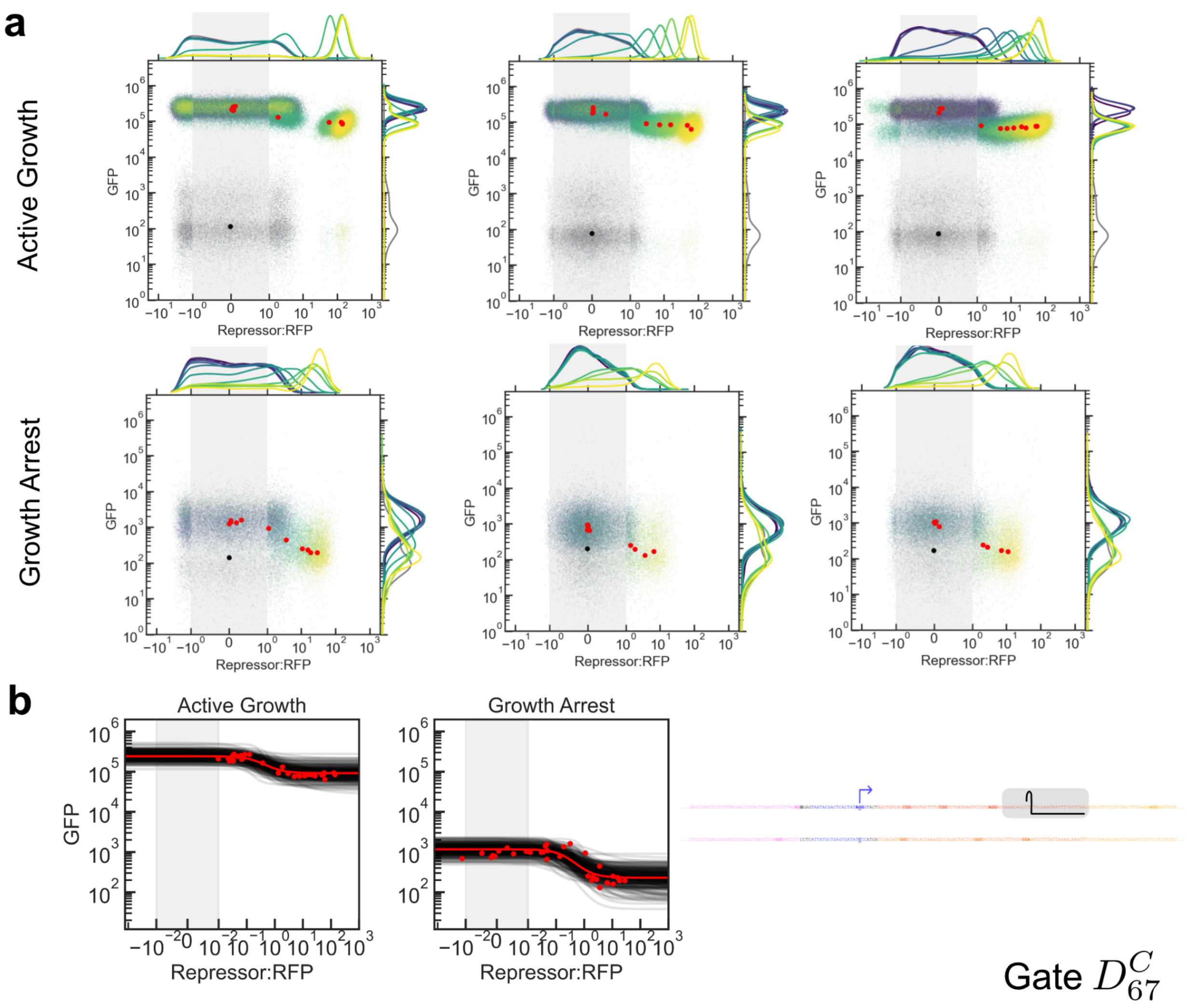


***Supplemental Fig. 9:*** *Repression curves for gate $D^{C}_{67}$. (**a**) Flow cytometry data. Three biological replicates (columns) were measured on three different days. Dots represent medians of distributions for each induction condition, and colors represent atc concentrations. Induction conditions for atc were: Active Growth, Replicate 1: 0, 0.02, 0.06, 0.2, 0.43, 0.93, 2, 6, 20, 60, 200 ng/mL. Active Growth, Replicate 2: 0, 0.02, 0.2, 0.43, 0.93, 2, 3.5, 6.3, 11.3, 20, 200 ng/mL. Active Growth, Replicate 3: 0, 0.2, 0.43, 0.93, 2, 2.6, 3.5, 6.3, 11.3, 20, 200 ng/mL. Growth Arrest, all replicates: 0, 0.02, 0.06, 0.2, 0.43, 0.93, 2, 6, 20, 60, 200 ng/mL. Grey points indicate cultures that were not induced with either IPTG or atc. Data are plotted on a symmetric log scale where the grey shaded region is linear. (**b**) 300 trajectories taken from the end of the MCMC process overlaid against the median (RFP,GFP) values from all three replicates.*

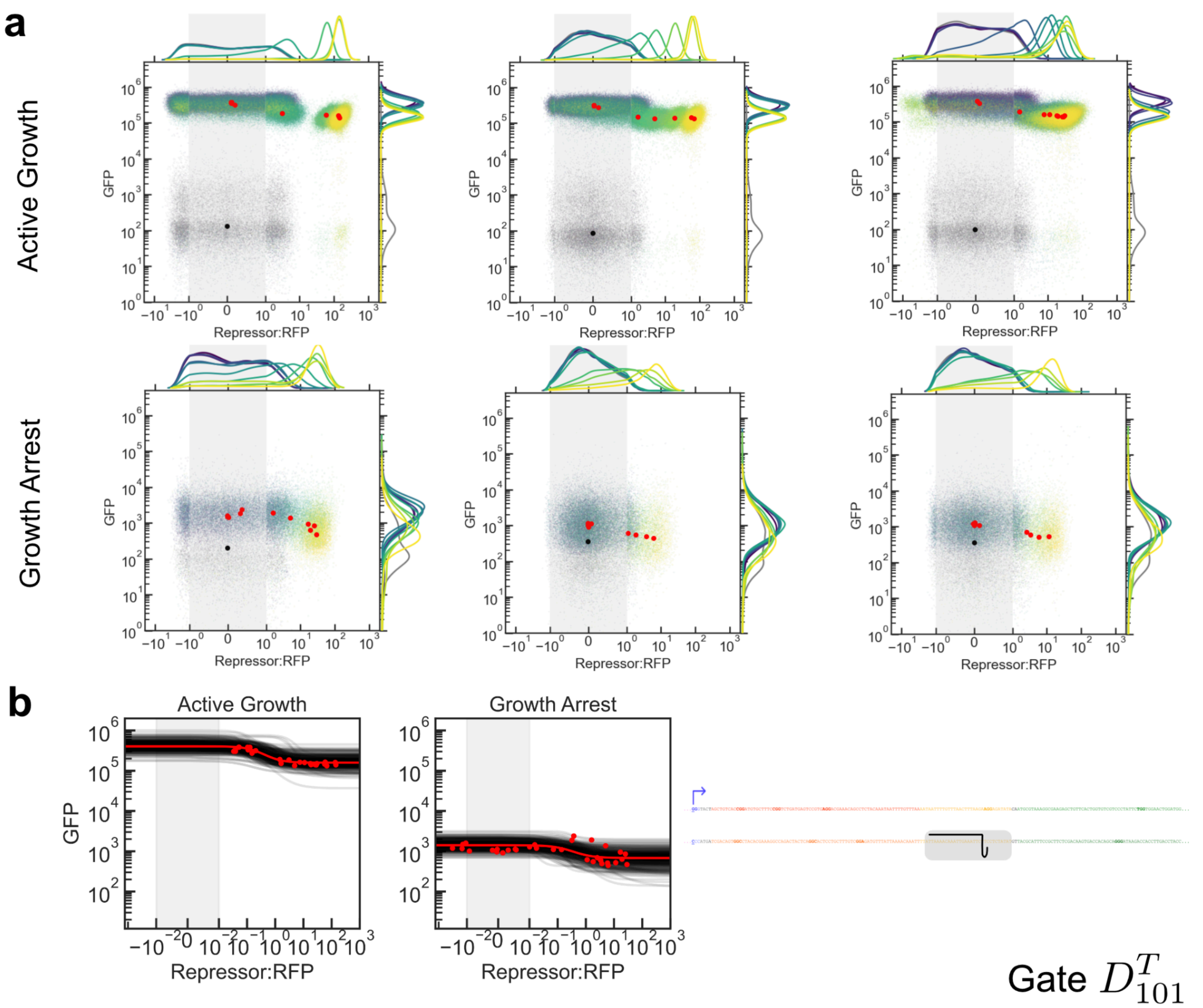


***Supplemental Fig. 10:*** *Repression curves for gate* $D^T_{101}$*. (**a**) Flow cytometry data. Three biological replicates (columns) were measured on three different days. Dots represent medians of distributions for each induction condition, and colors represent atc concentrations. Induction conditions for atc were: Active Growth, Replicate 1: 0, 0.02, 0.06, 0.2, 0.43, 0.93, 2, 6, 20, 60, 200 ng/mL. Active Growth, Replicate 2: 0, 0.02, 0.2, 0.43, 0.93, 2, 3.5, 6.3, 11.3, 20, 200 ng/mL. Active Growth, Replicate 3: 0, 0.2, 0.43, 0.93, 2, 2.6, 3.5, 6.3, 11.3, 20, 200 ng/mL. Growth Arrest, all replicates: 0, 0.02, 0.06, 0.2, 0.43, 0.93, 2, 6, 20, 60, 200 ng/mL. Grey points indicate cultures that were not induced with either IPTG or atc. Data are plotted on a symmetric log scale where the grey shaded region is linear. (**b**) 300 trajectories taken from the end of the MCMC process overlaid against the median (RFP,GFP) values from all three replicates.*

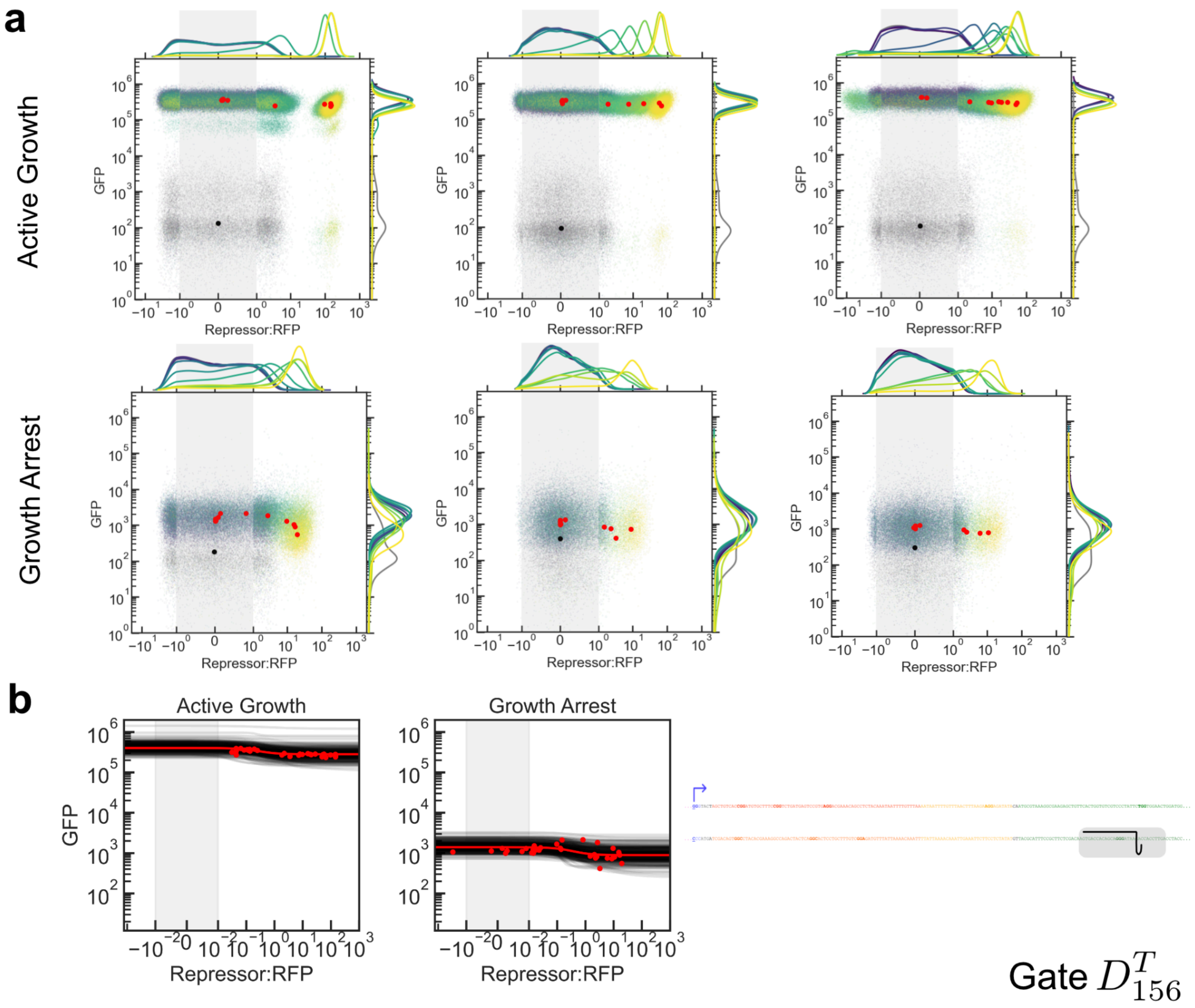


***Supplemental Fig. 11:*** *Repression curves for gate $D^T_{156}$. (**a**) Flow cytometry data. Three biological replicates (columns) were measured on three different days. Dots represent medians of distributions for each induction condition, and colors represent atc concentrations. Induction conditions for atc were: Active Growth, Replicate 1: 0, 0.02, 0.06, 0.2, 0.43, 0.93, 2, 6, 20, 60, 200 ng/mL. Active Growth, Replicate 2: 0, 0.02, 0.2, 0.43, 0.93, 2, 3.5, 6.3, 11.3, 20, 200 ng/mL. Active Growth, Replicate 3: 0, 0.2, 0.43, 0.93, 2, 2.6, 3.5, 6.3, 11.3, 20, 200 ng/mL. Growth Arrest, all replicates: 0, 0.02, 0.06, 0.2, 0.43, 0.93, 2, 6, 20, 60, 200 ng/mL. Grey points indicate cultures that were not induced with either IPTG or atc. Data are plotted on a symmetric log scale where the grey shaded region is linear. (**b**) 300 trajectories taken from the end of the MCMC process overlaid against the median (RFP,GFP) values from all three replicates.*

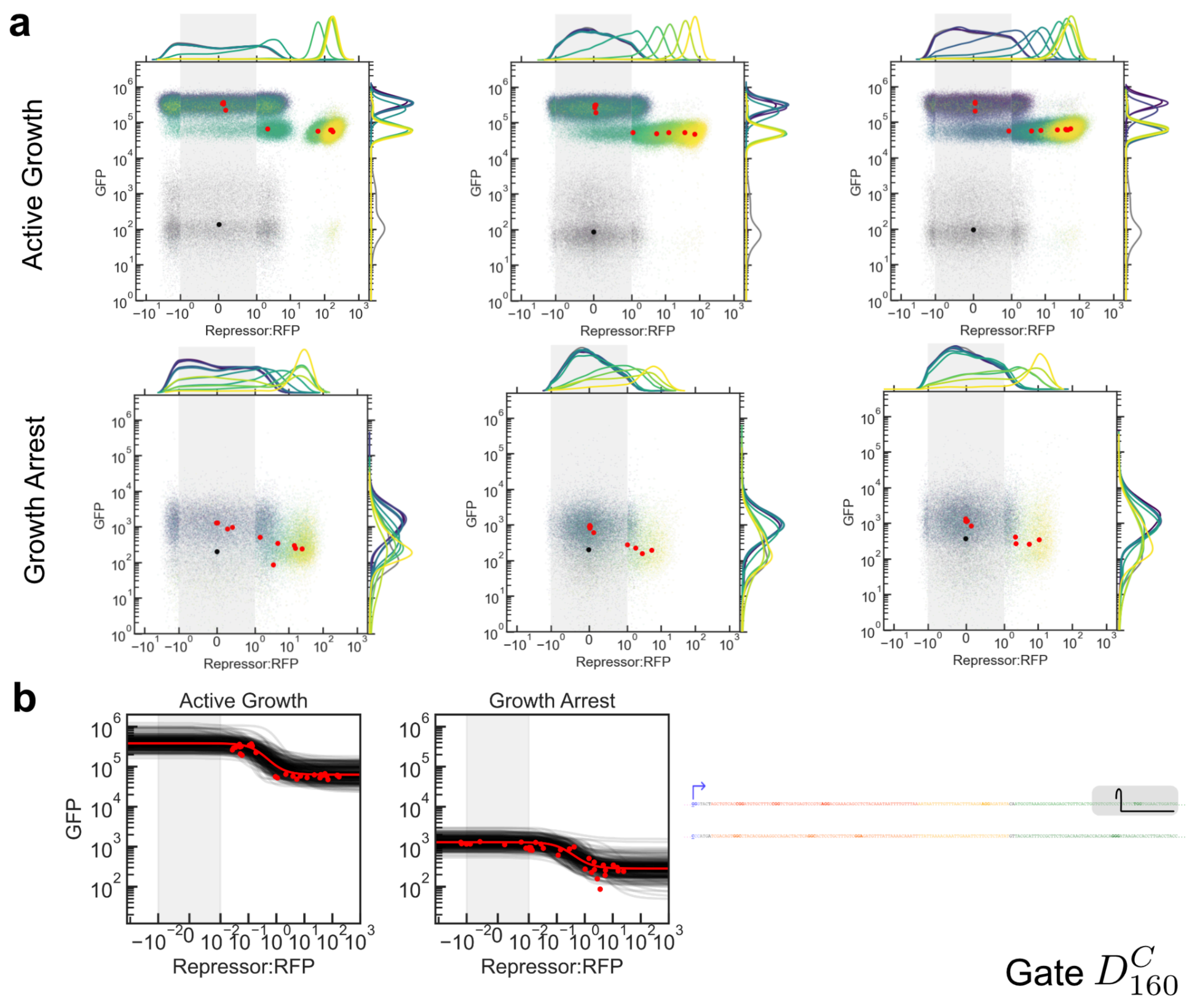


***Supplemental Fig. 12:*** *Repression curves for gate $D^{C}_{160}$. (**a**) Flow cytometry data. Three biological replicates (columns) were measured on three different days. Dots represent medians of distributions for each induction condition, and colors represent atc concentrations. Induction conditions for atc were: Active Growth, Replicate 1: 0, 0.02, 0.06, 0.2, 0.43, 0.93, 2, 6, 20, 60, 200 ng/mL. Active Growth, Replicate 2: 0, 0.02, 0.2, 0.43, 0.93, 2, 3.5, 6.3, 11.3, 20, 200 ng/mL. Active Growth, Replicate 3: 0, 0.2, 0.43, 0.93, 2, 2.6, 3.5, 6.3, 11.3, 20, 200 ng/mL. Growth Arrest, all replicates: 0, 0.02, 0.06, 0.2, 0.43, 0.93, 2, 6, 20, 60, 200 ng/mL. Grey points indicate cultures that were not induced with either IPTG or atc. Data are plotted on a symmetric log scale where the grey shaded region is linear. (**b**) 300 trajectories taken from the end of the MCMC process overlaid against the median (RFP,GFP) values from all three replicates.*

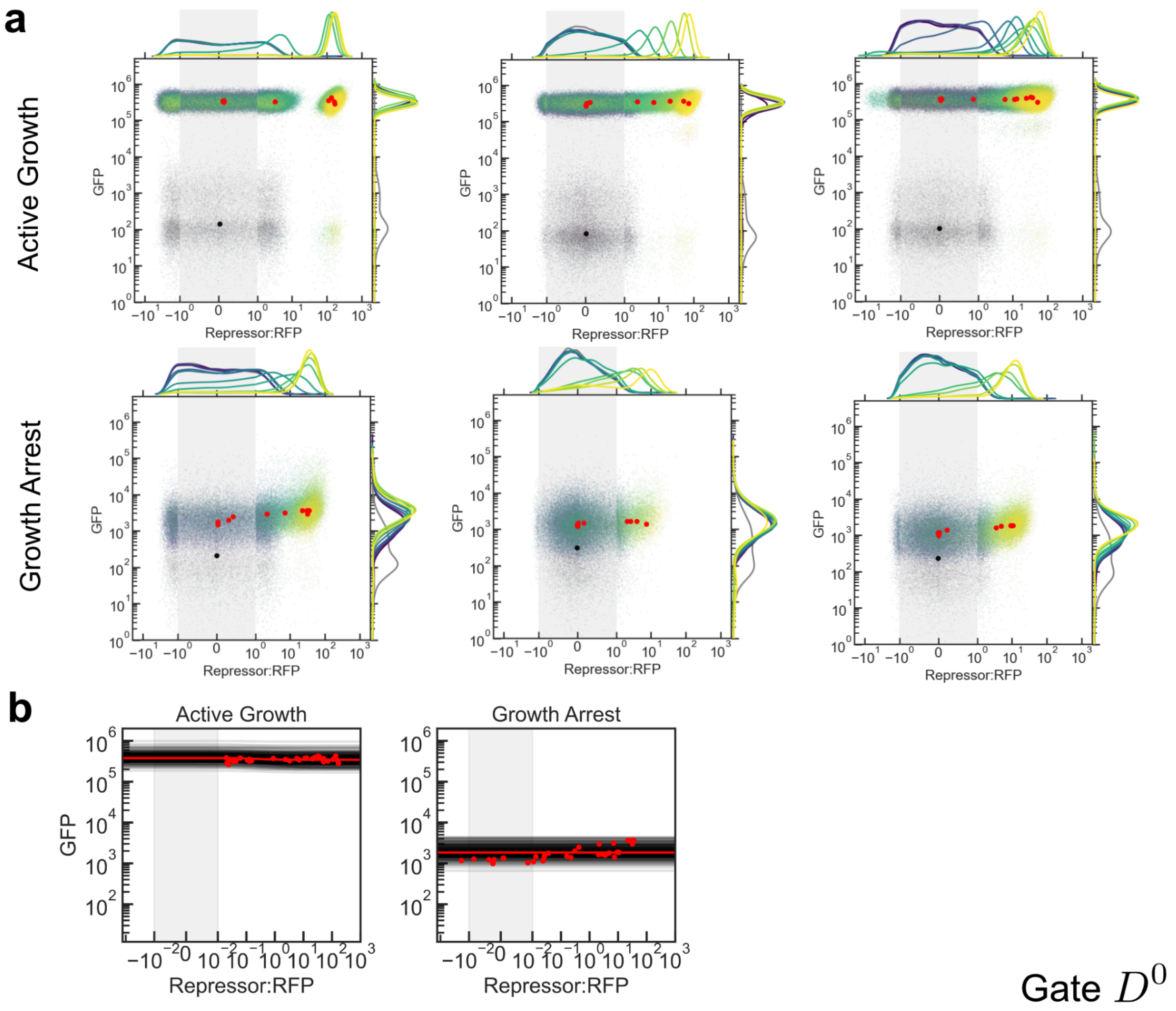


***Supplemental Fig. 13:*** *Repression curves for the negative control gate $D^0$. (**a**) Flow cytometry data. Three biological replicates (columns) were measured on three different days. Dots represent medians of distributions for each induction condition, and colors represent atc concentrations. Induction conditions for atc were: Active Growth, Replicate 1: 0, 0.02, 0.06, 0.2, 0.43, 0.93, 2, 6, 20, 60, 200 ng/mL. Active Growth, Replicate 2: 0, 0.02, 0.2, 0.43, 0.93, 2, 3.5, 6.3, 11.3, 20, 200 ng/mL. Active Growth, Replicate 3: 0, 0.2, 0.43, 0.93, 2, 2.6, 3.5, 6.3, 11.3, 20, 200 ng/mL. Growth Arrest, all replicates: 0, 0.02, 0.06, 0.2, 0.43, 0.93, 2, 6, 20, 60, 200 ng/mL. Grey points indicate cultures that were not induced with either IPTG or atc. Data are plotted on a symmetric log scale where the grey shaded region is linear. (**b**) 300 trajectories taken from the end of the MCMC process overlaid against the median (RFP,GFP) values from all three replicates.*

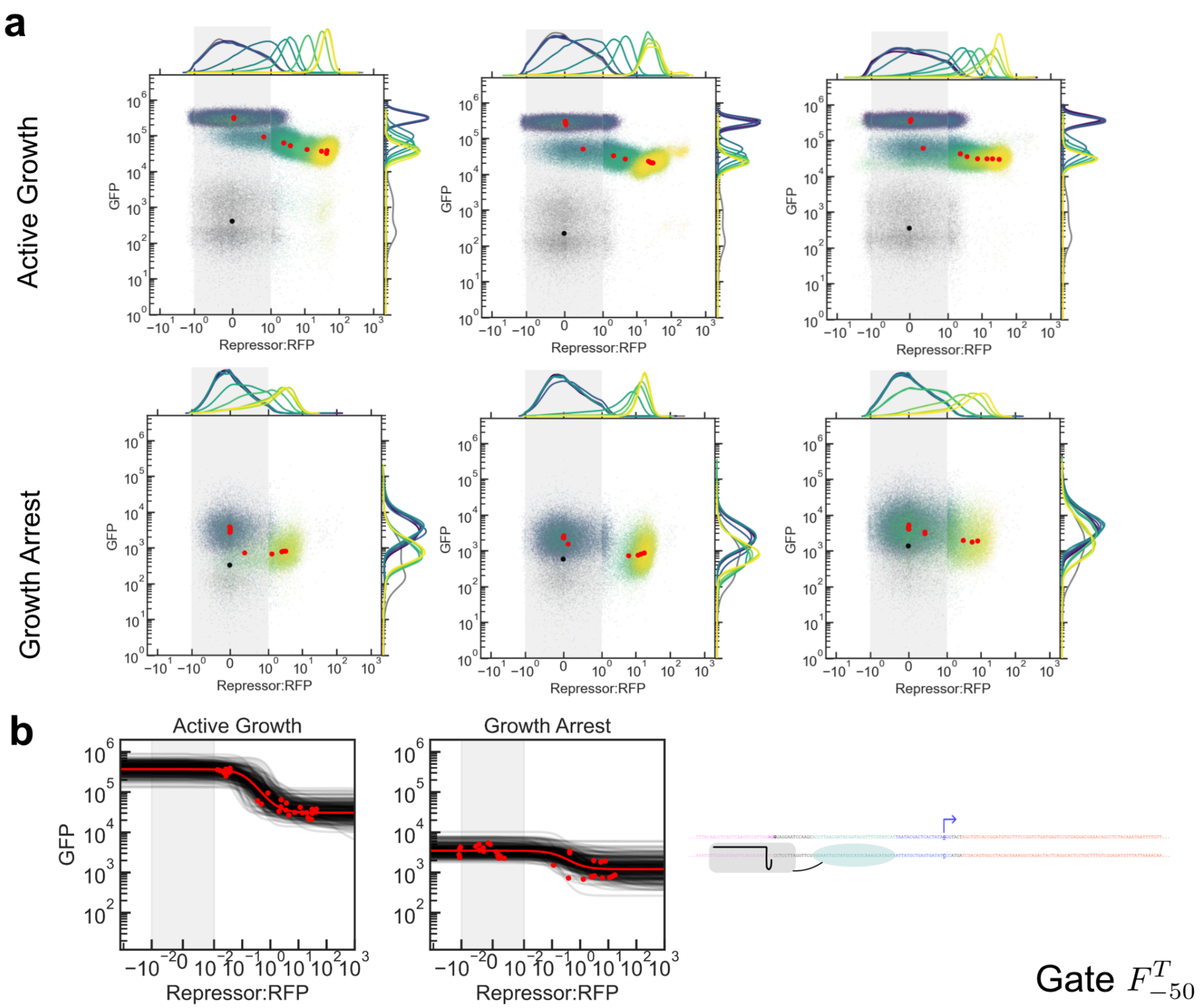


***Supplemental Fig. 14:*** *Repression curves for gate $F^{T}_{-50}$. (**a**) Flow cytometry data. Three biological replicates (columns) were measured on three different days. Dots represent medians of distributions for each induction condition, and colors represent atc concentrations. Induction conditions for atc were: Active Growth, all replicates: 0, 0.2, 0.43, 0.93, 2, 2.6, 3.5, 6.3, 11.3, 20, 100 ng/mL. Growth Arrest, all replicates: 0, 0.02, 0.06, 0.2, 0.43, 0.93, 2, 6, 20, 60, 200 ng/mL. Grey points indicate cultures that were not induced with either IPTG or atc. Data are plotted on a symmetric log scale where the grey shaded region is linear. (**b**) 300 trajectories taken from the end of the MCMC process overlaid against the median (RFP,GFP) values from all three replicates.*

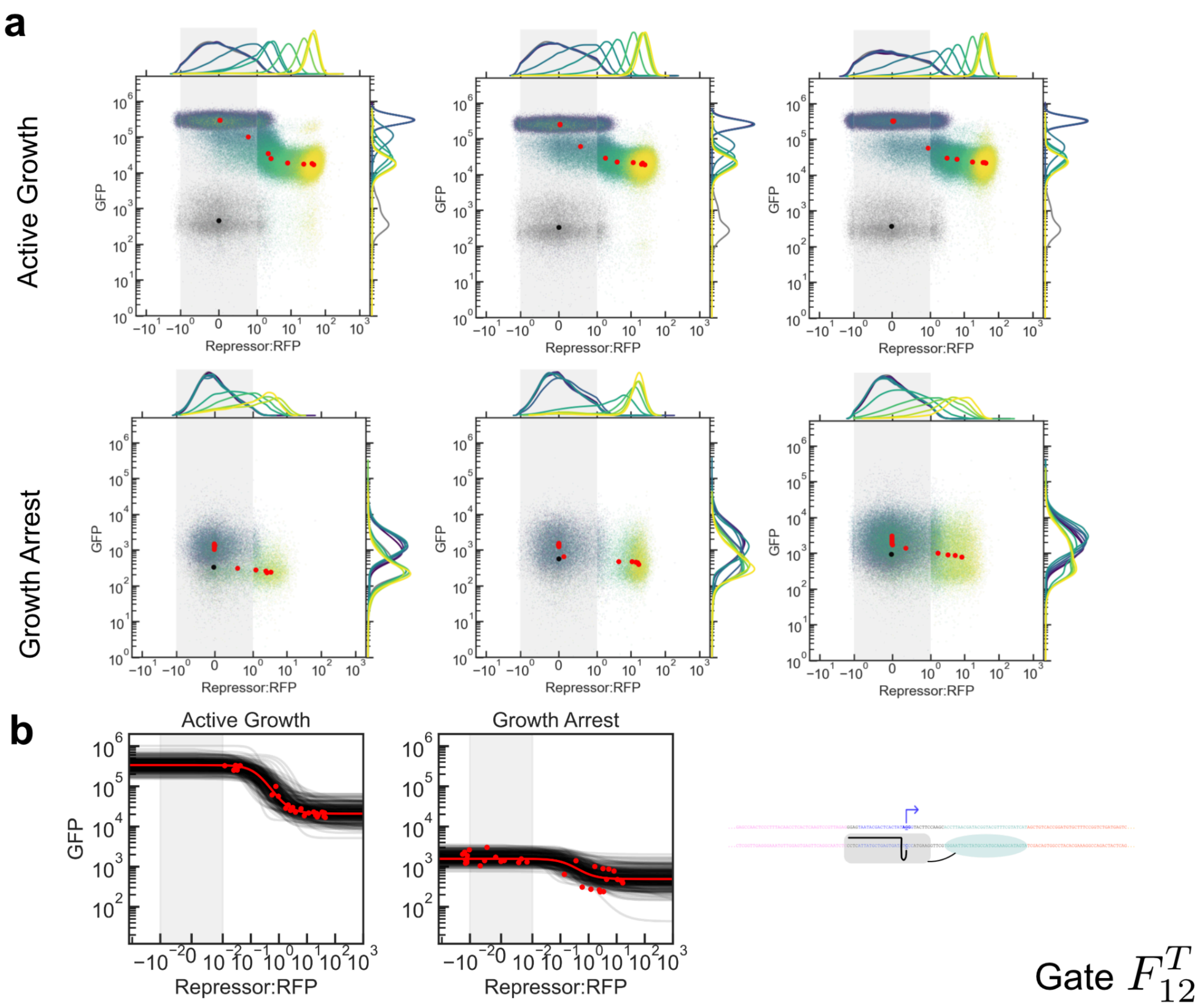


***Supplemental Fig. 15:*** *Repression curves for gate $F^{T}_{12}$. (**a**) Flow cytometry data. Three biological replicates (columns) were measured on three different days. Dots represent medians of distributions for each induction condition, and colors represent atc concentrations. Induction conditions for atc were: Active Growth, all replicates: 0, 0.2, 0.43, 0.93, 2, 2.6, 3.5, 6.3, 11.3, 20, 100 ng/mL. Growth Arrest, all replicates: 0, 0.02, 0.06, 0.2, 0.43, 0.93, 2, 6, 20, 60, 200 ng/mL. Grey points indicate cultures that were not induced with either IPTG or atc. Data are plotted on a symmetric log scale where the grey shaded region is linear. (**b**) 300 trajectories taken from the end of the MCMC process overlaid against the median (RFP,GFP) values from all three replicates.*

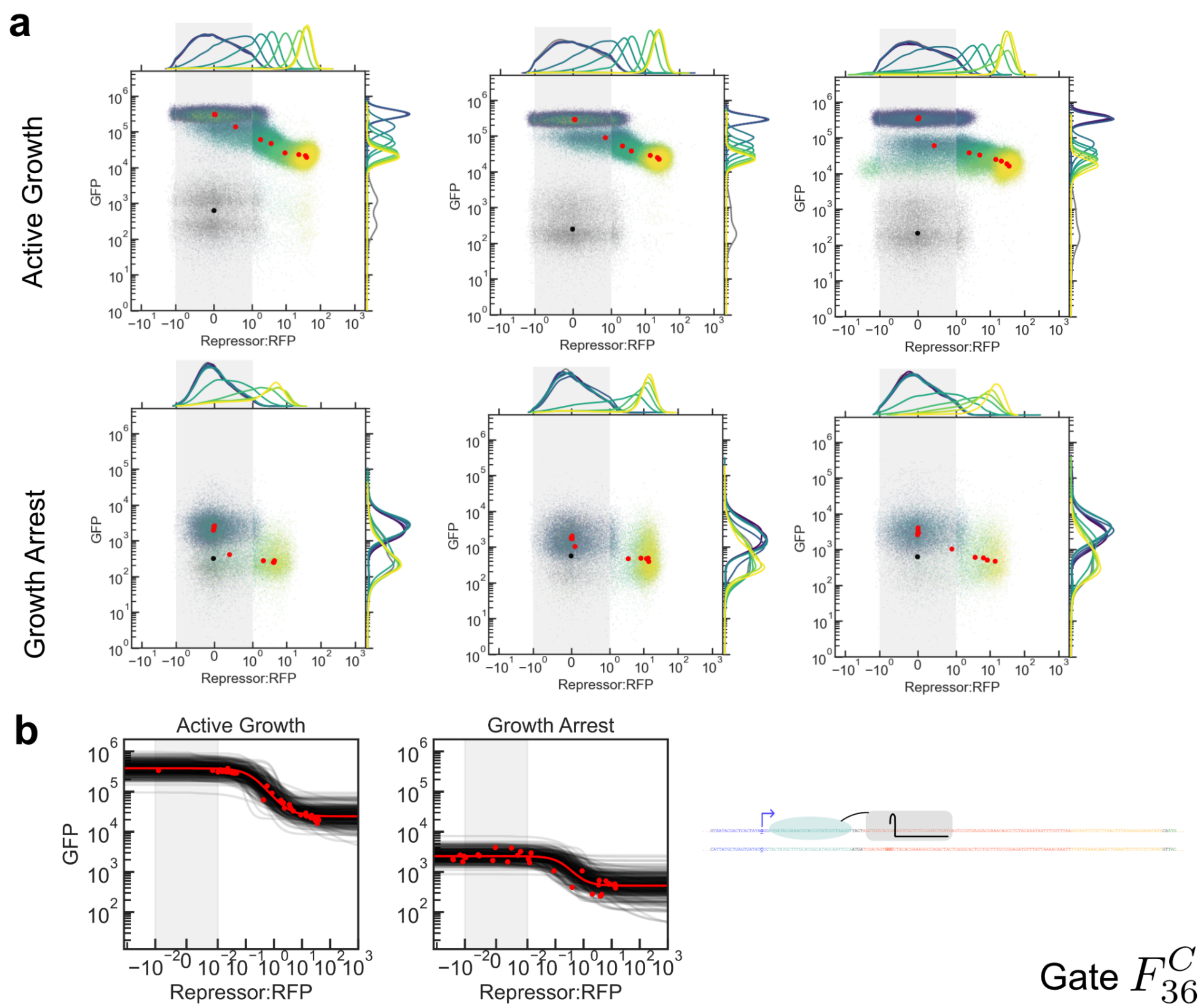


***Supplemental Fig. 16:*** *Repression curves for gate $F^{C}_{36}$. (**a**) Flow cytometry data. Three biological replicates (columns) were measured on three different days. Dots represent medians of distributions for each induction condition, and colors represent atc concentrations. Induction conditions for atc were: Active Growth, all replicates: 0, 0.2, 0.43, 0.93, 2, 2.6, 3.5, 6.3, 11.3, 20, 100 ng/mL. Growth Arrest, all replicates: 0, 0.02, 0.06, 0.2, 0.43, 0.93, 2, 6, 20, 60, 200 ng/mL. Grey points indicate cultures that were not induced with either IPTG or atc. Data are plotted on a symmetric log scale where the grey shaded region is linear. (**b**) 300 trajectories taken from the end of the MCMC process overlaid against the median (RFP,GFP) values from all three replicates.*

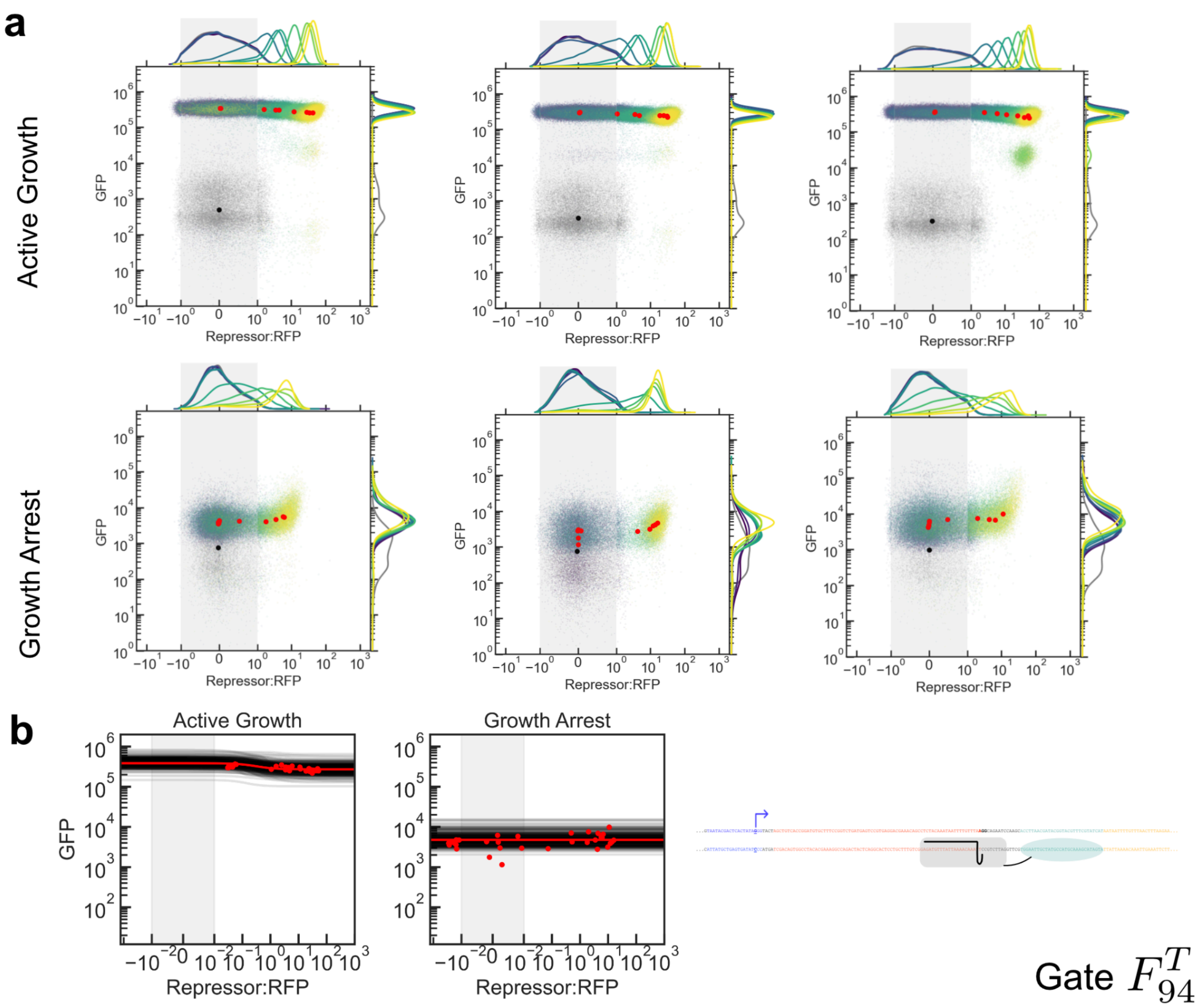


***Supplemental Fig. 17:*** *Repression curves for gate $F^{T}_{94}$. (**a**) Flow cytometry data. Three biological replicates (columns) were measured on three different days. Dots represent medians of distributions for each induction condition, and colors represent atc concentrations. Induction conditions for atc were: Active Growth, all replicates: 0, 0.2, 0.43, 0.93, 2, 2.6, 3.5, 6.3, 11.3, 20, 100 ng/mL. Growth Arrest, all replicates: 0, 0.02, 0.06, 0.2, 0.43, 0.93, 2, 6, 20, 60, 200 ng/mL. Grey points indicate cultures that were not induced with either IPTG or atc. Data are plotted on a symmetric log scale where the grey shaded region is linear. (**b**) 300 trajectories taken from the end of the MCMC process overlaid against the median (RFP,GFP) values from all three replicates.*

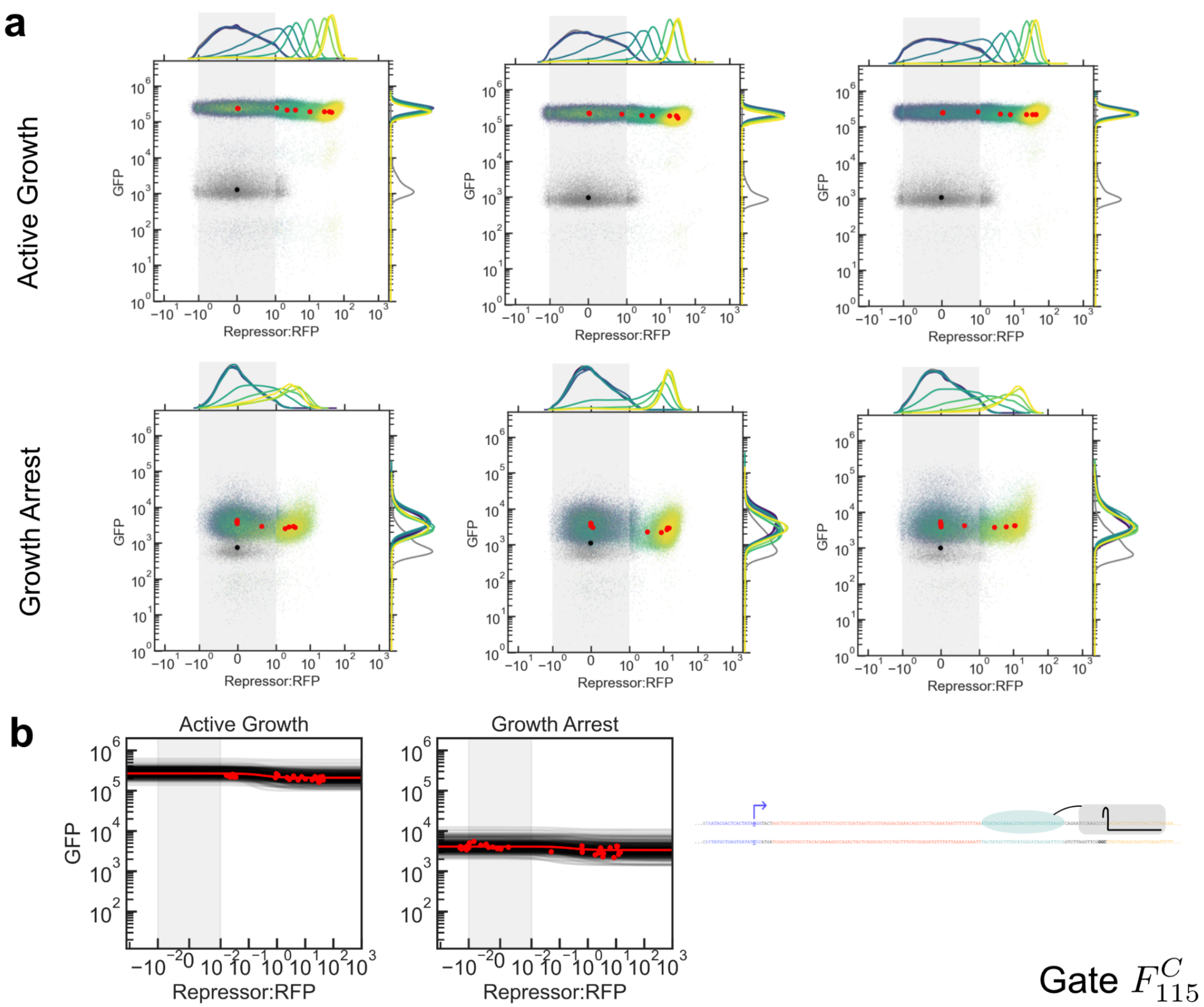


***Supplemental Fig. 18:*** *Repression curves for gate $F^{C}_{115}$. (**a**) Flow cytometry data. Three biological replicates (columns) were measured on three different days. Dots represent medians of distributions for each induction condition, and colors represent atc concentrations. Induction conditions for atc were: Active Growth, all replicates: 0, 0.2, 0.43, 0.93, 2, 2.6, 3.5, 6.3, 11.3, 20, 100 ng/mL. Growth Arrest, all replicates: 0, 0.02, 0.06, 0.2, 0.43, 0.93, 2, 6, 20, 60, 200 ng/mL. Grey points indicate cultures that were not induced with either IPTG or atc. Data are plotted on a symmetric log scale where the grey shaded region is linear. (**b**) 300 trajectories taken from the end of the MCMC process overlaid against the median (RFP,GFP) values from all three replicates.*

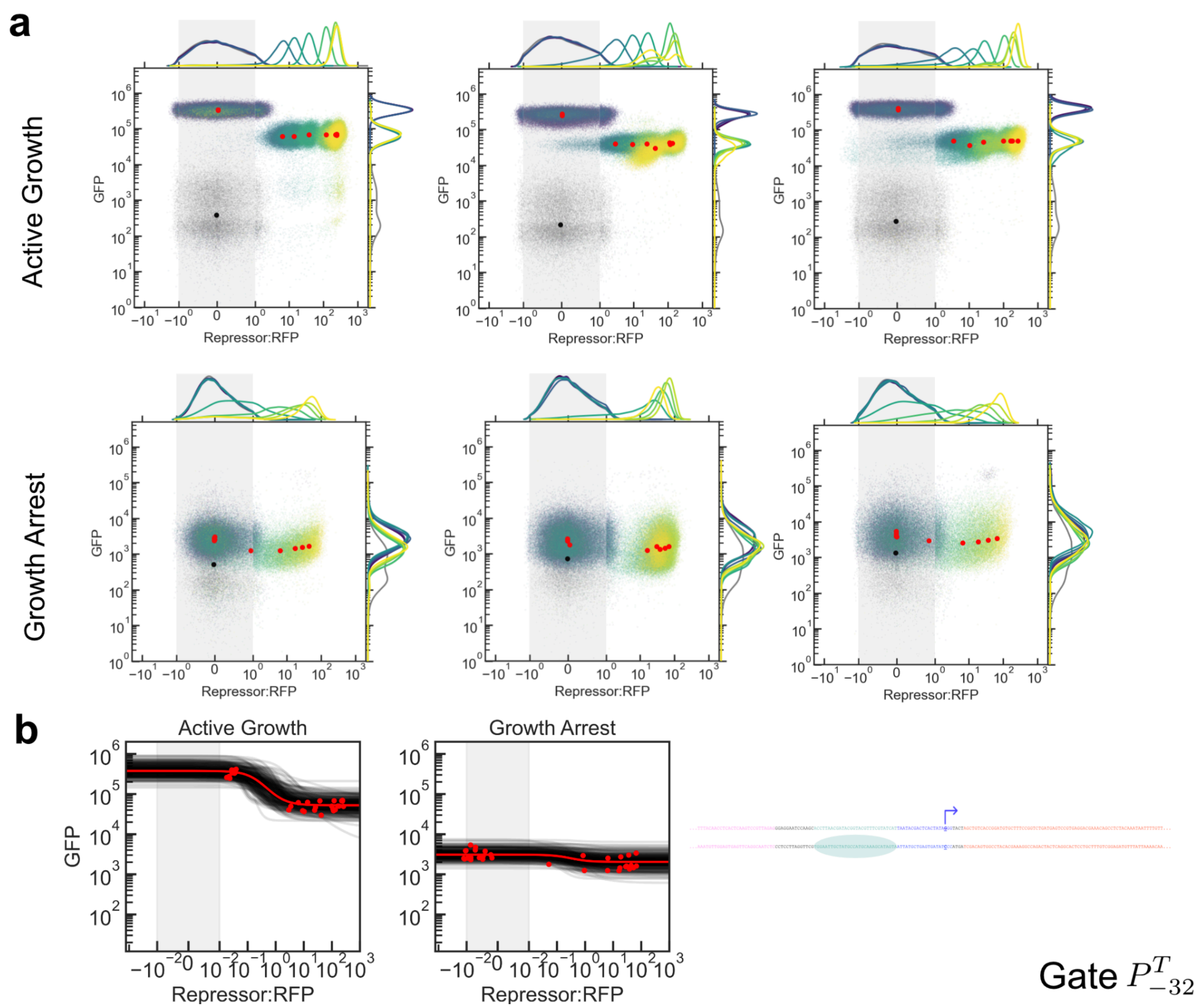


***Supplemental Fig. 19:*** *Repression curves for gate $P^{T}_{-32}$. (**a**) Flow cytometry data. Three biological replicates (columns) were measured on three different days. Dots represent medians of distributions for each induction condition, and colors represent atc concentrations. Induction conditions for atc were: Active Growth, all replicates: 0, 0.2, 0.43, 0.93, 2, 2.6, 3.5, 6.3, 11.3, 20, 100 ng/mL. Growth Arrest, all replicates: 0, 0.02, 0.06, 0.2, 0.43, 0.93, 2, 6, 20, 60, 200 ng/mL. Grey points indicate cultures that were not induced with either IPTG or atc. Data are plotted on a symmetric log scale where the grey shaded region is linear. (**b**) 300 trajectories taken from the end of the MCMC process overlaid against the median (RFP,GFP) values from all three replicates.*

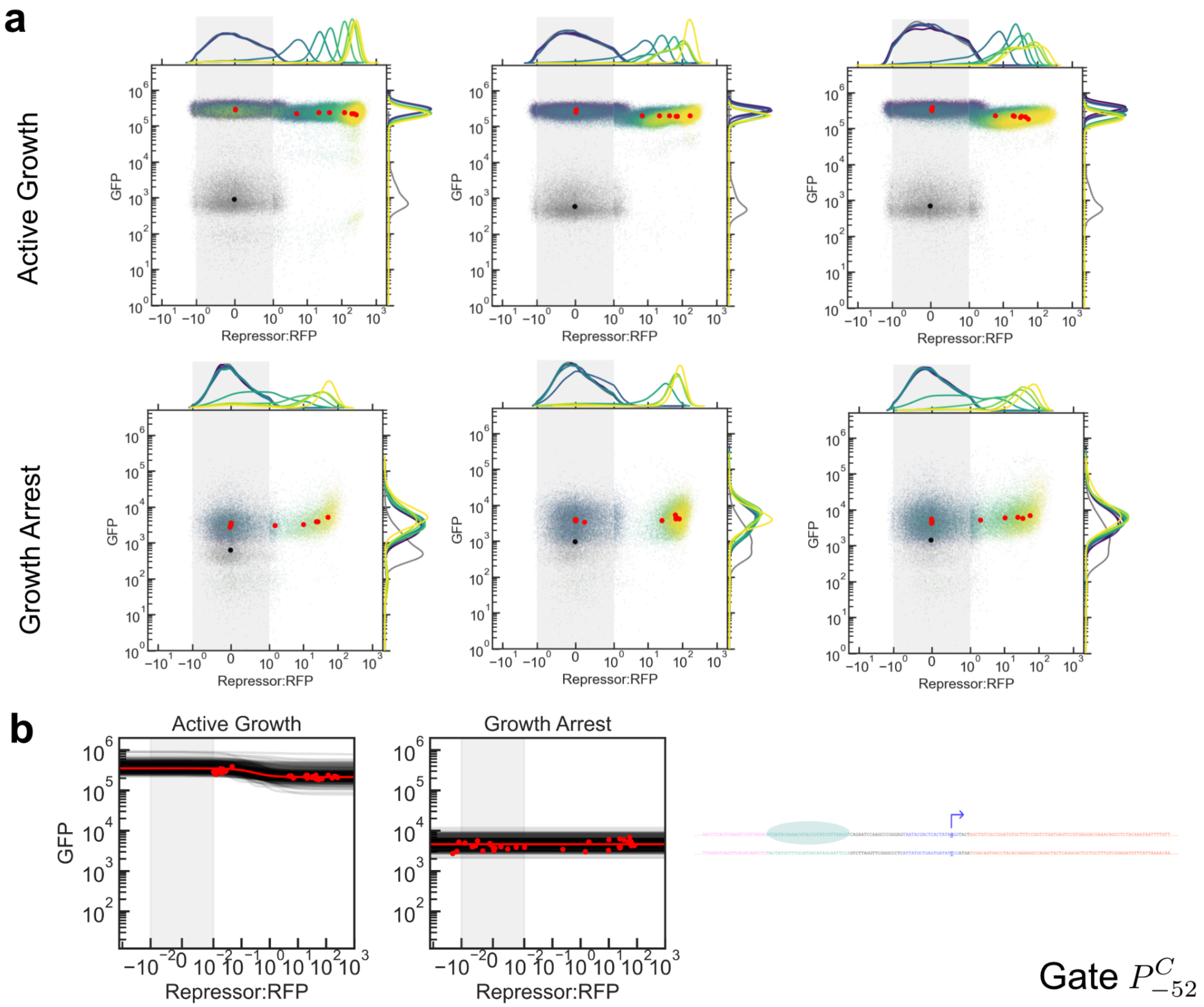


***Supplemental Fig. 20:*** *Repression curves for gate $P^{C}_{-52}$. (**a**) Flow cytometry data. Three biological replicates (columns) were measured on three different days. Dots represent medians of distributions for each induction condition, and colors represent atc concentrations. Induction conditions for atc were: Active Growth, all replicates: 0, 0.2, 0.43, 0.93, 2, 2.6, 3.5, 6.3, 11.3, 20, 100 ng/mL. Growth Arrest, all replicates: 0, 0.02, 0.06, 0.2, 0.43, 0.93, 2, 6, 20, 60, 200 ng/mL. Grey points indicate cultures that were not induced with either IPTG or atc. Data are plotted on a symmetric log scale where the grey shaded region is linear. (**b**) 300 trajectories taken from the end of the MCMC process overlaid against the median (RFP,GFP) values from all three replicates.*

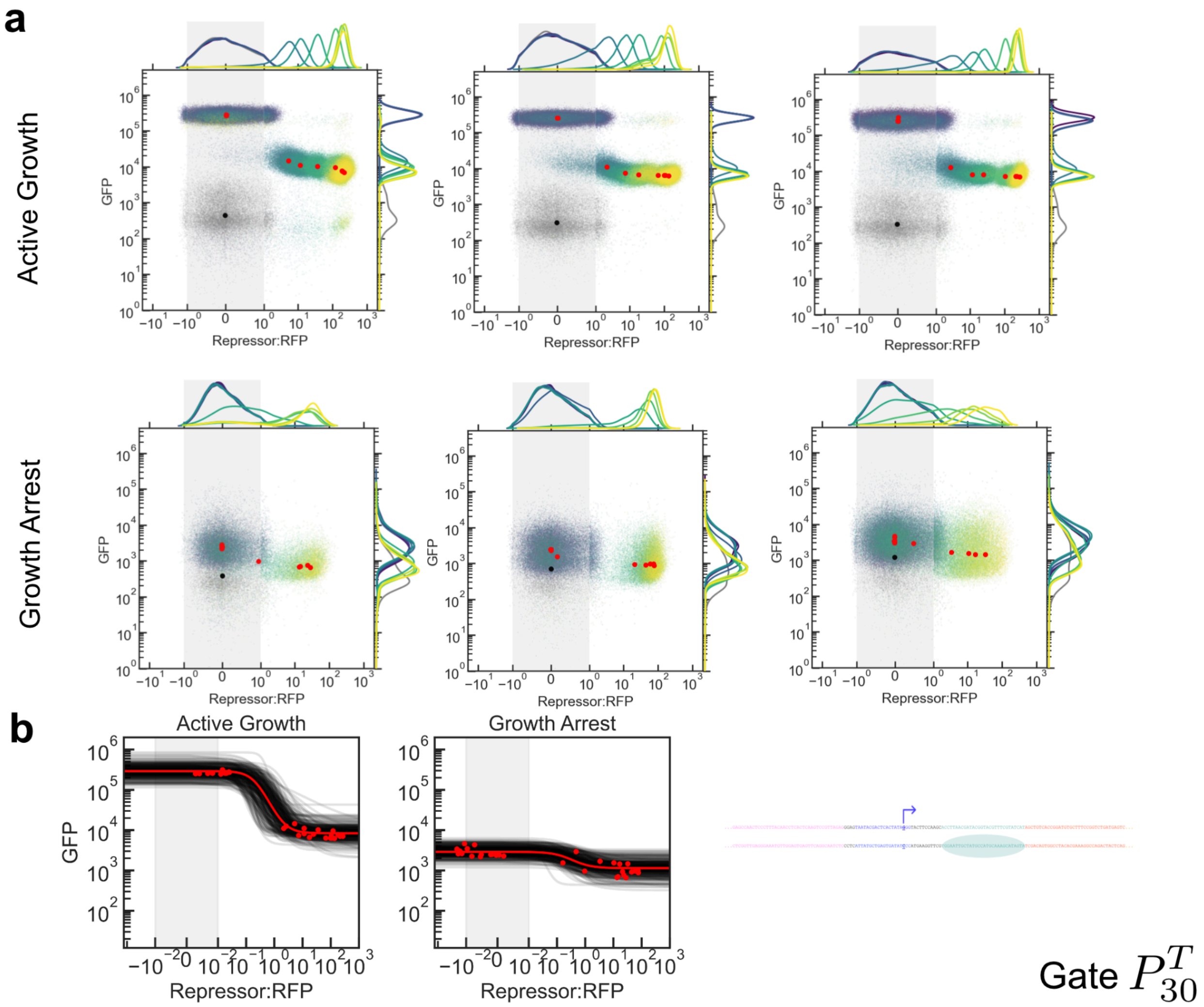


***Supplemental Fig. 21:*** *Repression curves for gate $P^T_{30}$. (**a**) Flow cytometry data. Three biological replicates (columns) were measured on three different days. Dots represent medians of distributions for each induction condition, and colors represent atc concentrations. Induction conditions for atc were: Active Growth, all replicates: 0, 0.2, 0.43, 0.93, 2, 2.6, 3.5, 6.3, 11.3, 20, 100 ng/mL. Growth Arrest, all replicates: 0, 0.02, 0.06, 0.2, 0.43, 0.93, 2, 6, 20, 60, 200 ng/mL. Grey points indicate cultures that were not induced with either IPTG or atc. Data are plotted on a symmetric log scale where the grey shaded region is linear. (**b**) 300 trajectories taken from the end of the MCMC process overlaid against the median (RFP,GFP) values from all three replicates.*

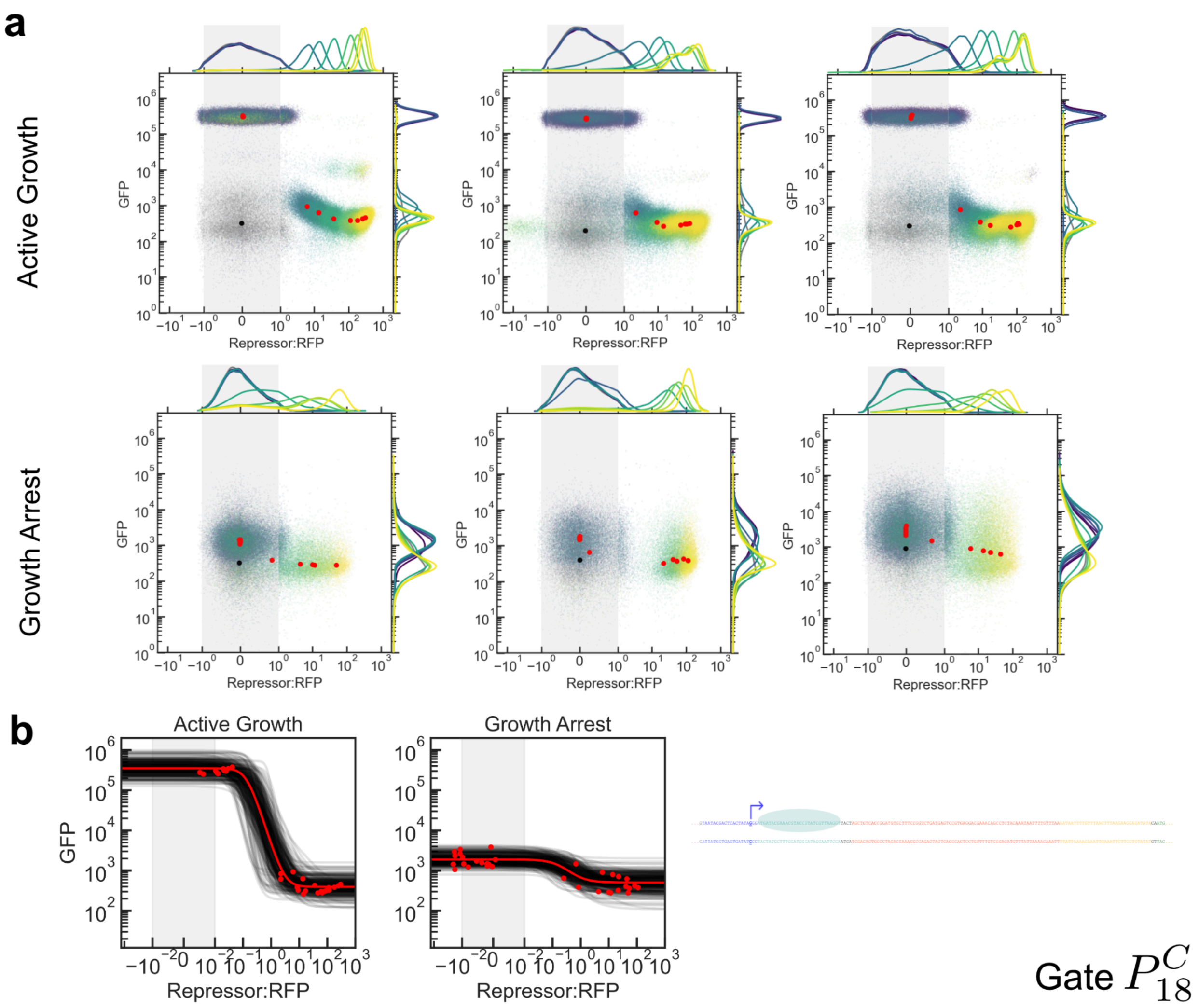


***Supplemental Fig. 22:*** *Repression curves for gate $P^{C}_{18}$. (**a**) Flow cytometry data. Three biological replicates (columns) were measured on three different days. Dots represent medians of distributions for each induction condition, and colors represent atc concentrations. Induction conditions for atc were: Active Growth, all replicates: 0, 0.2, 0.43, 0.93, 2, 2.6, 3.5, 6.3, 11.3, 20, 100 ng/mL. Growth Arrest, all replicates: 0, 0.02, 0.06, 0.2, 0.43, 0.93, 2, 6, 20, 60, 200 ng/mL. Grey points indicate cultures that were not induced with either IPTG or atc. Data are plotted on a symmetric log scale where the grey shaded region is linear. (**b**) 300 trajectories taken from the end of the MCMC process overlaid against the median (RFP,GFP) values from all three replicates.*

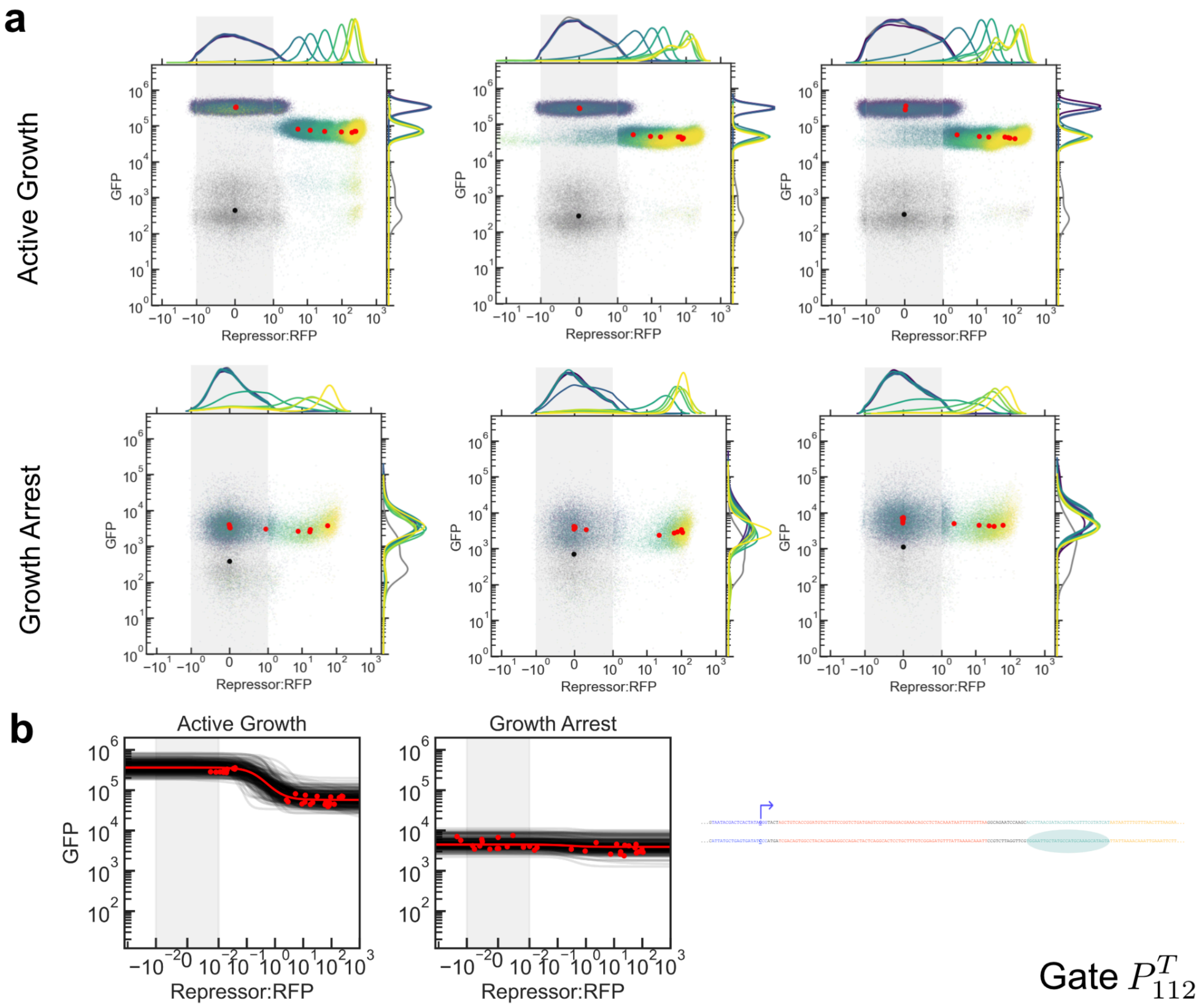


***Supplemental Fig. 23:*** *Repression curves for gate $P^T_{112}$. (**a**) Flow cytometry data. Three biological replicates (columns) were measured on three different days. Dots represent medians of distributions for each induction condition, and colors represent atc concentrations. Induction conditions for atc were: Active Growth, all replicates: 0, 0.2, 0.43, 0.93, 2, 2.6, 3.5, 6.3, 11.3, 20, 100 ng/mL. Growth Arrest, all replicates: 0, 0.02, 0.06, 0.2, 0.43, 0.93, 2, 6, 20, 60, 200 ng/mL. Grey points indicate cultures that were not induced with either IPTG or atc. Data are plotted on a symmetric log scale where the grey shaded region is linear. (**b**) 300 trajectories taken from the end of the MCMC process overlaid against the median (RFP,GFP) values from all three replicates.*

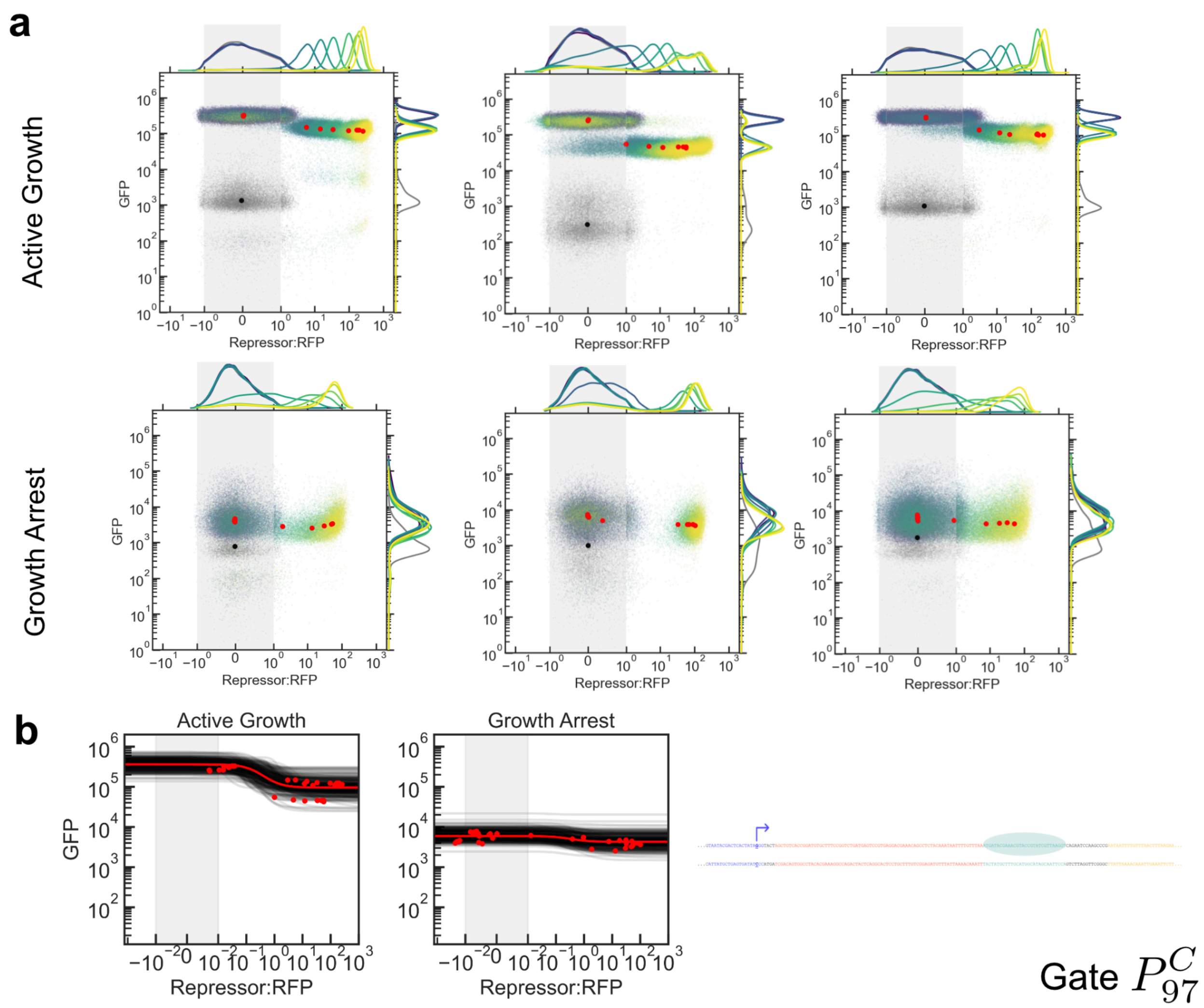


***Supplemental Fig. 24:*** *Repression curves for gate P$^{C}$$_{97}$. (**a**) Flow cytometry data. Three biological replicates (columns) were measured on three different days. Dots represent medians of distributions for each induction condition, and colors represent atc concentrations. Induction conditions for atc were: Active Growth, all replicates: 0, 0.2, 0.43, 0.93, 2, 2.6, 3.5, 6.3, 11.3, 20, 100 ng/mL. Growth Arrest, all replicates: 0, 0.02, 0.06, 0.2, 0.43, 0.93, 2, 6, 20, 60, 200 ng/mL. Grey points indicate cultures that were not induced with either IPTG or atc. Data are plotted on a symmetric log scale where the grey shaded region is linear. (**b**) 300 trajectories taken from the end of the MCMC process overlaid against the median (RFP,GFP) values from all three replicates.*

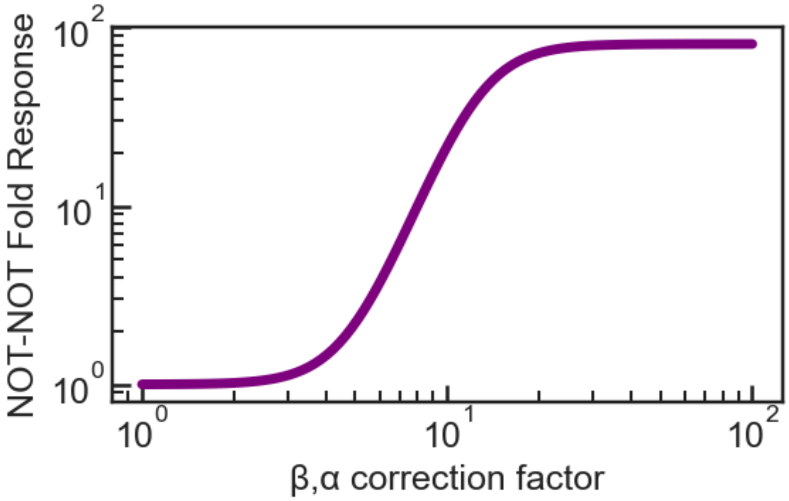


***Supplemental Fig. 25:*** *Predicted impact of mitigating the growth arrest-associated reduction in $\beta$, $\alpha$ on circuit peformance under growth arrest for the NOT-NOT circuit in Fig. 6. Fold response is calculated as the maximal possible OFF to ON fold change achievable by the NOT-NOT circuit, which is $H_2(\alpha_1)/H_2(\beta_1+\alpha_1)$ where $H_2(x)$ is the Hill function of the second repression curve (PhlF) and $\beta_1$, $\alpha_1$ are parameters from the first repression curve (SrpR). The $\beta$, $\alpha$ correction factor is globally multiplied against the growth-arrested values of $\beta$ and $\alpha$ in Fig. 6, so a correction factor of 1 (i.e., no correction) leads to the circuit performance shown in Fig. 6.*

### Supplemental Data

| Cassette | Sequence |
| --- | --- |
| T7, GFP cassette (Expression cassette for all D gates)<br><br>UNS 1 - PLacO - B0033 - T7 RNAP - T12m - UNS 5 - PT7 - RiboJ - UTR1 - sfGFP - T7 Terminator - UNS X | CATTACTCGCATCCATTCTCAGGCTGTCTCGTCTCGTCTCGGAGAATTGTGAGCGGATAACAATTGACATTGTGAGCGGATAA<br>CAAGATACTGAGCACATCAGCAGGACGCACTGACCTACTAGAGTCACACAGGACTACTAAATGAACACGATTAACATCGCTAA<br>GAACGACTTCTCTGACATCGAACTGGCTGCTATCCCGTTCAACACTCTGGCTGACCATTACGGTGAGCGTTTAGCTCGCGAAC<br>AGTTGGCCCTTGAGCATGAGTCTTACGAGATGGGTGAAGCACGCTTCCGCAAGATGTTTGAGCGTCAACTTAAAGCTGGTGAG<br>GTTGCGGATAACGCTGCCGCCAAGCCTCTCATCACTACCCTACTCCCTAAGATGATTGCACGCATCAACGACTGGTTTGAGGA<br>AGTGAAAGCTAAGCGCGGCAAGCGCCCGACAGCCTTCCAGTTCCTGCAAGAAATCAAGCCGGAAGCCGTAGCGTACATCACCA<br>TTAAGACCACTCTGGCTTGCCTAACCAGTGCTGACAATACAACCGTTCAGGCTGTAGCAAGCGCAATCGGTCGGGCCATTGAG<br>GACGAGGCTCGCTTCGGTCGTATCCGTGACCTTGAAGCTAAGCACTTCAAGAAAAACGTTGAGGAACAACTCAACAAGCGCGT<br>AGGGCACGTCTACAAGAAAGCATTTATGCAAGTTGTCGAGGCTGACATGCTCTCTAAGGGTCTACTCGGTGGCGAGGCGTGGT<br>CTTCGTGGCATAAGGAAGACTCTATTCATGTAGGAGTACGCTGCATCGAGATGCTCATTGAGTCAACCGGAATGGTTAGCTTA<br>CACCGCCAAAATGCTGGCGTAGTAGGTCAAGACTCTGAGACTATCGAACTCGCACCTGAATACGCTGAGGCTATCGCAACCCG<br>TGCAGGTGCGCTGGCTGGCATCTCTCCGATGTTCCAACCTTGCGTAGTTCCTCCTAAGCCGTGGACTGGCATTACTGGTGGTG<br>GCTATTGGGCTAACGGTCGTCGTCCTCTGGCGCTGGTGCGTACTCACAGTAAGAAAGCACTGATGCGCTACGAAGACGTTTAC<br>ATGCCTGAGGTGTACAAAGCGATTAACATTGCGCAAAACACCGCATGGAAAATCAACAAGAAAGTCCTAGCGGTCGCCAACGT<br>AATCACCAAGTGGAAGCATTGTCCGGTCGAGGACATCCCTGCGATTGAGCGTGAAGAACTCCCGATGAAACCGGAAGACATCG<br>ACATGAATCCTGAGGCTCTCACCGCGTGGAAACGTGCTGCCGCTGCTGTGTACCGCAAGGACAAGGCTCGCAAGTCTCGCCGT<br>ATCAGCCTTGAGTTCATGCTTGAGCAAGCCAATAAGTTTGCTAACCATAAGGCCATCTGGTTCCCTTACAACATGGACTGGCG<br>CGGTCGTGTTTACGCTGTGTCAATGTTCAACCCGCAAGGTAACGATATGACCAAAGGACTGCTTACGCTGGCGAAAGGTAAAC<br>CAATCGGTAAGGAAGGTTACTACTGGCTGAAAATCCACGGTGCAAACTGTGCGGGTGTCGATAAGGTTCCGTTCCCTGAGCGC<br>ATCAAGTTCATTGAGGAAAACCACGAGAACATCATGGCTTGCGCTAAGTCTCCACTGGAGAACACTTGGTGGGCTGAGCAAGA<br>TTCTCCGTTCTGCTTCCTTGCGTTCTGCTTTGAGTACGCTGGGGTACAGCACCACGGCCTGAGCTATAACTGCTCCCTTCCGC<br>TGGCGTTTGACGGGTCTTGCTCTGGCATCCAGCACTTCTCCGCGATGCTCCGAGATGAGGTAGGTGGTCGCGCGGTTAACTTG<br>CTTCCTAGTGAAACCGTTCAGGACATCTACGGGATTGTTGCTAAGAAAGTCAACGAGATTCTACAAGCAGACGCAATCAATGG<br>GACCGATAACGAAGTAGTTACCGTGACCGATGAGAACACTGGTGAAATCTCTGAGAAAGTCAAGCTGGGCACTAAGGCACTGG<br>CTGGTCAATGGCTGGCTTACGGTGTTACTCGCAGTGTGACTAAGCGTTCAGTCATGACGCTGGCTTACGGGTCCAAAGAGTTC<br>GGCTTCCGTCAACAAGTGCTGGAAGATACCATTCAGCCAGCTATTGATTCCGGCAAGGGTCTGATGTTCACTCAGCCGAATCA<br>GGCTGCTGGATACATGGCTAAGCTGATTTGGGAATCTGTGAGCGTGACGGTGGTAGCTGCGGTTGAAGCAATGAACTGGCTTA<br>AGTCTGCTGCTAAGCTGCTGGCTGCTGAGGTCAAAGATAAGAAGACTGGAGAGATTCTTCGCAAGCGTTGCGCTGTGCATTGG<br>GTAACTCCTGATGGTTTCCCTGTGTGGCAGGAATACAAGAAGCCTATTCAGACGCGCTTGAACCTGATGTTCCTCGGTCAGTT<br>CCGCTTACAGCCTACCATTAACACCAACAAAGATAGCGAGATTGATGCACACAAACAGGAGTCTGGTATCGCTCCTAACTTTG<br>TACACAGCCAAGACGGTAGCCACCTTCGTAAGACTGTAGTGTGGGCACACGAGAAGTACGGAATCGAATCTTTTGCACTGATT<br>CACGACTCCTTCGGTACCATTCCGGCTGACGCTGCGAACCTGTTCAAAGCAGTGCGCGAAACTATGGTTGACACATATGAGTC<br>TTGTGATGTACTGGCTGATTTCTACGACCAGTTCGCTGACCAGTTGCACGAGTCTCAATTGGACAAAATGCCAGCACTTCCGG<br>CTAAAGGTAACTTGAACCTCCGTGACATCTTAGAGTCGGACTTCGCGTTCGCGTAAAGGTTTCAGCCAAAAAACTTAAGACCG<br>CCGGTCTTGTCCACTACCTTGCAGTAATGCGGTGGACAGGATCGGCGGTTTTCTTTTCTCTTCTCAAGCTTGAGCCAACTCCC<br>TTTACAACCTCACTCAAGTCCGTTAGAGGGAGTAATACGACTCACTATAGGGTACTAGCTGTCACCGGATGTGCTTTCCGGTC<br>TGATGAGTCCGTGAGGACGAAACAGCCTCTACAAATAATTTTGTTTAAAATAATTTTGTTTAACTTTAAGAAGGAGATATACA<br>ATGCGTAAAGGCGAAGAGCTGTTCACTGGTGTCGTCCCTATTCTGGTGGAACTGGATGGTGATGTCAACGGTCATAAGTTTTC<br>CGTGCGTGGCGAGGGTGAAGGTGACGCAACTAATGGTAAACTGACGCTGAAGTTCATCTGTACTACTGGTAAACTGCCGGTAC<br>CTTGGCCGACTCTGGTAACGACGCTGACTTATGGTGTTCAGTGCTTTGCTCGTTATCCGGACCATATGAAGCAGCATGACTTC<br>TTCAAGTCCGCCATGCCGGAAGGCTATGTGCAGGAACGCACGATTTCCTTTAAGGATGACGGCACGTACAAAACGCGTGCGGA<br>AGTGAAATTTGAAGGCGATACCCTGGTAAACCGCATTGAGCTGAAAGGCATTGACTTTAAAGAAGATGGCAATATCCTGGGCC<br>ATAAGCTGGAATACAATTTTAACAGCCACAATGTTTACATCACCGCCGATAAACAAAAAAATGGCATTAAAGCGAATTTTAAA<br>ATTCGCCACAACGTGGAGGATGGCAGCGTGCAGCTGGCTGATCACTACCAGCAAAACACTCCAATCGGTGATGGTCCTGTTCT<br>GCTGCCAGACAATCACTATCTGAGCACGCAAAGCGTTCTGTCTAAAGATCCGAACGAGAAACGCGATCATATGGTTCTGCTGG<br>AGTTCGTAACCGCAGCGGGCATCACGCATGGTATGGATGAACTGTACAAATGATGAAGGTGCTGCTGCCACCGCTGAGCAATA<br>ACTAGCATAACCCCTTGGGGCCTCTAAACGGGTCTTGAGGGGTTTTTTGCTGAAAGGAGGAACTATAGCTTCCAGGATACATA<br>GATTACCACAACTCCGAGCCCTTCCACC |
| dCas9, gRNA cassette (Repressor cassette for all D gates)<br><br>UNS 1 - PTet - B0034 - dCas9:GGGSx2 Linker:mScarlet3 - T13m - UNS 6 - POR1OR2 - Hammerhead Ribozyme - gRNA (with variable sequence) - T14m - UNS X | CATTACTCGCATCCATTCTCAGGCTGTCTCGTCTCGTCTCGGAGTCCCTATCAGTGATAGAGATTGACATCCCTATCAGTGAT<br>AGAGATACTGAGCACTACTAGAGAAAGAGGAGAAATACTAAATGGATAAGAAATACTCAATAGGCTTAGCTATCGGCACAAAT<br>AGCGTCGGATGGGCGGTGATCACTGATGAATATAAGGTTCCGTCTAAAAAGTTCAAGGTTCTGGGAAATACAGACCGCCACAG<br>TATCAAAAAAAATCTTATAGGGGCTCTTTTATTTGACAGTGGAGAGACAGCGGAAGCGACTCGTCTCAAACGGACAGCTCGTA<br>GAAGGTATACACGTCGGAAGAATCGTATTTGTTATCTACAGGAGATTTTTTCAAATGAGATGGCGAAAGTAGATGATAGTTTC<br>TTTCATCGACTTGAAGAGTCTTTTTTGGTGGAAGAAGACAAGAAGCATGAACGTCATCCTATTTTTGGAAATATAGTAGATGA<br>AGTTGCTTATCATGAGAAATATCCAACTATCTATCATCTGCGAAAAAAATTGGTAGATTCTACTGATAAAGCGGATTTGCGCT<br>TAATCTATTTGGCCTTAGCGCATATGATTAAGTTTCGTGGTCATTTTTTGATTGAGGGAGATTTAAATCCTGATAATAGTGAT<br>GTGGACAAACTATTTATCCAGTTGGTACAAACCTACAATCAATTATTTGAAGAAAACCCTATTAACGCAAGTGGAGTAGATGC<br>TAAAGCGATTCTTTCTGCACGATTGAGTAAATCAAGACGATTAGAAAATCTCATTGCTCAGCTCCCCGGTGAGAAGAAAAATG<br>GCTTATTTGGGAATCTCATTGCTTTGTCATTGGGTTTGACCCCTAATTTTAAATCAAATTTTGATTTGGCAGAAGATGCTAAA<br>TTACAGCTTTCAAAAGATACTTACGATGATGATTTAGATAATTTATTGGCGCAAATTGGAGATCAATATGCTGATTTGTTTTT<br>GGCAGCTAAGAATTTATCAGATGCTATTTTACTTTCAGATATCCTAAGAGTAAATACTGAAATAACTAAGGCTCCCCTATCAG<br>CTTCAATGATTAAACGCTACGATGAACATCATCAAGACTTGACTCTTTTAAAAGCTTTAGTTCGACAACAACTTCCAGAAAAG<br>TATAAAGAAATCTTTTTTGATCAATCAAAAAACGGATATGCAGGTTATATTGATGGGGGAGCTAGCCAAGAAGAATTTTATAA<br>ATTTATCAAACCAATTTTAGAAAAAATGGATGGTACTGAGGAATTATTGGTGAAACTAAATCGTGAAGATTTGCTGCGCAAGC<br>AACGGACCTTTGACAACGGCTCTATTCCCCATCAAATTCACTTGGGTGAGCTGCATGCTATTTTGAGAAGACAAGAAGACTTT<br>TATCCATTTTTAAAAGACAATCGTGAGAAGATTGAAAAAATCTTGACTTTTCGAATTCCTTATTATGTTGGTCCATTGGCGCG<br>TGGCAATAGTCGTTTTGCATGGATGACTCGGAAGTCTGAAGAAACAATTACCCCATGGAATTTTGAAGAAGTTGTCGATAAAG<br>GTGCTTCAGCTCAATCATTTATTGAACGCATGACAAACTTTGATAAAAATCTTCCAAATGAAAAAGTACTACCAAAACATAGT<br>TTGCTTTATGAGTATTTTACGGTTTATAACGAATTGACAAAGGTCAAATATGTTACTGAAGGAATGCGAAAACCAGCATTTCT<br>TTCAGGTGAACAGAAGAAAGCCATTGTTGATTTACTCTTCAAAACAAATCGAAAAGTAACCGTTAAGCAATTAAAAGAAGATT<br>ATTTCAAAAAAATAGAATGTTTTGATAGTGTTGAAATTTCAGGAGTTGAAGATAGATTTAATGCTTCATTAGGTACCTACCAT |

| | |
|---|---|
| | GATTTGCTAAAAATTATTAAAGATAAAGATTTTTTGGATAATGAAGAAAATGAAGATATCTTAGAGGATATTGTTTTAACATT<br>GACCTTATTTGAAGATAGGGAGATGATTGAGGAAAGACTTAAAACATATGCTCACCTCTTTGATGATAAGGTGATGAAACAGC<br>TTAAACGTCGCCGTTATACTGGTTGGGGACGTTTGTCTCGAAAATTGATTAATGGTATTAGGGATAAGCAATCTGGCAAAACA<br>ATATTAGATTTTTTGAAATCAGATGGTTTTGCCAATCGCAATTTTATGCAGCTGATCCATGATGATAGTTTGACATTTAAAGA<br>AGACATTCAAAAAGCACAAGTGTCTGGACAAGGCGATAGTTTACATGAACATATTGCAAATTTAGCTGGTAGCCCTGCTATTA<br>AAAAAGGTATTTTACAGACTGTAAAAGTTGTTGATGAATTGGTCAAAGTAATGGGGCGGCATAAGCCAGAAAATATCGTTATT<br>GAAATGGCACGTGAAAATCAGACAACTCAAAAGGGCCAGAAAAATTCGCGAGAGCGTATGAAACGAATCGAAGAAGGTATCAA<br>AGAATTAGGAAGTCAGATTCTTAAAGAGCATCCTGTTGAAAATACTCAATTGCAAAATGAAAAGCTCTATCTCTATTATCTCC<br>AAAATGGAAGAGACATGTATGTGGACCAAGAATTAGATATTAATCGTTTAAGTGATTATGATGTCGATGCCATTGTTCCACAA<br>AGTTTCCTTAAAGACGATTCAATAGACAATAAGGTCTTAACGCGTTCTGATAAAAATCGTGGTAAATCGGATAACGTTCCAAG<br>TGAAGAAGTAGTCAAAAAGATGAAAAACTATTGGAGACAACTTCTAAACGCCAAGTTAATCACTCAACGTAAGTTTGATAATT<br>TAACGAAAGCTGAACGTGGAGGTTTGAGTGAACTTGATAAAGCTGGTTTTATCAAACGCCAATTGGTTGAAACTCGCCAAATC<br>ACTAAGCATGTGGCACAAATTTTGGATAGTCGCATGAATACTAAATACGATGAAAATGATAAACTTATTCGAGAGGTTAAAGT<br>GATTACCTTAAAATCTAAATTAGTTTCTGACTTCCGAAAAGATTTCCAATTCTATAAAGTACGTGAGATTAACAATTACCATC<br>ATGCCCATGATGCGTATCTAAATGCCGTCGTTGGAACTGCTTTGATTAAGAAATATCCAAAACTTGAATCGGAGTTTGTCTAT<br>GGTGATTATAAAGTTTATGATGTTCGTAAAATGATTGCTAAGTCTGAGCAAGAAATAGGCAAAGCAACCGCAAAATATTTCTT<br>TTACTCTAATATCATGAACTTCTTCAAAACAGAAATTACACTTGCAAATGGAGAGATTCGCAAACGCCCTCTAATCGAAACTA<br>ATGGGGAAACTGGAGAAATTGTCTGGGATAAAGGGCGAGATTTTGCCACAGTGCGCAAAGTATTGTCCATGCCCCAAGTCAAT<br>ATTGTCAAGAAAACAGAAGTACAGACAGGCGGATTCTCCAAGGAGTCAATTTTACCAAAAAGAAATTCGGACAAGCTTATTGC<br>TCGTAAAAAAGACTGGGATCCAAAAAAATATGGTGGTTTTGATAGTCCAACGGTAGCTTATTCAGTCCTAGTGGTTGCTAAGG<br>TGGAAAAAGGGAAATCGAAGAAGTTAAAATCCGTTAAAGAGTTACTAGGGATCACAATTATGGAAAGAAGTTCCTTTGAAAAA<br>AATCCGATTGACTTTTTAGAAGCTAAAGGATATAAGGAAGTTAAAAAAGACTTAATCATTAAACTACCTAAATATAGTCTTTT<br>TGAGTTAGAAAACGGTCGTAAACGGATGCTGGCTAGTGCCGGAGAATTACAAAAAGGAAATGAGCTGGCTCTGCCAAGCAAAT<br>ATGTGAATTTTTTATATTTAGCTAGTCATTATGAAAAGTTGAAGGGTAGTCCAGAAGATAACGAACAAAAACAATTGTTTGTG<br>GAGCAGCATAAGCATTATTTAGATGAGATTATTGAGCAAATCAGTGAATTTTCTAAGCGTGTTATTTTAGCAGATGCCAATTT<br>AGATAAAGTTCTTAGTGCATATAACAAACATAGAGACAAACCAATACGTGAACAAGCAGAAAATATTATTCATTTATTTACGT<br>TGACGAATCTTGGAGCTCCCGCTGCTTTTAAATATTTTGATACAACAATTGATCGTAAACGATATACGTCTACAAAAGAAGTT<br>TTAGATGCCACTCTTATCCATCAATCCATCACTGGTCTTTATGAAACACGCATTGATTTGAGTCAGCTAGGAGGTGACGGTGG<br>TGGTAGCGGTGGTGGTTCAGACTCAACCGAAGCGGTAATCAAGGAGTTTATGCGCTTCAAGGTGCACATGGAGGGGTCAATGA<br>ATGGGCACGAGTTCGAGATCGAAGGAGAAGGTGAGGGTCGTCCATACGAAGGCACTCAGACCGCTAAGTTACGTGTCACCAAA<br>GGAGGCCCTTTGCCCTTCTCATGGGATATTTTATCACCACAATTTATGTACGGATCACGTGCATTTACAAAGCATCCGGCTGA<br>CATTCCTGATTATTGGAAGCAATCGTTCCCCGAGGGTTTAAATGGGAGCGTGTGATGAACTTTGAAGATGGGGGCGCCGTCT<br>CAGTTGCTCAGGACACCTCTCTGGAGGATGGTACCCTTATTTATAAGGTAAAGCTGCGTGGAACAAACTTCCCTCCTGACGGC<br>CCTGTCATGCAAAAAAAAACCATGGGGTGGGAAGCGAGTACAGAGCGTTTATATCCCGAGGATGTTGTACTGAAAGGGGACAT<br>CAAAATGGCTTTACGTTTGAAGGATGGAGGTCGCTACCTGGCGGACTTTAAGACAACATACCGCGCGAAAAAACCTGTCCAAA<br>TGCCCGGCGCTTTTAACATTGACCGTAAGTTGGACATCACAAGTCATAATGAAGATTACACGGTCGTTGAGCAGTACGAACGT<br>AGCGTCGCTCGCCATTCCACCGGCGGAAGTGGGGGTAGTTAAAGGTGGAAACACAGAAAAAAGCCCGCACCTGACAGTGCGGG<br>CTTTTTTTTTCGACCAAAGGGCTTCTCGTTCGCTGCCACCTAAGAATACTCTACGGTCACATACGGAGTGCTGTTCCGCTGGG<br>CATGCTGAGCTAACACCGTGCGTGTTGACAATTTTACCTCTGGCGGTGATAATGGTTGCAGCTACTAGTAGCCTGATGAGTCC<br>GTGAGGACGAAACGAGTAAGCTCGTCGCTACT**NNNNNNNNNNNNNNNNNNNN**GTTTAGAGCTAGAAATAGCAAGTTAAAAT<br>AAGGCTAGTCCGTTATCAACTTGAAAAAGTGGCACCGAGTCGGTGCTTTTTTTAGGTGACGAACAATAAGGCCTCCCTAACGG<br>GGGGCCTTTTTTATTGATAACAAAAGCTTCCAGGATACATAGATTACCACAACTCCGAGCCCTTCCACC |
| $D^{0}$ gRNA | AACAAACGCAGAATCCAAGC |
| $D^{T}_{-28}$ / $F^{T}_{-50}$ / $P^{T}_{-32}$ gRNA | CCTCACTCAAGTCCGTTAGA |
| $D^{C}_{-23}$ / $F^{C}_{-34}$ / $P^{C}_{-52}$ gRNA* | AGTCGTATTACTCCCTCTAA |
| $D^{T}_{-6}$ / $F^{T}_{12}$ / $P^{T}_{30}$ gRNA | GGAGTAATACGACTCACTAT |
| $D^{T}_{12}$ gRNA | TATAGGGTACTAGCTGTCAC |
| $D^{C}_{23}$ / $F^{C}_{36}$ / $P^{C}_{18}$ gRNA | ATCAGACCGGAAAGCACATC |
| $D^{T}_{25}$ gRNA | CTGTCACCGGATGTGCTTTC |
| $D^{T}_{43}$ gRNA | TCCGGTCTGATGAGTCCGTG |
| $D^{C}_{50}$ gRNA | TAGAGGCTGTTTCGTCCTCA |
| $D^{C}_{67}$ gRNA | TTAAACAAAATTATTTGTAG |
| $D^{T}_{101}$ gRNA | AATTTTGTTTAACTTTAAGA |
| $D^{T}_{156}$ gRNA | CACTGGTGTCGTCCCTATTC |
| $D^{C}_{160}$ gRNA | CATCCAGTTCCACCAGAATA |

| | |
|---|---|
| $F^{T}_{94}$ / $P^{T}_{112}$ gRNA | TCTACAAATAATTTTGTTTA |
| $F^{C}_{117}$ / $P^{C}_{97}$ gRNA | TAAAGTTAAACAAAATTATT |
| T7, GFP cassette for $F^{T}_{-50}$ and $P^{T}_{-32}$<br><br>UNS 1 - PLacO - B0033 - T7 RNAP - T12m - UNS 5 - spacer - PhlO (rev. comp.) - PT7 - RiboJ - UTR1 - sfGFP - T7 Terminator - UNS X | [Identical sequence to PLacO T7 RNAP expression cassette for D gates]GAGCCAACTCCCTTTACAACCTCACTCAAGTCCGTTAGAGGGAGGAATCCAAGCACCTTAACGATACGGTACGTT<br>TCGTATCATTAATACGACTCACTATAGGGTACTAGCTGTCACCGGATGTGCTTTCCGGTCTGATGAGTCCGTGAGGACGAAAC<br>AGCCTCTACAAATAATTTTGTTTAAAATAATTTTGTTTAACTTTAAGAAGGAGATATACAATGCGTAAAGGCGAAGAGCTGTT<br>CACTGGTGTCGTCCCTATTCTGGTGGAACTGGATGGTGATGTCAACGGTCATAAGTTTTCCGTGCGTGGCGAGGGTGAAGGTG<br>ACGCAACTAATGGTAAACTGACGCTGAAGTTCATCTGTACTACTGGTAAACTGCCGGTACCTTGGCCGACTCTGGTAACGACG<br>CTGACTTATGGTGTTCAGTGCTTTGCTCGTTATCCGGACCATATGAAGCAGCATGACTTCTTCAAGTCCGCCATGCCGGAAGG<br>CTATGTGCAGGAACGCACGATTTCCTTTAAGGATGACGGCACGTACAAAACGCGTGCGGAAGTGAAATTTGAAGGCGATACCC<br>TGGTAAACCGCATTGAGCTGAAAGGCATTGACTTTAAAGAAGATGGCAATATCCTGGGCCATAAGCTGGAATACAATTTTAAC<br>AGCCACAATGTTTACATCACCGCCGATAAACAAAAAAATGGCATTAAAGCGAATTTTAAAATTCGCCACAACGTGGAGGATGG<br>CAGCGTGCAGCTGGCTGATCACTACCAGCAAAACACTCCAATCGGTGATGGTCCTGTTCTGCTGCCAGACAATCACTATCTGA<br>GCACGCAAAGCGTTCTGTCTAAAGATCCGAACGAGAAACGCGATCATATGGTTCTGCTGGAGTTCGTAACCGCAGCGGGCATC<br>ACGCATGGTATGGATGAACTGTACAAATGATGAAGGTGCTGCTGCCACCGCTGAGCAATAACTAGCATAACCCCTTGGGGCCT<br>CTAAACGGGTCTTGAGGGGTTTTTTGCTGAAAGGAGGAACTATAGCTTCCAGGATACATAGATTACCACAACTCCGAGCCCTT<br>CCACC |
| T7, GFP cassette for $F^{C}_{-34}$ and $P^{C}_{-52}$<br><br>UNS 1 - PLacO - B0033 - T7 RNAP - T12m - UNS 5 - PhlO - - spacer - PT7 - RiboJ - UTR1 - sfGFP - T7 Terminator - UNS X | [Identical sequence to PLacO T7 RNAP expression cassette for D gates]GAGCCAACTCCCTTTACAACCTCACTCAAGTCCGTTAGAGATGATACGAAACGTACCGTATCGTTAAGGTCAGAA<br>TCCAAGCCCGTAATACGACTCACTATAGGGTACTAGCTGTCACCGGATGTGCTTTCCGGTCTGATGAGTCCGTGAGGACGAAA<br>CAGCCTCTACAAATAATTTTGTTTAAAATAATTTTGTTTAACTTTAAGAAGGAGATATACAATGCGTAAAGGCGAAGAGCTGT<br>TCACTGGTGTCGTCCCTATTCTGGTGGAACTGGATGGTGATGTCAACGGTCATAAGTTTTCCGTGCGTGGCGAGGGTGAAGGT<br>GACGCAACTAATGGTAAACTGACGCTGAAGTTCATCTGTACTACTGGTAAACTGCCGGTACCTTGGCCGACTCTGGTAACGAC<br>GCTGACTTATGGTGTTCAGTGCTTTGCTCGTTATCCGGACCATATGAAGCAGCATGACTTCTTCAAGTCCGCCATGCCGGAAG<br>GCTATGTGCAGGAACGCACGATTTCCTTTAAGGATGACGGCACGTACAAAACGCGTGCGGAAGTGAAATTTGAAGGCGATACC<br>CTGGTAAACCGCATTGAGCTGAAAGGCATTGACTTTAAAGAAGATGGCAATATCCTGGGCCATAAGCTGGAATACAATTTTAA<br>CAGCCACAATGTTTACATCACCGCCGATAAACAAAAAAATGGCATTAAAGCGAATTTTAAAATTCGCCACAACGTGGAGGATG<br>GCAGCGTGCAGCTGGCTGATCACTACCAGCAAAACACTCCAATCGGTGATGGTCCTGTTCTGCTGCCAGACAATCACTATCTG<br>AGCACGCAAAGCGTTCTGTCTAAAGATCCGAACGAGAAACGCGATCATATGGTTCTGCTGGAGTTCGTAACCGCAGCGGGCAT<br>CACGCATGGTATGGATGAACTGTACAAATGATGAAGGTGCTGCTGCCACCGCTGAGCAATAACTAGCATAACCCCTTGGGGCC<br>TCTAAACGGGTCTTGAGGGGTTTTTTGCTGAAAGGAGGAACTATAGCTTCCAGGATACATAGATTACCACAACTCCGAGCCCT<br>TCCACC |
| T7, GFP cassette for $F^{T}_{12}$ and $P^{T}_{30}$<br><br>UNS 1 - PLacO - B0033 - T7 RNAP - T12m - UNS 5 - PT7 - spacer - PhlO (rev. comp.) - RiboJ - UTR1 - sfGFP - T7 Terminator - UNS X | [Identical sequence to PLacO T7 RNAP expression cassette for D gates]GAGCCAACTCCCTTTACAACCTCACTCAAGTCCGTTAGAGGGAGTAATACGACTCACTATAGGGTACTTCCAAGC<br>ACCTTAACGATACGGTACGTTTCGTATCATAGCTGTCACCGGATGTGCTTTCCGGTCTGATGAGTCCGTGAGGACGAAACAGC<br>CTCTACAAATAATTTTGTTTAAAATAATTTTGTTTAACTTTAAGAAGGAGATATACAATGCGTAAAGGCGAAGAGCTGTTCAC<br>TGGTGTCGTCCCTATTCTGGTGGAACTGGATGGTGATGTCAACGGTCATAAGTTTTCCGTGCGTGGCGAGGGTGAAGGTGACG<br>CAACTAATGGTAAACTGACGCTGAAGTTCATCTGTACTACTGGTAAACTGCCGGTACCTTGGCCGACTCTGGTAACGACGCTG<br>ACTTATGGTGTTCAGTGCTTTGCTCGTTATCCGGACCATATGAAGCAGCATGACTTCTTCAAGTCCGCCATGCCGGAAGGCTA<br>TGTGCAGGAACGCACGATTTCCTTTAAGGATGACGGCACGTACAAAACGCGTGCGGAAGTGAAATTTGAAGGCGATACCCTGG<br>TAAACCGCATTGAGCTGAAAGGCATTGACTTTAAAGAAGATGGCAATATCCTGGGCCATAAGCTGGAATACAATTTTAACAGC<br>CACAATGTTTACATCACCGCCGATAAACAAAAAAATGGCATTAAAGCGAATTTTAAAATTCGCCACAACGTGGAGGATGGCAG<br>CGTGCAGCTGGCTGATCACTACCAGCAAAACACTCCAATCGGTGATGGTCCTGTTCTGCTGCCAGACAATCACTATCTGAGCA<br>CGCAAAGCGTTCTGTCTAAAGATCCGAACGAGAAACGCGATCATATGGTTCTGCTGGAGTTCGTAACCGCAGCGGGCATCACG<br>CATGGTATGGATGAACTGTACAAATGATGAAGGTGCTGCTGCCACCGCTGAGCAATAACTAGCATAACCCCTTGGGGCCTCTA<br>AACGGGTCTTGAGGGGTTTTTTGCTGAAAGGAGGAACTATAGCTTCCAGGATACATAGATTACCACAACTCCGAGCCCTTCCA<br>CC |
| T7, GFP cassette for $F^{C}_{36}$ and $P^{C}_{18}$<br><br>UNS 1 - PLacO - B0033 - T7 RNAP - T12m - UNS 5 - PT7 - PhlO - RiboJ - UTR1 - sfGFP - T7 Terminator - UNS X | [Identical sequence to PLacO T7 RNAP expression cassette for D gates]GAGCCAACTCCCTTTACAACCTCACTCAAGTCCGTTAGAGGGAGTAATACGACTCACTATAGGGATGATACGAAA<br>CGTACCGTATCGTTAAGGTTACTAGCTGTCACCGGATGTGCTTTCCGGTCTGATGAGTCCGTGAGGACGAAACAGCCTCTACA<br>AATAATTTTGTTTAAAATAATTTTGTTTAACTTTAAGAAGGAGATATACAATGCGTAAAGGCGAAGAGCTGTTCACTGGTGTC<br>GTCCCTATTCTGGTGGAACTGGATGGTGATGTCAACGGTCATAAGTTTTCCGTGCGTGGCGAGGGTGAAGGTGACGCAACTAA<br>TGGTAAACTGACGCTGAAGTTCATCTGTACTACTGGTAAACTGCCGGTACCTTGGCCGACTCTGGTAACGACGCTGACTTATG<br>GTGTTCAGTGCTTTGCTCGTTATCCGGACCATATGAAGCAGCATGACTTCTTCAAGTCCGCCATGCCGGAAGGCTATGTGCAG<br>GAACGCACGATTTCCTTTAAGGATGACGGCACGTACAAAACGCGTGCGGAAGTGAAATTTGAAGGCGATACCCTGGTAAACCG<br>CATTGAGCTGAAAGGCATTGACTTTAAAGAAGATGGCAATATCCTGGGCCATAAGCTGGAATACAATTTTAACAGCCACAATG<br>TTTACATCACCGCCGATAAACAAAAAAATGGCATTAAAGCGAATTTTAAAATTCGCCACAACGTGGAGGATGGCAGCGTGCAG<br>CTGGCTGATCACTACCAGCAAAACACTCCAATCGGTGATGGTCCTGTTCTGCTGCCAGACAATCACTATCTGAGCACGCAAAG<br>CGTTCTGTCTAAAGATCCGAACGAGAAACGCGATCATATGGTTCTGCTGGAGTTCGTAACCGCAGCGGGCATCACGCATGGTA<br>TGGATGAACTGTACAAATGATGAAGGTGCTGCTGCCACCGCTGAGCAATAACTAGCATAACCCCTTGGGGCCTCTAAACGGGT<br>CTTGAGGGGTTTTTTGCTGAAAGGAGGAACTATAGCTTCCAGGATACATAGATTACCACAACTCCGAGCCCTTCCACC |
| T7, GFP cassette for $F^{T}_{94}$ and $P^{T}_{112}$ | [Identical sequence to PLacO T7 RNAP expression cassette for D gates]GAGCCAACTCCCTTTACAACCTCACTCAAGTCCGTTAGAGGGAGTAATACGACTCACTATAGGGTACTAGCTGTC |

| | |
|---|---|
| UNS 1 - PLacO - B0033 - T7 RNAP - T12m - UNS 5 - PT7 - RiboJ - spacer - PhlO (rev. comp.) - UTR1 - sfGFP - T7 Terminator - UNS X | ACCGGATGTGCTTTCCGGTCTGATGAGTCCGTGAGGACGAAACAGCCTCTACAAATAATTTTGTTTAAGGCAGAATCCAAGCA<br>CCTTAACGATACGGTACGTTTCGTATCATAATAATTTTGTTTAACTTTAAGAAGGAGATATACAATGCGTAAAGGCGAAGAGC<br>TGTTCACTGGTGTCGTCCCTATTCTGGTGGAACTGGATGGTGATGTCAACGGTCATAAGTTTTCCGTGCGTGGCGAGGGTGAA<br>GGTGACGCAACTAATGGTAAACTGACGCTGAAGTTCATCTGTACTACTGGTAAACTGCCGGTACCTTGGCCGACTCTGGTAAC<br>GACGCTGACTTATGGTGTTCAGTGCTTTGCTCGTTATCCGGACCATATGAAGCAGCATGACTTCTTCAAGTCCGCCATGCCGG<br>AAGGCTATGTGCAGGAACGCACGATTTCCTTTAAGGATGACGGCACGTACAAAACGCGTGCGGAAGTGAAATTTGAAGGCGAT<br>ACCCTGGTAAACCGCATTGAGCTGAAAGGCATTGACTTTAAAGAAGATGGCAATATCCTGGGCCATAAGCTGGAATACAATTT<br>TAACAGCCACAATGTTTACATCACCGCCGATAAACAAAAAAATGGCATTAAAGCGAATTTTAAAATTCGCCACAACGTGGAGG<br>ATGGCAGCGTGCAGCTGGCTGATCACTACCAGCAAAACACTCCAATCGGTGATGGTCCTGTTCTGCTGCCAGACAATCACTAT<br>CTGAGCACGCAAAGCGTTCTGTCTAAAGATCCGAACGAGAAACGCGATCATATGGTTCTGCTGGAGTTCGTAACCGCAGCGGG<br>CATCACGCATGGTATGGATGAACTGTACAAATGATGAAGGTGCTGCTGCCACCGCTGAGCAATAACTAGCATAACCCCTTGGG<br>GCCTCTAAACGGGTCTTGAGGGGTTTTTTGCTGAAAGGAGGAACTATAGCTTCCAGGATACATAGATTACCACAACTCCGAGC<br>CCTTCCACC |
| T7, GFP cassette for $F^{C}_{115}$ and $P^{C}_{97}$<br><br>UNS 1 - PLacO - B0033 - T7 RNAP - T12m - UNS 5 - PT7 - RiboJ - PhlO - spacer - UTR1 - sfGFP - T7 Terminator - UNS X | [Identical sequence to PLacO T7 RNAP expression cassette for D gates]GAGCCAACTCCCTTTACAACCTCACTCAAGTCCGTTAGAGGGAGTAATACGACTCACTATAGGGTACTAGCTGTC<br>ACCGGATGTGCTTTCCGGTCTGATGAGTCCGTGAGGACGAAACAGCCTCTACAAATAATTTTGTTTAAATGATACGAAACGTA<br>CCGTATCGTTAAGGTCAGAATCCAAGCCCGAATAATTTTGTTTAACTTTAAGAAGGAGATATACAATGCGTAAAGGCGAAGAG<br>CTGTTCACTGGTGTCGTCCCTATTCTGGTGGAACTGGATGGTGATGTCAACGGTCATAAGTTTTCCGTGCGTGGCGAGGGTGA<br>AGGTGACGCAACTAATGGTAAACTGACGCTGAAGTTCATCTGTACTACTGGTAAACTGCCGGTACCTTGGCCGACTCTGGTAA<br>CGACGCTGACTTATGGTGTTCAGTGCTTTGCTCGTTATCCGGACCATATGAAGCAGCATGACTTCTTCAAGTCCGCCATGCCG<br>GAAGGCTATGTGCAGGAACGCACGATTTCCTTTAAGGATGACGGCACGTACAAAACGCGTGCGGAAGTGAAATTTGAAGGCGA<br>TACCCTGGTAAACCGCATTGAGCTGAAAGGCATTGACTTTAAAGAAGATGGCAATATCCTGGGCCATAAGCTGGAATACAATT<br>TTAACAGCCACAATGTTTACATCACCGCCGATAAACAAAAAAATGGCATTAAAGCGAATTTTAAAATTCGCCACAACGTGGAG<br>GATGGCAGCGTGCAGCTGGCTGATCACTACCAGCAAAACACTCCAATCGGTGATGGTCCTGTTCTGCTGCCAGACAATCACTA<br>TCTGAGCACGCAAAGCGTTCTGTCTAAAGATCCGAACGAGAAACGCGATCATATGGTTCTGCTGGAGTTCGTAACCGCAGCGG<br>GCATCACGCATGGTATGGATGAACTGTACAAATGATGAAGGTGCTGCTGCCACCGCTGAGCAATAACTAGCATAACCCCTTGG<br>GGCCTCTAAACGGGTCTTGAGGGGTTTTTTGCTGAAAGGAGGAACTATAGCTTCCAGGATACATAGATTACCACAACTCCGAG<br>CCCTTCCACC |
| dCas9*:PhlF, gRNA cassette for all F gates<br><br>UNS 1 - PTet - B0034 - dCas9*:linker:PhlF:GGGSx2 Linker:mScarlet3 - T13m - UNS 6 - POR1OR2 - Hammerhead Ribozyme - gRNA (with variable sequence) - T14m - UNS X | CATTACTCGCATCCATTCTCAGGCTGTCTCGTCTCGTCTCGGAGTCCCTATCAGTGATAGAGATTGACATCCCTATCAGTGAT<br>AGAGATACTGAGCACTACTAGAGAAAGAGGAGAAATACTAAATGGATAAGAAATACTCAATAGGCTTAGCTATCGGCACAAAT<br>AGCGTCGGATGGGCGGTGATCACTGATGAATATAAGGTTCCGTCTAAAAAGTTCAAGGTTCTGGGAAATACAGACCGCCACAG<br>TATCAAAAAAAATCTTATAGGGGCTCTTTTATTTGACAGTGGAGAGACAGCGGAAGCGACTCGTCTCAAACGGACAGCTCGTA<br>GAAGGTATACACGTCGGAAGAATCGTATTTGTTATCTACAGGAGATTTTTTCAAATGAGATGGCGAAAGTAGATGATAGTTTC<br>TTTCATCGACTTGAAGAGTCTTTTTTGGTGGAAGAAGACAAGAAGCATGAACGTCATCCTATTTTTGGAAATATAGTAGATGA<br>AGTTGCTTATCATGAGAAATATCCAACTATCTATCATCTGCGAAAAAAATTGGTAGATTCTACTGATAAAGCGGATTTGCGCT<br>TAATCTATTTGGCCTTAGCGCATATGATTAAGTTTCGTGGTCATTTTTTGATTGAGGGAGATTTAAATCCTGATAATAGTGAT<br>GTGGACAAACTATTTATCCAGTTGGTACAAACCTACAATCAATTATTTGAAGAAAACCCTATTAACGCAAGTGGAGTAGATGC<br>TAAAGCGATTCTTTCTGCACGATTGAGTAAATCAAGACGATTAGAAAATCTCATTGCTCAGCTCCCCGGTGAGAAGAAAAATG<br>GCTTATTTGGGAATCTCATTGCTTTGTCATTGGGTTTGACCCCTAATTTTAAATCAAATTTTGATTTGGCAGAAGATGCTAAA<br>TTACAGCTTTCAAAAGATACTTACGATGATGATTTAGATAATTTATTGGCGCAAATTGGAGATCAATATGCTGATTTGTTTTT<br>GGCAGCTAAGAATTTATCAGATGCTATTTTACTTTCAGATATCCTAAGAGTAAATACTGAAATAACTAAGGCTCCCCTATCAG<br>CTTCAATGATTAAACGCTACGATGAACATCATCAAGACTTGACTCTTTTAAAAGCTTTAGTTCGACAACAACTTCCAGAAAAG<br>TATAAAGAAATCTTTTTTGATCAATCAAAAAACGGATATGCAGGTTATATTGATGGGGGAGCTAGCCAAGAAGAATTTTATAA<br>ATTTATCAAACCAATTTTAGAAAAAATGGATGGTACTGAGGAATTATTGGTGAAACTAAATCGTGAAGATTTGCTGCGCAAGC<br>AACGGACCTTTGACAACGGCTCTATTCCCCATCAAATTCACTTGGGTGAGCTGCATGCTATTTTGAGAAGACAAGAAGACTTT<br>TATCCATTTTTAAAAGACAATCGTGAGAAGATTGAAAAAATCTTGACTTTTCGAATTCCTTATTATGTTGGTCCATTGGCGCG<br>TGGCAATAGTCGTTTTGCATGGATGACTCGGAAGTCTGAAGAAACAATTACCCCATGGAATTTTGAAGAAGTTGTCGATAAAG<br>GTGCTTCAGCTCAATCATTTATTGAACGCATGACAAACTTTGATAAAAATCTTCCAAATGAAAAAGTACTACCAAAACATAGT<br>TTGCTTTATGAGTATTTTACGGTTTATAACGAATTGACAAAGGTCAAATATGTTACTGAAGGAATGCGAAAACCAGCATTTCT<br>TTCAGGTGAACAGAAGAAAGCCATTGTTGATTTACTCTTCAAAACAAATCGAAAAGTAACCGTTAAGCAATTAAAAGAAGATT<br>ATTTCAAAAAAATAGAATGTTTTGATAGTGTTGAAATTTCAGGAGTTGAAGATAGATTTAATGCTTCATTAGGTACCTACCAT<br>GATTTGCTAAAAATTATTAAAGATAAAGATTTTTTGGATAATGAAGAAAATGAAGATATCTTAGAGGATATTGTTTTAACATT<br>GACCTTATTTGAAGATAGGGAGATGATTGAGGAAAGACTTAAAACATATGCTCACCTCTTTGATGATAAGGTGATGAAACAGC<br>TTAAACGTCGCCGTTATACTGGTTGGGGACGTTTGTCTCGAAAATTGATTAATGGTATTAGGGATAAGCAATCTGGCAAAACA<br>ATATTAGATTTTTTGAAATCAGATGGTTTTGCCAATCGCAATTTTATGCAGCTGATCCATGATGATAGTTTGACATTTAAAGA<br>AGACATTCAAAAAGCACAAGTGTCTGGACAAGGCGATAGTTTACATGAACATATTGCAAATTTAGCTGGTAGCCCTGCTATTA<br>AAAAAGGTATTTTACAGACTGTAAAAGTTGTTGATGAATTGGTCAAAGTAATGGGGCGGCATAAGCCAGAAAATATCGTTATT<br>GAAATGGCACGTGAAAATCAGACAACTCAAAAGGGCCAGAAAAATTCGCGAGAGCGTATGAAACGAATCGAAGAAGGTATCAA<br>AGAATTAGGAAGTCAGATTCTTAAAGAGCATCCTGTTGAAAATACTCAATTGCAAAATGAAAAGCTCTATCTCTATTATCTCC<br>AAAATGGAAGAGACATGTATGTGGACCAAGAATTAGATATTAATCGTTTAAGTGATTATGATGTCGATGCCATTGTTCCACAA<br>AGTTTCCTTAAAGACGATTCAATAGACAATAAGGTCTTAACGCGTTCTGATAAAAATCGTGGTAAATCGGATAACGTTCCAAG<br>TGAAGAAGTAGTCAAAAAGATGAAAAACTATTGGAGACAACTTCTAAACGCCAAGTTAATCACTCAACGTAAGTTTGATAATT<br>TAACGAAAGCTGAACGTGGAGGTTTGAGTGAACTTGATAAAGCTGGTTTTATCAAACGCCAATTGGTTGAAACTCGCCAAATC<br>ACTAAGCATGTGGCACAAATTTTGGATAGTCGCATGAATACTAAATACGATGAAAATGATAAACTTATTCGAGAGGTTAAAGT<br>GATTACCTTAAAATCTAAATTAGTTTCTGACTTCCGAAAAGATTTCCAATTCTATAAAGTACGTGAGATTAACAATTACCATC<br>ATGCCCATGATGCGTATCTAAATGCCGTCGTTGGAACTGCTTTGATTAAGAAATATCCAAAACTTGAATCGGAGTTTGTCTAT<br>GGTGATTATAAAGTTTATGATGTTCGTAAAATGATTGCTAAGTCTGAGCAAGAAATAGGCAAAGCAACCGCAAAATATTTCTT<br>TTACTCTAATATCATGAACTTCTTCAAAACAGAAATTACACTTGCAAATGGAGAGATTCGCAAACGCCCTCTAATCGAAACTA<br>ATGGGGAAACTGGAGAAATTGTCTGGGATAAAGGGCGAGATTTTGCCACAGTGCGCAAAGTATTGTCCATGCCCCAAGTCAAT<br>ATTGTCAAGAAAACAGAAGTACAGACAGGCGGATTCTCCAAGGAGTCAATTTTACCAAAAAGAAATTCGGACAAGCTTATTGC<br>TCGTAAAAAAGACTGGGATCCAAAAAAATATGGTGGTTTTGATAGTCCAACGGTAGCTTATTCAGTCCTAGTGGTTGCTAAGG<br>TGGAAAAAGGGAAATCGAAGAAGTTAAAATCCGTTAAAGAGTTACTAGGGATCACAATTATGGAAAGAAGTTCCTTTGAAAAA<br>AATCCGATTGACTTTTTAGAAGCTAAAGGATATAAGGAAGTTAAAAAAGACTTAATCATTAAACTACCTAAATATAGTCTTTT<br>TGAGTTAGAAAACGGTCGTAAACGGATGCTGGCTAGTGCCGGAGAATTACAAAAAGGAAATGAGCTGGCTCTGCCAAGCAAAT |

| | |
|---|---|
| | ATGTGAATTTTTTATATTTAGCTAGTCATTATGAAAAGTTGAAGGGTAGTCCAGAAGATAACGAACAAAAACAATTGTTTGTG<br>GAGCAGCATAAGCATTATTTAGATGAGATTATTGAGCAAATCAGTGAATTTTCTAAGCGTGTTATTTTAGCAGATGCCAATTT<br>AGATAAAGTTCTTAGTGCATATAACAAACATAGAGACAAACCAATACGTGAACAAGCAGAAAATATTATTCATTTATTTACGT<br>TGACGAATCTTGGAGCTCCCGCTGCTTTTAAATATTTTGATACAACAATTGATCGTAAAAAGTATACGTCTACAAAAGAAGTT<br>TTAGATGCCACTCTTATCCATCAATCCATCACTGGTCTTTATGAAACACGCATTGATTTGAGTCAGCTAGGAGGTGACGGCAC<br>CGGCGGGCCCAAGAAGAAGAGGAAGGTATACCCATACGATGTTCCTGACTATGCGGGCTATCCCTATGACGTCCCGGACTATG<br>CAGGATCGTATCCTTATGACGTTCCAGATTACGCTGGATCCGCCGCTCCGGCAGCTAAGAAAAAGAAACTGGATTACCCGTAT<br>GACGTACCTGATTACGCTGGTTATCCCTATGATGTCCCGGACTACGCTGGCTCGTACCCTTATGATGTACCTGACTACGCTTT<br>CGAATCCGGAGCACGTACCCCGAGCCGTAGCAGCATTGGTAGCCTGCGTAGTCCGCATACCCATAAAGCAATTCTGACCAGCA<br>CCATTGAAATCCTGAAAGAATGTGGTTATAGCGGTCTGAGCATTGAAAGCGTTGCACGTCGTGCCGGTGCAAGCAAACCGACC<br>ATTTATCGTTGGTGGACCAATAAAGCAGCACTGATTGCCGAAGTGTATGAAAATGAAAGCGAACAGGTGCGTAAATTTCCGGA<br>TCTGGGTAGCTTTAAAGCCGATCTGGATTTTCTGCTGCGTAATCTGTGGAAAGTTTGGCGTGAAACCATTTGTGGTGAAGCAT<br>TTCGTTGTGTTATTGCAGAAGCACAGCTGGACCCTGCAACCCTGACCCAGCTGAAAGATCAGTTTATGGAACGTCGTCGTGAG<br>ATGCCGAAAAAACTGGTTGAAAATGCCATTAGCAATGGTGAACTGCCGAAAGATACCAATCGTGAACTGCTGCTGGATATGAT<br>TTTTGGTTTTTGTTGGTATCGCCTGCTGACCGAACAGCTGACCGTTGAACAGGATATTGAAGAATTTACCTTCCTGCTGATTA<br>ATGGTGTTTGTCCGGGTACACAGCGTGGTGGTGGTAGCGGTGGTGGTTCAGACTCAACCGAAGCGGTAATCAAGGAGTTTATG<br>CGCTTCAAGGTGCACATGGAGGGGTCAATGAATGGGCACGAGTTCGAGATCGAAGGAGAAGGTGAGGGTCGTCCATACGAAGG<br>CACTCAGACCGCTAAGTTACGTGTCACCAAAGGAGGCCCTTTGCCCTTCTCATGGGATATTTTATCACCACAATTTATGTACG<br>GATCACGTGCATTTACAAAGCATCCGGCTGACATTCCTGATTATTGGAAGCAATCGTTCCCCGAGGGTTTTAAATGGGAGCGT<br>GTGATGAACTTTGAAGATGGGGGCGCCGTCTCAGTTGCTCAGGACACCTCTCTGGAGGATGGTACCCTTATTTATAAGGTAAA<br>GCTGCGTGGAACAAACTTCCCTCCTGACGGCCCTGTCATGCAAAAAAAAACCATGGGGTGGGAAGCGAGTACAGAGCGTTTAT<br>ATCCCGAGGATGTTGTACTGAAAGGGGACATCAAAATGGCTTTACGTTTGAAGGATGGAGGTCGCTACCTGGCGGACTTTAAG<br>ACAACATACCGCGCGAAAAAACCTGTCCAAATGCCCGGCGCTTTTAACATTGACCGTAAGTTGGACATCACAAGTCATAATGA<br>AGATTACACGGTCGTTGAGCAGTACGAACGTAGCGTCGCTCGCCATTCCACCGGCGGAAGTGGGGGTAGTTAAAGGTGGAAAC<br>ACAGAAAAAAGCCCGCACCTGACAGTGCGGGCTTTTTTTTTCGACCAAAGGGCTTCTCGTTCGCTGCCACCTAAGAATACTCT<br>ACGGTCACATACGGAGTGCTGTTCCGCTGGGCATGCTGAGCTAACACCGTGCGTGTTGACAATTTTACCTCTGGCGGTGATAA<br>TGGTTGCAGCTACTAGTAGCCTGATGAGTCCGTGAGGACGAAACGAGTAAGCTCGTCGCTACTNNNNNNNNNNNNNNNNNNN<br>NGTTTTAGAGCTAGAAATAGCAAGTTAAAATAAGGCTAGTCCGTTATCAACTTGAAAAAGTGGCACCGAGTCGGTGCTTTTTT<br>TAGGTGACGAACAATAAGGCCTCCCTAACGGGGGGCCTTTTTTATTGATAACAAAAGCTTCCAGGATACATAGATTACCACAA<br>CTCCGAGCCCTTCCACC |
| PhlF, gRNA cassette for all P gates<br><br>UNS 1 - PTet - B0034 - PhlF:GGGSx2 Linker:mScarlet3 - T13m - UNS 6 - POR1OR2 - Hammerhead Ribozyme - gRNA (with variable sequence) - T14m - UNS X | CATTACTCGCATCCATTCTCAGGCTGTCTCGTCTCGTCTCGGAGTCCCTATCAGTGATAGAGATTGACATCCCTATCAGTGAT<br>AGAGATACTGAGCACTACTAGAGAAAGAGGAGAAATACTAAGCACGTACCCCGAGCCGTAGCAGCATTGGTAGCCTGCGTAGT<br>CCGCATACCCATAAAGCAATTCTGACCAGCACCATTGAAATCCTGAAAGAATGTGGTTATAGCGGTCTGAGCATTGAAAGCGT<br>TGCACGTCGTGCCGGTGCAAGCAAACCGACCATTTATCGTTGGTGGACCAATAAAGCAGCACTGATTGCCGAAGTGTATGAAA<br>ATGAAAGCGAACAGGTGCGTAAATTTCCGGATCTGGGTAGCTTTAAAGCCGATCTGGATTTTCTGCTGCGTAATCTGTGGAAA<br>GTTTGGCGTGAAACCATTTGTGGTGAAGCATTTCGTTGTGTTATTGCAGAAGCACAGCTGGACCCTGCAACCCTGACCCAGCT<br>GAAAGATCAGTTTATGGAACGTCGTCGTGAGATGCCGAAAAAACTGGTTGAAAATGCCATTAGCAATGGTGAACTGCCGAAAG<br>ATACCAATCGTGAACTGCTGCTGGATATGATTTTTGGTTTTTGTTGGTATCGCCTGCTGACCGAACAGCTGACCGTTGAACAG<br>GATATTGAAGAATTTACCTTCCTGCTGATTAATGGTGTTTGTCCGGGTACACAGCGTGGTGGTGGTAGCGGTGGTGGTTCAGA<br>CTCAACCGAAGCGGTAATCAAGGAGTTTATGCGCTTCAAGGTGCACATGGAGGGGTCAATGAATGGGCACGAGTTCGAGATCG<br>AAGGAGAAGGTGAGGGTCGTCCATACGAAGGCACTCAGACCGCTAAGTTACGTGTCACCAAAGGAGGCCCTTTGCCCTTCTCA<br>TGGGATATTTTATCACCACAATTTATGTACGGATCACGTGCATTTACAAAGCATCCGGCTGACATTCCTGATTATTGGAAGCA<br>ATCGTTCCCCGAGGGTTTTAAATGGGAGCGTGTGATGAACTTTGAAGATGGGGGCGCCGTCTCAGTTGCTCAGGACACCTCTC<br>TGGAGGATGGTACCCTTATTTATAAGGTAAAGCTGCGTGGAACAAACTTCCCTCCTGACGGCCCTGTCATGCAAAAAAAAACC<br>ATGGGGTGGGAAGCGAGTACAGAGCGTTTATATCCCGAGGATGTTGTACTGAAAGGGGACATCAAAATGGCTTTACGTTTGAA<br>GGATGGAGGTCGCTACCTGGCGGACTTTAAGACAACATACCGCGCGAAAAAACCTGTCCAAATGCCCGGCGCTTTTAACATTG<br>ACCGTAAGTTGGACATCACAAGTCATAATGAAGATTACACGGTCGTTGAGCAGTACGAACGTAGCGTCGCTCGCCATTCCACC<br>GGCGGAAGTGGGGGTAGTTAAAGGTGGAAACACAGAAAAAAGCCCGCACCTGACAGTGCGGGCTTTTTTTTTCGACCAAAGGG<br>CTTCTCGTTCGCTGCCACCTAAGAATACTCTACGGTCACATACGGAGTGCTGTTCCGCTGGGCATGCTGAGCTAACACCGTGC<br>GTGTTGACAATTTTACCTCTGGCGGTGATAATGGTTGCAGCTACTAGTAGCCTGATGAGTCCGTGAGGACGAAACGAGTAAGC<br>TCGTCGCTACTNNNNNNNNNNNNNNNNNNNNGTTTTAGAGCTAGAAATAGCAAGTTAAAATAAGGCTAGTCCGTTATCAACT<br>TGAAAAAGTGGCACCGAGTCGGTGCTTTTTTTAGGTGACGAACAATAAGGCCTCCCTAACGGGGGGCCTTTTTTATTGATAAC<br>AAAAGCTTCCAGGATACATAGATTACCACAACTCCGAGCCCTTCCACC |

* Note that this gRNA does not bind to any valid sequence in the $F^{C}_{-34}$ gate, but is labeled as such here because the gRNA was still expressed in the cells containing the $F^{C}_{-34}$ and $P^{C}_{-52}$ gates.

**Supplemental Table 1:** Sequences of all engineered circuit components.
